# UstiGate: A standardized modular cloning platform for advanced engineering of the fungal chassis *Ustilago maydis*

**DOI:** 10.64898/2026.06.11.731564

**Authors:** Joana C. Hasenklever, Valeria A. Paderi, Dennis Hasenklever, Ilka M. Axmann, Kerstin Schipper

## Abstract

The corn smut fungus *Ustilago maydis* is an established model for fungal biology and an emerging host for biotechnology. However, its broader application in synthetic biology has been constrained by a limited repertoire of standardized genetic parts and tools for multigene engineering. Here, we introduce UstiGate, a modular toolkit that combines MoClo-compatible DNA assembly with targeted genomic integration in *U. maydis*. The toolkit comprises 92 parts and vectors, including more than 20 promoter regions as well as terminators, selectable markers, fluorescent reporters and modules for different genomic integration sites. Quantitative reporter analyses at both, population and single-cell level, revealed a broad range of promoter strengths and characterized their behavior across growth phases, media, genomic loci and reporter contexts. We further identified a maltose-inducible, glucose-repressed promoter that enables autoinduction and established carotenoid-biosynthesis genes as integration sites supporting visual pre-selection of transformants. Finally, UstiGate enabled the assembly and simultaneous genomic integration of four transcriptional units in a single transformation. UstiGate thus provides a standardized and quantitatively characterized platform for standard applications as well as multigene engineering, enabling advanced synthetic biology applications such as pathway transplantation in *U. maydis*.

## 1. Introduction

While several ascomycetes are readily established as fungal production hosts, a suitable chassis closely related to mushroom-forming fungi from the phylum Basidiomycota, is still lacking ^1,2^. This is particularly relevant, as a close phylogenetic relationship may benefit the heterologous production of complex compounds, such as bioactive mushroom metabolites, by providing greater genetic and metabolic compatibility including the cellular machinery required for the proper function of complex biosynthetic enzymes ^1,3,4^.

With its yeast-like growth form, short generation time and well-established genetic engineering, the corn smut fungus *Ustilago maydis* provides a promising foundation for the development of a basidiomycete production host suitable for industrial application ^5^. Besides its long-standing role in fundamental biology topics such as plant-pathogen interaction, cell and RNA biology as well as signal transduction ^6–10^, applied research aspects have increasingly come into focus ^5,7^. *U. maydis* naturally produces several industrially relevant biomolecules including organic acids like itaconate or malate ^11^, glycolipids ^12–14^, triglycerides ^15,16^, and hydrolytic enzymes ^17–19^. Moreover, with regard to heterologous products, research has focused on the unconventional secretion of nanobodies ^20–23^ as well as on the synthesis of sesquiterpenes, representing the first heterologous secondary metabolites produced in *U. maydis* ^24^.

Genetic modifications in *U. maydis* are mostly conducted by homologous recombination using homologous flanks of about 1,000 bp ^25,26^. Unlike for example *Yarrowia lipolytica*, *U. maydis* supports efficient homologous recombination with rates up to 59 %, facilitating precise genome-editing ^26,27^. A first systematic vector set was implemented in 2004 ^28,29^ and adapted to Golden Gate ten years later ^26^, proving very effective for functional characterization of single genes. The vectors are available with five dominant selection markers, conferring resistance to hygromycin (HygR), nourseothricin (NatR), phleomycin (PhleoR), carboxin (CbxR) or geneticin (G418R) ^30–33^, and flippase (FLP)-mediated recombination allows for the generation of marker-free strains ^34^. Frequently used site-specific ectopic integration sites are the *ip* locus ^32^ and protease genes, such as *upp3* or *pep4*, where the deletion has no effect on cell morphology or growth of yeast-like *U. maydis* cells ^22,35^. Moreover, a small selection of frequently employed promoters is available, including native constitutive and inducible as well as synthetic promoters ^36–43^. Other relevant tools available for *U. maydis* include CRISPR/Cas9 methodology ^44–46^, tetracycline-controlled expression systems ^47,48^, fluorescent proteins (FPs), luciferases, epitope tags, degron sequences and a polycistronic gene expression system ^49,50^.

Although a practical, constantly evolving cloning system and a number of useful genetic tools are available for a wide range of applications, the genetic toolbox for *U. maydis* has proved to be limited when it comes to its application in synthetic biology. Wide-spanning metabolic engineering, for instance, or entire pathway transplantation require a system facilitating the generation of increasingly complex multigene expression constructs build from transcriptional units (TUs) including a suitable repertoire of promoters as well as efficient subsequent strain generation.

In this regard, the quick and flexible assembly of various genetic elements is essential for efficient “design-build-test-learn cycles” ^51^. The Golden Gate enables efficient assembly of complex constructs by exploiting the feature of type IIs restriction enzymes to hydrolyze DNA outside their recognition site, allowing scarless assembly of multiple DNA fragments in a one-pot restriction-ligation reaction ^52^. Modular cloning is a standardized hierarchal extension of Golden Gate cloning. Based on defined type IIs restriction enzyme overhangs, the so-called syntax, DNA parts can be excised from storage vectors and assembled in a predefined, modular manner into destination vectors. The hierarchical system progresses from libraries of assembly-ready individual genetic elements to basic modules and ultimately multigene constructs, providing high flexibility and streamlined construct generation ^53,54^.

The first two published strategies, the Modular Cloning (MoClo) ^55^ and the GoldenBraid ^56^ system, build upon specifically designed destination vectors for insertion of individual building blocks as well as for TUs, enabling straightforward assembly of both TUs and multigene constructs. In the plant synthetic biology field, these systems have been advanced to facilitate part exchangeability between the two Golden Gate syntaxes up to a common standard for exchange of DNA parts ^57–59^. Besides, GoldenBraid has been adapted and progressively expanded for filamentous fungi (FungalBraid) ^60–62^ and tailored for the social amoeba ^63^, while MoClo has provided the basis for numerous modular toolkits designed for microbial models like *Saccharomyces cerevisiae* ^64^, *Escherichia coli* ^65,66^, the microalga *Chlamydomonas reinhardtii* ^67^ and cyanobacteria (CyanoGate) ^68,69^.

Here, we developed UstiGate as a modular design-to-genome toolkit for synthetic biology applications in *U. maydis*. UstiGate retains compatibility with the established MoClo syntax ^55^, refined by the additional level of terminal acceptor vectors as well as distinct linkers from the CyanoGate system ^68^. Apart from providing an optimized cloning strategy that generates fully exchangeable genetic elements for multigene engineering, we established a new locus for efficient strain generation based on a colorimetric read-out. Applicability of the system was demonstrated at different levels, ranging from the characterization of a large promoter collection to engineering a quadruple fluorescent strain in a single step. Together, UstiGate established a quantitative and scalable foundation for multigene engineering in this basidiomycete model and biotechnology host.

## 2. Results and discussion

### 2.1. Implementation of the UstiGate system

To implement UstiGate for efficient application tailored to *U. maydis*, we broke down the highly complex MoClo system ^55,57^ to level zero (Lvl0) and level one (Lvl1), while adding the level T (LvlT) vectors from CyanoGate ^68^ as final acceptor vectors. Thus, we use a three-level hierarchy for assembly: Lvl0 harbors individual genetic elements, Lvl1 enables their assembly into functional modules such as TUs, and the terminal LvlT allows the assembly of multiple Lvl1 modules into a multigene expression construct (Fig. 1). The type IIs restriction sites within each level are compatible from one vector to the next. Their positional designation thereby reflects the order in which integrated elements or modules are directionally assembled into the next hierarchical level.

**Fig. 1:**
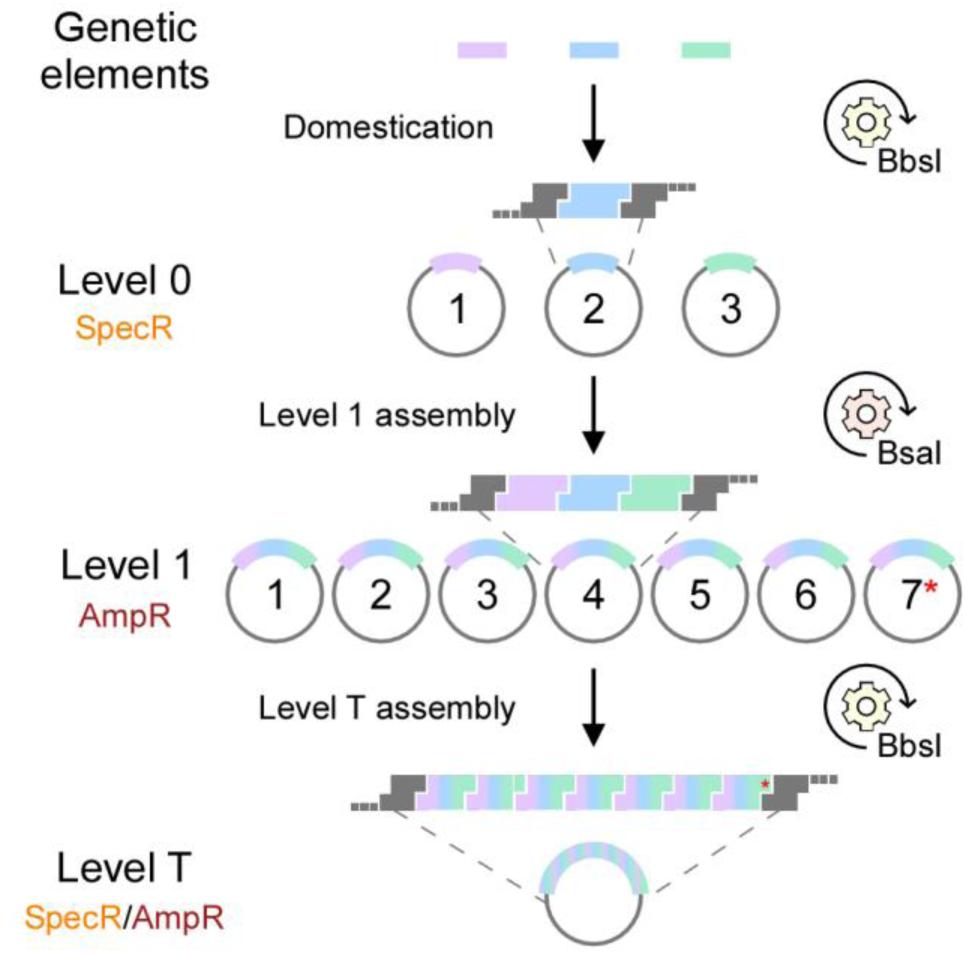
Level architecture of UstiGate in a nutshell. UstiGate is based on levels 0 and 1 of the MoClo system ^55^ and uses level T CyanoGate vectors ^68^ as final acceptors. Genetic elements are domesticated into level 0 vectors via BbsI. Three level 0 parts can be combined into each of the seven level 1 vectors using BsaI. A modified level 1 position 7 acceptor vector (7*) enables assembly of seven level 1 modules into a level T backbone without an additional end-linker. The vectors of the different levels alternately confer resistance to spectinomycin (SpecR) and ampicillin (AmpR).

UstiGate mainly employs three Lvl0 acceptor vectors, position one to three (Lvl0 P1-P3), corresponding to the original MoClo vectors pL0-PU, pL0-SC and pL0-T ^55^. Individual genetic elements first require domestication for Lvl0 integration using BbsI. This process involves the introduction of restriction sites required for subsequent cloning upstream and downstream of the element and the concurrent removal of any internal occurrences of these sites. In addition to the recognition sites of the type IIs restriction enzymes BbsI and BsaI, this also includes SspI and SfiI sites. While SspI sites are later required for the generation of linear transformation constructs with homologous blunt ends promoting homologous recombination ^26^, SfiI sites enable rapid exchange of genetic elements between the flanking regions across vectors ^28^. Once integrated into a Lvl0 vector, genetic elements are ready to use for all subsequent Golden Gate assemblies. Primer design is only required during the initial domestication.

In the next step, three Lvl0 parts can be assembled in each of the seven Lvl1 vectors (Lvl1 P1-P7; Fig. **1**) in a Golden Gate reaction with BsaI. Notably, the spatial order of Lvl1 overhangs in the MoClo system has been designed to be circular instead of linear, with the first fusion site also being the last. This design requires the use of an end-linker for assembly into the next higher level ^55^. To circumvent the standard necessity for an end-linker, we modified the Lvl1 P7 acceptor vector (Lvl1_P7_mod) so that seven Lvl1 parts can be assembled into a LvlT backbone without any additional end-linker.

The acceptor vectors of the different levels confer alternating resistance to spectinomycin (SpecR, Lvl0) and ampicillin (AmpR, Lvl1) allowing efficient counter selection against the backbones of the preceding level. For LvlT, both resistance options are available. Moreover, each vector harbors a LacZα dropout cassette enabling blue-white screening when X-Gal is used ^55,68^.

The common method to generate deletion mutants or stable genomic integrations in *U. maydis* relies on homologous recombination using sequences identical to the regions flanking the deletion or insertion site on the chromosome, the so-called upstream- (UF) and downstream (DF) flanks (Fig. 2A). Linearized constructs with homologous ends and flank lengths of around 1,000 bp thereby result in the highest recombination rate ^26^. Beside homologous flanks, the constructs contain a resistance cassette, and for insertion constructs, the transcription unit (TU) of interest (Fig. 2A).

**Fig. 2:**
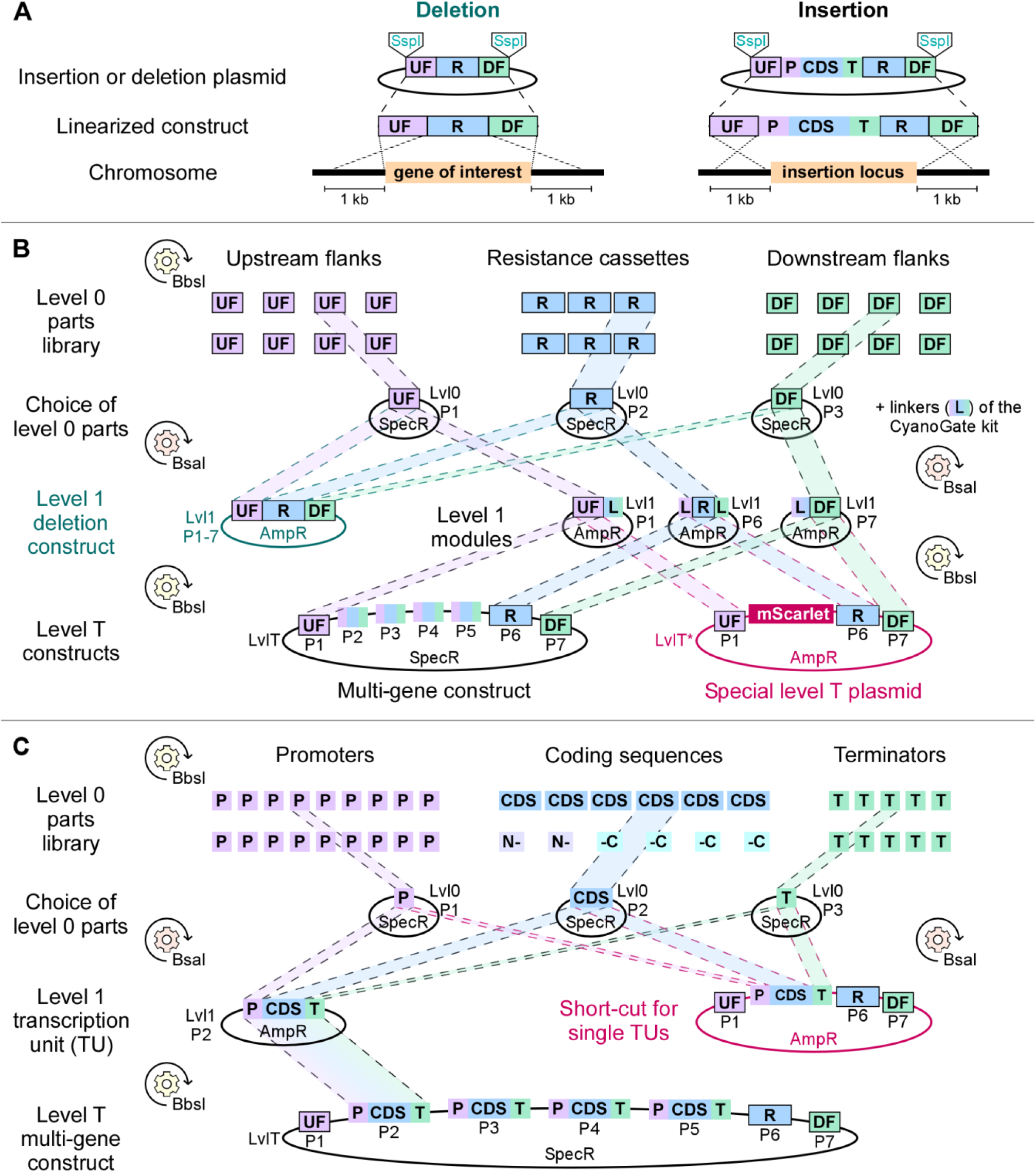
Detailed overview of the UstiGate system. **A)** General principle to generate gene deletion mutants or stable genomic integrations in *U. maydis*. **B)** Assembly of upstream flank (UF), downstream flank (DF) and resistance cassette (R) to a deletion construct and transfer of these elements via level 1 (Lvl1) using suitable linkers (L) to either a level T (LvlT) multigene construct or a special LvlT plasmid (LvlT*). P1-7, position 1 - 7. **C)** Assembly of a transcriptional unit (TU), consisting of promoter (P), coding sequence (CDS) and terminator (T), into a Lvl1 vector for subsequent transfer to a LvlT multigene construct or its direct assembly into a LvlT* plasmid.

For the generation of similar constructs with MoClo, some principles and adaptations from CyanoGate have been adopted comparable to Vasudevan et al. (2019) ^68^, who also needed to adapt the original MoClo approach for genomic modifications via homologous recombination. UFs and DFs are integrated into Lvl0 P1 and P3, respectively. Together with a resistance cassette as a Lvl0 P2 part, a simple deletion construct can be assembled in an arbitrary Lvl1 acceptor vector (Fig. 2B). To allow for linearization prior to the *U. maydis* transformation, SspI sites need to be added to the 5′end of the UF and the 3′end of the DF during the domestication. Ideally, half of the recognition site is complementary to the respective flank ends, so that SspI, as a blunt cutter, creates a fragment with perfect homologous ends ^26^.

For insertion constructs, UF, DF and the resistance cassette are transferred to Lvl1 to enable their later combination with TUs in LvlT. For that, we assigned UF and DF to the outer positions Lvl1 P1 and P7, while Lvl1 P6 is reserved for the resistance cassette of choice. The transfer of a single Lvl0 part to Lvl1 is enabled by linkers of the CyanoGate kit replacing the usually necessary Lvl0 parts (Fig. 2B; ^68^). With P1, P6 and P7 occupied, four positions (P2-P5) are still available enabling the parallel integration of up to four TUs into LvlT. Hence, instead of one ^26^ we are now able to introduce up to four TUs into the *U. maydis* genome in a single transformation step.

For the assembly of TUs, promoters, coding sequences (CDS) and terminators are first integrated into Lvl0 P1-P3, respectively. To integrate sequences for N- or C-terminal protein tags or for the generation of other variable gene fusions, we adopted the subdivision of Lvl0 P2 from the original MoClo system resulting in Lvl0 P2a and P2b, which correspond to the original vectors pL0-S and pL0-C ^55^. TUs then can be assembled in Lvl1 P2-P5 and integrated into LvlT together with the flanks of the desired insertion locus and the resistance cassette of choice as Lvl1 P1, P7 and P6 parts, respectively (Fig. 2C). For this purpose, the LvlT vector conferring SpecR should be used to enable specific selection for the final LvlT and prevent selection for Lvl1 (AmpR). In the case that less than four TUs are needed, so-called dummy fragments are available for each position replacing individual Lvl1 P1-P7 parts ^55^. Notably, we broadened the repertoire by generating six new dummies replacing two to four positions at once (Supplementary Fig. S1A), thus facilitating the assembly of final LvlT plasmids with varying numbers of TUs and less DNA parts in one assembly.

As not all applications need the complexity and variability of a standard LvlT assembly, but rather the fast and easy assembly of only one TU targeting always the same insertion locus, we developed the so-called special LvlT (LvlT*) system, a short-cut within the MoClo system. For that, the Lvl1 parts of UF, DF and resistance cassette (P1, P7 and P6) are assembled in LvlT together with an mScarlet-I dropout cassette replacing P2 to P5 and thus creating a LvlT* plasmid (Fig. 2B). The mScarlet-I dropout cassette thereby allows for pink-white screening during cloning without the necessity to add a specific substrate to the selection plates ^70^. Moreover, the mScarlet-I cassette is flanked with BsaI sites identical to the ones in Lvl1 acceptor vectors so that Lvl0 parts can be assembled directly in this LvlT* plasmid without the need for Lvl1 (Fig. 2C). To enable the efficient assembly of Lvl0 parts in LvlT*, the plasmid needs to confer AmpR instead of SpecR. This slightly impedes the generation of the LvlT* plasmid itself as the vectors of the needed Lvl1 parts also contain AmpR. However, with an additional restriction digest step after the Golden Gate reaction, the background of *E. coli* colonies harboring remaining Lvl1 vectors can be sufficiently reduced. Furthermore, *E. coli* colonies harboring the desired LvlT* plasmid can be easily identified due to their intense pink color (Supplementary Fig. S1B). Once generated, these LvlT* plasmids enable a one-step assembly of complete insertion constructs where the components of the TU can be flexibly selected. This is especially useful for high-throughput applications like screening of genetic components in projects where different enzymes or protein variants are tested or where expression conditions need to be varied.

In summary, we established a standardized MoClo-based system with a broad applicability in basic research and synthetic biology approaches. To this end, we selected, adopted and combined the best suited elements from existing systems and, where necessary, expanded the toolbox by establishing new elements and short-cuts tailored to *U. maydis* engineering. Importantly, we kept the original MoClo syntax throughout UstiGate, so that its fully compatible to further extensions. This is in contrast to alternative Golden Gate systems ^53^, for example the GreenGate system previously utilized for *U. maydis* ^47,71^. Moreover, while FungalBraid is a great opportunity for streamlining genetic manipulation in fungi, it is based on *Agrobacterium* mediated transformation ^60^. However, in the *U. maydis* community, genetic engineering is traditionally based on polyethylene glycol (PEG) mediated transformation ^25^.

### 2.2. Establishment of insertion loci with colorimetric read-out

Genomic insertion loci that enable reliable gene expression without compromising cellular fitness are essential for both fundamental research and applications in biotechnology or synthetic biology. The most popular locus for site-specific ectopic integrations in *U. maydis* is the *ip* locus ^32^, where insertion of complete linearized plasmids (including the backbone) results in two *ip* copies flanking the integrated construct and potentially causing genetic instability of the inserted TU ^23,35^. Besides, mainly the protease genes *upp3* or *pep4* have been used as integration sites and counter selection based on two different selection markers has been implemented to speed up genetic modification ^22,35^. As a parallel approach and to even further accelerate strain generation, we here established two new insertion loci where we take advantage of the color change caused by interfering with the carotenoid pathway that provides retinal as a chromophore for photoactive opsins ^72,73^. To produce retinal, geranylgeranyl-pyrophosphate (GGPP) is first converted to phytoene by the bifunctional enzyme Car2, which is then desaturated to lycopene by Car1. Subsequently, the cyclisation of lycopene to β-carotene is also catalyzed by Car2, prior to the cleavage of β-carotene by the β-carotene oxygenase Cco1, resulting in two molecules of retinal ^24,Fig. 3A; 72^. In *U. maydis* strains harboring an intact carotenoid pathway, culture cell pellets appear yellow/orange due to accumulation of carotenoids, primarily β-carotene ^Fig. 3B; 72^. Interference with the carotenoid pathway, however, results in changes in pigmentation: The deletion of *cco1* for example intensifies the yellow color due to an increase in β-carotene accumulation ^Fig. 3B; 72^, whereas the cell pellets of *car1*Δ or *car2*Δ strains appear white as the carotenoid synthesis is abolished ^Fig. 3B; 24^.

**Fig. 3:**
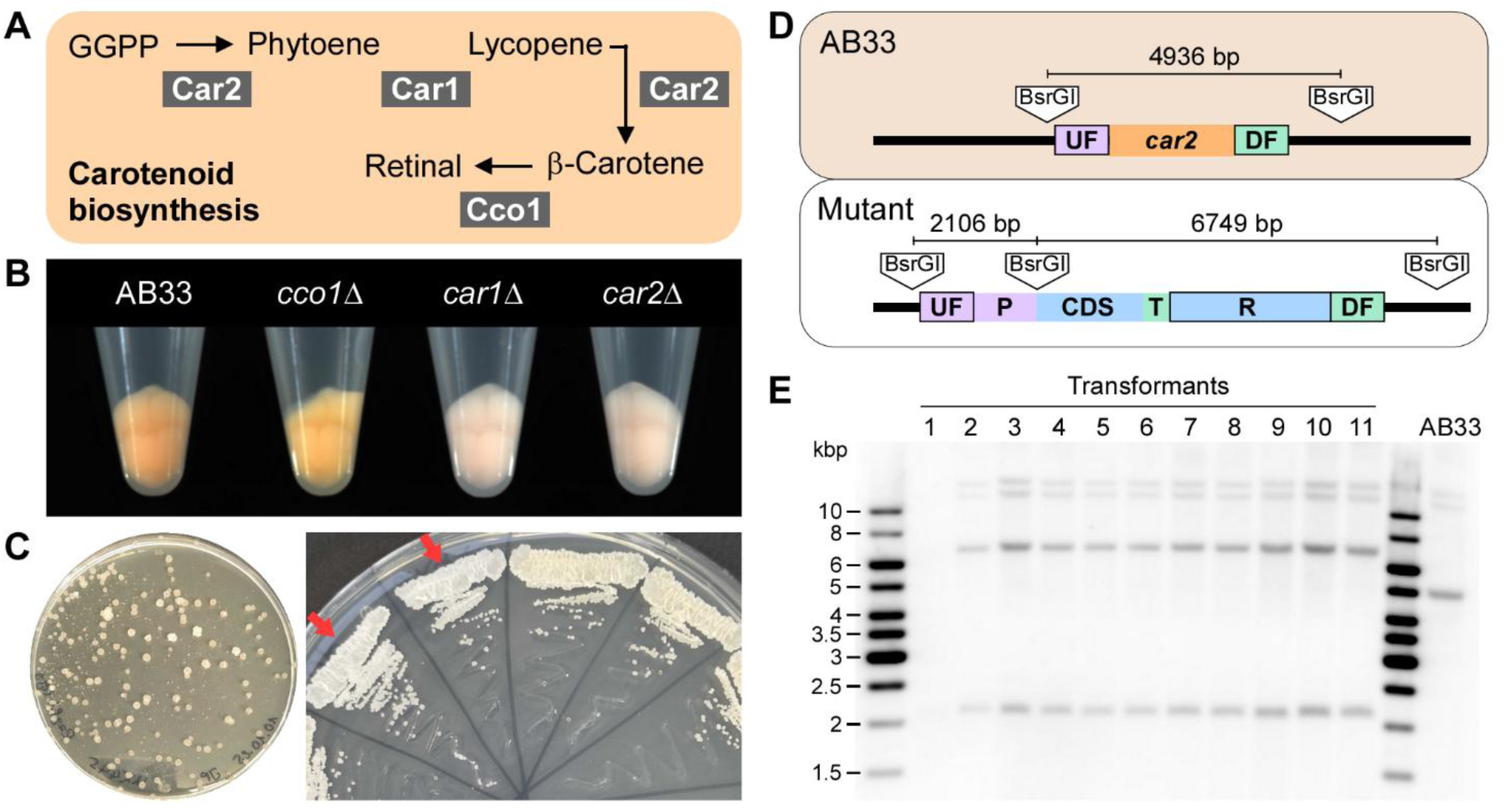
Efficient color-based strain generation by disrupting carotenoid biosynthesis. **A)** Carotenoid pathway for retinal biosynthesis in *U. maydis* starting from geranylgeranyl-pyrophosphate (GGPP). Car2, phytoene synthase and lycopene cyclase; Car1, phytoene desaturase; Cco1, β-carotene oxygenase. **B)** Cell pellets of *U. maydis* AB33 and derived deletion mutants. **C)** *U. maydis* transformation (left) and single-out plate (right) during strain generation targeting the *car2* locus. Red arrows mark white clones indicating correct integration into the *car2* locus. **D)** Schematic visualization of the *car2* locus in *U. maydis* AB33 and a derivative strain carrying an integrated expression construct. UF, DF, up- and downstream flank; P, promoter; CDS, coding sequence; T, terminator; R, resistance cassette. **E)** Example for efficient strain generation: Southern blot analysis after pre-selection of transformants by yellow-white screening with the *car2* locus as an integration site. *U. maydis* genomic DNA was digested with BsrGI and probes binding to UF and DF were used. Expected fragment sizes for progenitor strain (AB33) and correct transformants are visualized in **D**.

Here we focused on the *car2* locus, as deletion of this gene results in the most noticeable color change. While the color difference is not reliably visible on transformation plates, whitish transformants can easily be selected for further verification after re-plating for isolation of single colonies (Fig. 3C). Alternatively, harvested cell pellets of small cultures clearly show the color difference, and proceeding to Southern blot analysis with white clones frequently identifies approximately (approx.) 90 % positives (Fig. 3D, E).

The intense yellow color caused by *cco1* deletion is less reliable, as the color difference is sometimes subtle, but can be enhanced by incubating the agar plates at 4 °C for some days. A sequential use of the loci, targeting *cco1* first and then *car2*, is a good strategy when two insertion loci are required.

Importantly, eliminating retinal synthesis has no effect on growth, morphology or pathogenicity under standard laboratory conditions ^72^. Still, fungal carotenoids can for example have a protective role against oxidative stress and exposure to visible light or ultraviolet irradiation ^74^. Moreover, they can constitute intermediates in the synthesis of biological active compounds besides retinal, such as the sexual pheromones trisporic acids ^74^. In fundamental research, it is therefore important to consider that abolishing carotenoid synthesis may affect cellular function depending on the biological context. However, disruption of pigment biosynthesis as a visual marker for identifying correctly integrated clones has previously been applied in other fungi in the biotechnology or synthetic biology field, including the black fungus *Knufia petricola* ^75^ or the two basidiomycete red yeasts *Rhodosporidium toruloides* ^76^ and *Xanthophyllomyces dendrorhous* ^77^. The use of *car2* as insertion locus therefore provides a powerful color-based screening tool that substantially accelerates *U. maydis* strain generation and can be widely applied in the biotechnological and synthetic biology context.

### 2.3. UstiGate-based characterization of promoters at single-cell level

While a small selection of promoters is used for gene expression in the *U. maydis* community, a systematic and comprehensive characterization was lacking. Here, we used the power of the modular system and integrated more than 20 promoters in the UstiGate toolbox for their characterization in

*U. maydis*. Each promoter was assembled with *eGFP* ^78^ and the terminator of the nopaline synthase gene (*nos*) from *Agrobacterium tumefaciens* ^Tnos; 38^ in a LvlT* plasmid targeting the *car2* locus (Fig. 4A). Employing yellow-white screening allowed efficient generation of the numerous strains. In general, we aimed for a broad repertoire of constitutive promoters. We included the already published constitutive promoters, derived from the native genes for heat shock protein 70 (Phsp70) ^36^, glyceraldehyde 3-phosphate dehydrogenase (Ptdh3 = Pgap) ^37^, hypothetical sugar phosphate phosphatase (Psgp1 = P_UMAG_05521_) ^47^, ribosomal proteins of the large subunit (Prpl10 and Prpl40) ^24^ and translation elongation factor (Ptef) ^38^ as well as the latter’s synthetic derivative Potef ^38^. In addition, we added 13 putative native promoters, corresponding to genes involved in housekeeping processes (Supplementary Table S1). To this end, we relied on the available mRNA sequencing dataset for *U. maydis* grown in axenic culture ^79^ (Supplementary Fig. S2) and searched for example for homologs of *S. cerevisiae* proteins whose underlying genes are natively expressed by strong promoters as described for the yeast toolkit ^64^. For most promoters, a 1,000 bp region upstream of the start codon was selected and domesticated into UstiGate Lvl0 P1 (Supplementary Table S1).

**Fig. 4:**
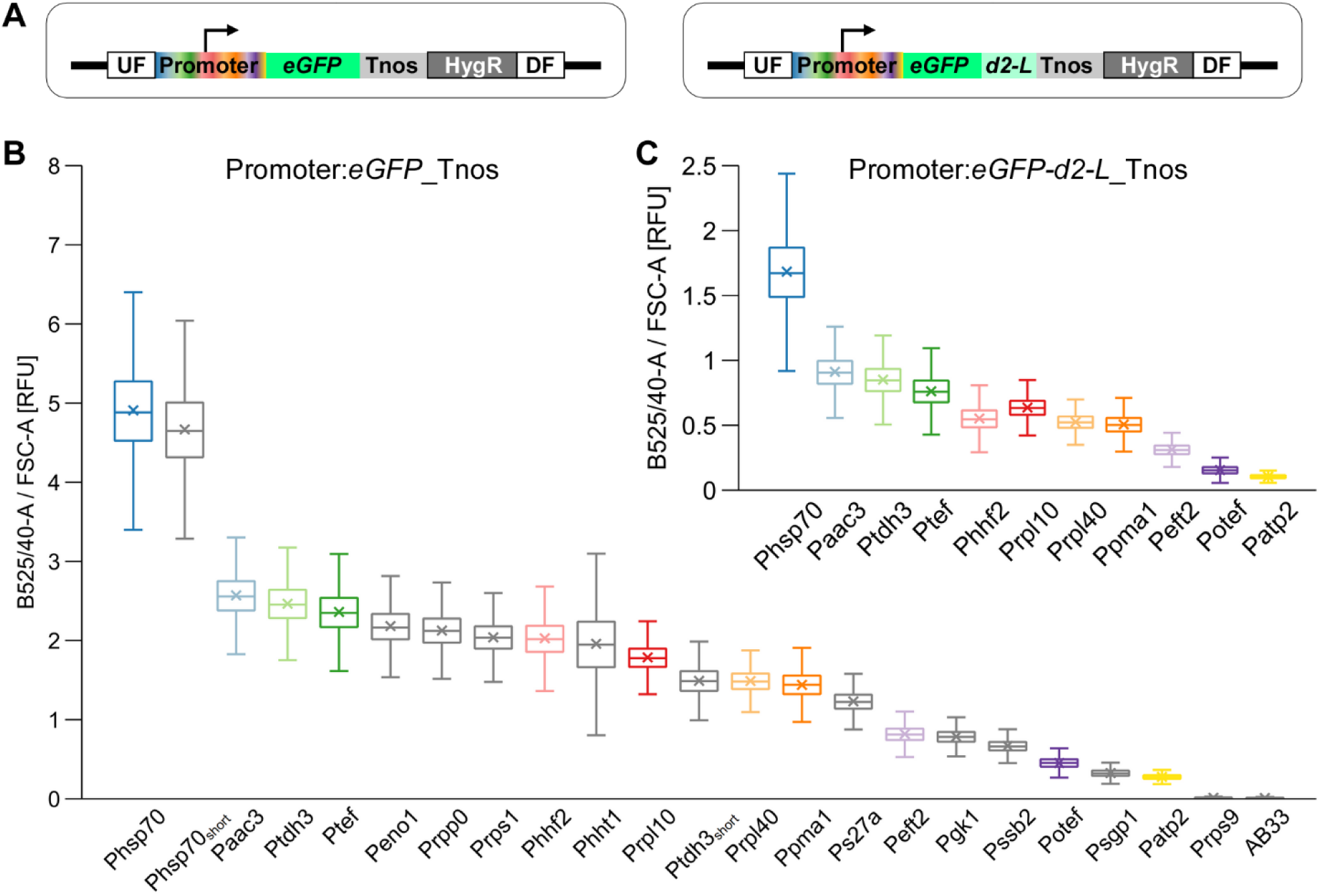
Comparison of different promoters at the single-cell level. **A)** Schematic overview of the genetic setup of the reporter strains. Putative promoter regions were fused with e*GFP* (left) or a gene fusion yielding a destabilized eGFP variant (e*GFP-d2-L*, right) and integrated into the *car2* locus. UF, DF, up - and downstream flank of the *car2* locus; Tnos, nopaline synthase terminator; HygR, hygromycin resistance cassette. **B, C)** eGFP reporter strains were analyzed during the exponential growth phase (OD_600_ ∼1) on single-cell level using flow cytometry. eGFP fluorescence intensity of each cell was determined in the B525/40-A channel and normalized to forward-scatter area (FSC-A) to account for differences in cell size. Boxplots show the statistics including the arithmetic mean (x) of ∼ 30,000 cells per strain analyzed in three independent experiments. Range of the whiskers indicate population variability referring to the extent of dispersion around the median. **C)** Approx. half of the promoters (shown with colored boxplots in **B** were additionally screened with the destabilized eGFP variant (eGFP-d2-L).

Beside determining bulk fluorescence in a plate reader (Supplementary Fig. S3), we analyzed the eGFP reporter strains at single-cell level using flow cytometry, providing insights not only into the average fluorescence intensity of a population, but also into its variability referring to the extent of dispersion around the median. For each strain, a total of 30,000 yeast-like cells were analyzed during the exponential (Fig. 4B) and the stationary growth phase (Supplementary Fig. S4). Except for the Prps9 strain, all other 21 promoter reporter strains exhibited clearly higher eGFP fluorescence in the exponential growth phase as compared to the progenitor strain (AB33), indicating promoter activity (Fig. 4B). A broad range of fluorescence read-outs was detected, with the strongest promoter, Phsp70, leading to a nearly 18-fold stronger fluorescence than Patp2 at the very low end. Notably, we identified several promoters with very similar expression strengths that are interchangeable and thus can reduce the risk of undesired homologous recombination in multigene constructs due to sequence repeats. In regard to population variability, only the strain with Phht1 deviates from the other reporter strains: Normalized to their overall fluorescence intensity, the dispersion of the Phht1 reporter population is nearly twice as high compared to the majority of the other promoters (Supplementary Fig. S5).

To ensure that eGFP accumulation, resulting from its high stability, did not distort the results, a destabilized eGFP variant was generated by fusion to a degron sequence (eGFP-d2-L) and used to further characterize 11 of the 21 promoters in a near real-time manner. Apart from the expected overall reduced fluorescence levels, the distribution of promoter strengths remained largely unchanged, with the exception of Phhf2, where fluorescence appeared slightly weaker in comparison to the degron-free version (Fig. 4C). Of note, the destabilized eGFP version for the first time also enables to monitor the promoter activity in the stationary growth phase (Supplementary Fig. S4), which was overall reduced while fluorescence levels of Phhf2 and Ppma1 stayed comparatively elevated. Moreover, the population variability clearly increases when growth stagnates.

Interestingly, the unique approach revealed that the frequently applied Potef promoter exhibits comparatively low expression strength. In this artificially assembled promoter, tetracycline-responsive elements were reported to enhance the activity of the *tef* core promoter by approx. 8-fold ^38^, which contradicts our findings showing that Ptef is 5-fold stronger than Potef. A second unexpected finding concerns Ptdh3 and Phsp70. The latter is described to exhibit a strong basal activity that can be further stimulated under stress conditions ^36,80^. Both promoters were included in the promoter screen in two different lengths (Supplementary Table S1): an approx. 1,000 bp version and a shorter version commonly used to drive resistance cassettes. For Phsp70, both versions approx. exhibit the same expression strength whereas Ptdh3 shows 1.6-fold higher expression strength than Ptdh3_short_ (Fig. 4B).

In conclusion, we comparatively characterized existing promoters under constant settings for the first time in *U. maydis* and further expand the promoter library with a broad set of additional promoters. This significantly advances the selection of promoters for genetic engineering of *U. maydis*.

### 2.4. Promoter characterization across different media

While cytometry provides detailed information on fluorescence levels in individual cells within a population, it is limited to specific measurement timepoints. To continuously monitor promoter activity during the growth of *U. maydis*, we cultivated the reporter strains of the selected 11 promoters in a microbioreactor (BioLector), enabling continuous measurement of scattered light, as a read-out for fungal biomass, and fluorescence. Moreover, we tested three different commonly used media: A defined mineral medium (Verduyn), a complete medium (CM), both supplemented with 10 g L^-1^ glucose, and a modified Verduyn medium containing excess glucose and lower nitrogen content to induce nitrogen limitation (Verduyn-N). Importantly, although the overall growth differs in speed and final biomass depending on the medium, all promoter reporter strains exhibit similar growth behavior within each medium (Supplementary Fig. S6). Moreover, the majority of promoters also maintain their relative expression strengths, particularly during exponential growth, independent of the medium, although overall absolute strengths may change (Fig. 5). The distribution of the promoter strengths in Verduyn medium, also used for the previous cytometry measurements, is largely consistent with the cytometry results throughout the cultivation period (Fig. 5A, B). The fluorescence read-outs of the eGFP-d2-L strains clearly demonstrate, in near real-time, that overall promoter activity is immediately reduced upon entry into the stationary growth phase (Fig. 5B, Supplementary Fig. S6). Even the comparatively elevated fluorescence levels of Phhf2 and Ppma1 determined in the cytometer during the stationary growth phase (Supplementary Fig. S4) can clearly be observed. For these strains, fluorescence increases slightly later, but promoter activity seems to persist longer resulting in comparatively higher fluorescence during the stationary phase (Fig. 5A, B). This phenomenon is even more pronounced in CM medium, where Paac3 additionally exhibits comparatively higher fluorescence levels in the later growth phase (Fig. 5C, D).

**Fig. 5:**
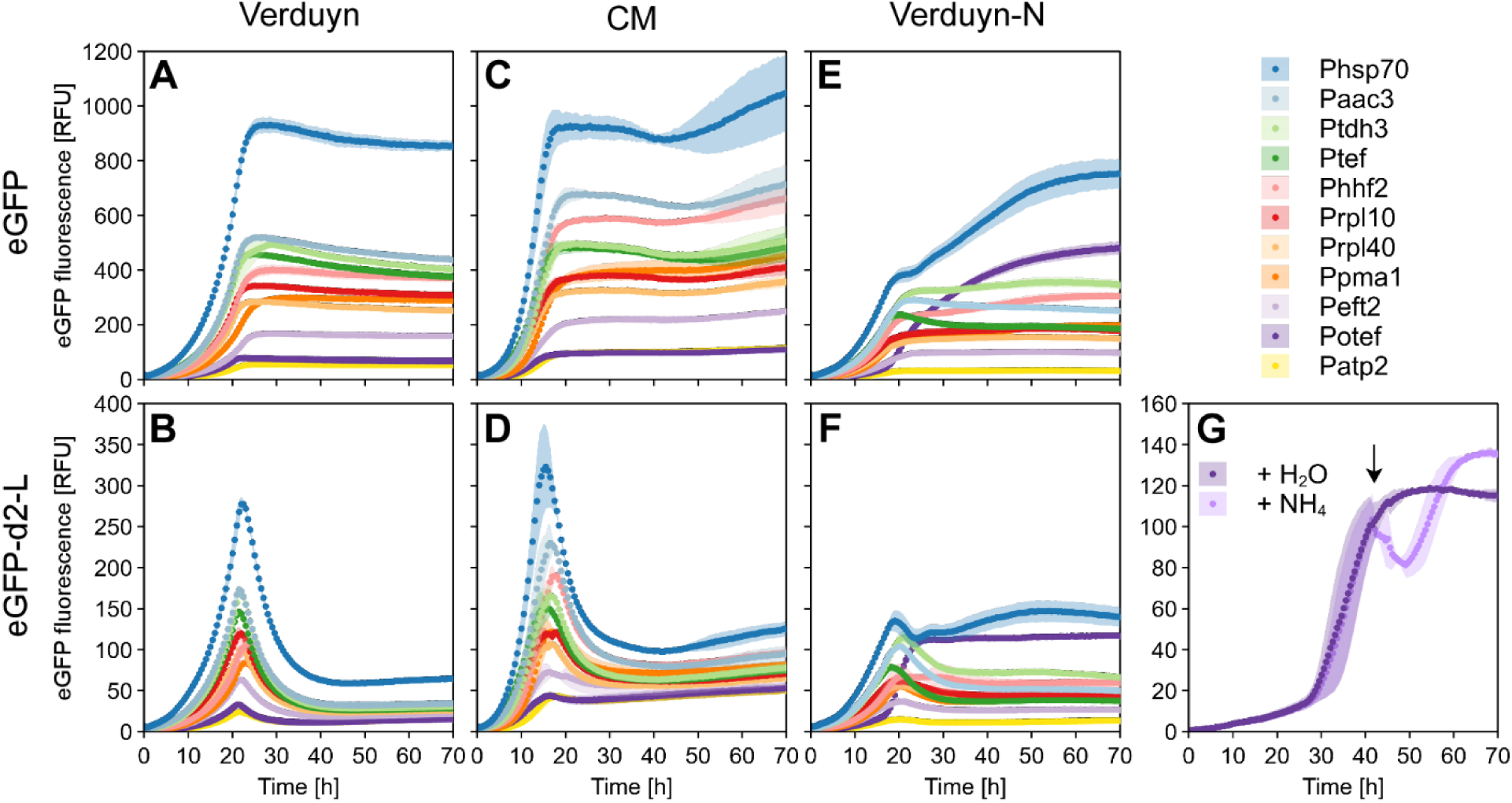
Online monitoring of promoter activity in different media. eGFP (**A**, **C**, **E**) or a destabilized version (eGFP-d2-L; **B**, **D**, **F**) were used as reporters with a selection of promoters. Cells were grown in Verduyn mineral medium (**A**, **B**), complete medium (CM; **C**, **D**) or a modified Verduyn variant with excess glucose and reduced nitrogen content (Verduyn-N; **E**, **F**). Online monitoring was performed in a microbioreactor (BioLector) measuring backscatter (gain 20), as a read-out for fungal biomass, and eGFP fluorescence (gain 70). Backscatter data of the cultivations are provided in Supplementary Fig. S6. **G)** Potef is throttled by nitrogen supplementation as demonstrated with spiking in ammonium at about 42 h of cultivation (arrow). The graphs show arithmetic mean and standard deviation of biological triplicates (**A**, **B**, **E**, **F**) or duplicates (**C**, **D**, **G**) across independent experiments.

In Verduyn-N medium, the promoter distribution during the exponential growth is nearly identical to that observed in Verduyn and CM. However, the transition into the stationary phase is accompanied by some changes, likely because growth in this medium stagnates due to nitrogen-limitation rather than glucose depletion. Next to Phsp70 and Phhf2, Ptdh3 exhibits comparatively high promoter activity while Ptef is reduced in strength (Fig. 5E, F). In contrast, the synthetic derivative of the latter, Potef ^38^, displays a substantial rise in promoter activity upon entrance into the stationary growth phase. Apparently, Potef is induced upon nitrogen limitation, becoming the second strongest promoter during the late growth phase (Fig. 5E, F). To validate this unexpected observation, we supplemented a nitrogen-limited Potef^d2-L^ culture with ammonium, clearly demonstrating that Potef activity is throttled by nitrogen availability (Fig. 5G). This observation is of particular interest, as most secondary metabolites and microbial oil are produced upon nitrogen limitation where Potef could be suited for genetic upregulation during engineering ^5,16,81^

In summary, albeit distinct nutrient limitations during the stationary phase can lead to promoter-specific responses, the relative activity of the tested native *U. maydis* promoters is mostly robust throughout cultivations in different media, which is consistent with a large-scale study in *S. cerevisiae* showing that most promoters are globally scaled across changing growth conditions while maintaining their relative expression strengths ^82^.

### 2.5. Expression strength and population variability in different loci

To analyze whether the insertion locus influences promoter strength and to compare the two new insertion loci, *car2* and *cco1*, with the established protease loci (*upp3* and *pep4*) regarding population variability, we selected four promoters spanning our promoter spectrum and integrated respective eGFP reporter TUs into all four loci. In the exponential growth phase, Phsp70_short_ and Paac3 exhibit the highest expression strength at the *upp3* locus, closely followed by *car2* and *pep4* (Fig. 6). At the *cco1* locus, both promoters exhibit only 83 % activity. In contrast, Prpl10 nearly shows identical activity across all four loci, with only 1 - 3 % variation between the loci. Potef exhibits the highest activity at the *upp3* locus, followed by *car2* and *cco1*. By far the lowest values for Potef were determined at the *pep4* locus, reaching only 59 % of the activity observed at the *upp3* locus. However, this strain also displays 50 % higher population variability than the other Potef strains, which may contribute to the reduced median value (Supplementary Fig. S7A). Apart from Potef^pep4^, the *pep4* strains rather show the least variations, indicating that the locus itself does not inherently cause increased variability. Overall, with this single exception, no substantial differences in population variability were observed between the loci in exponentially growing yeast-like cells as well as in the stationary growth phase (Supplementary Fig. S7A, B). Regarding promoter strength across the different loci, only Phsp70_short_ and Paac3 show comparatively lower activity at the *pep4* locus in the late growth phase (Supplementary Fig. S8), otherwise, the overall pattern remains consistent with that observed during exponential growth.

**Fig. 6:**
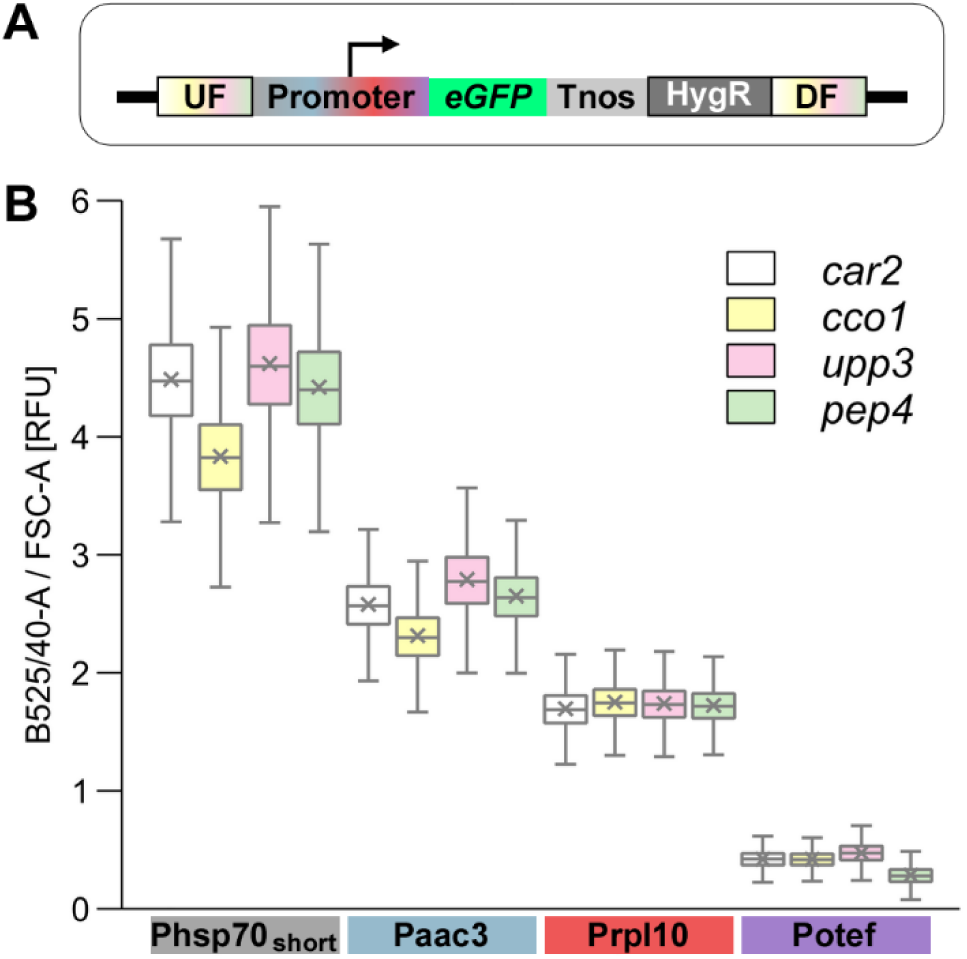
Comparative evaluation of different insertion loci. **A)** Schematic representation of the genetic setup of the utilized reporter strains. Phsp70, Paac3, Prpl10 and Potef were fused with *eGFP* and integrated into the *car2*, *cco1*, *upp3* and *pep4* locus, respectively. UF, DF, up- and downstream flank of the four loci; Tnos, nopaline synthase terminator; HygR, hygromycin resistance cassette. **B)** Comparison of four different insertion loci. eGFP reporter strains were measured during the exponential growth phase (OD_600_ ∼1) on single-cell level using flow cytometry. eGFP fluorescence intensity of each cell was determined in the B525/40-A channel and normalized to forward-scatter area (FSC-A) to account for differences in cell size. Boxplots show the statistics including the arithmetic mean (x) of ∼ 30,000 cells per strain analyzed in three independent experiments. Range of the whiskers indicate population variability referring to the extent of dispersion around the median.

Altogether, the results indicate that the insertion locus generally does not have a substantial effect on expression strength or population variability. This is in accordance with observations in *S. cerevisiae* where the use of two separate loci resulted in expression distributions nearly identical to those obtained using of a single locus ^64^. However, usage of the *otef* promoter at the *pep4* locus may not be advisable, although, a potential role of e*GFP* in the observed phenomenon can also not be excluded as, to our knowledge, no other reporter has yet been tested in this context. Remarkably, the population variability reported for the *pep4* locus in AB33 hyphae ^35^ is also based on a Potef:*eGFP* construct and may potentially be circumvented by using a different promoter or (fluorescent) protein.

### 2.6. Fluorescent reporters and their impact on relative promoter activity

As fluorescent reporter proteins are highly valuable tools in both cell and synthetic biology ^83,84^, we aimed to expand the UstiGate toolkit with a diverse palette of FPs. To this end, we individually integrated twelve Prpl10-driven fluorescent reporter cassettes into the genome of *U. maydis*, using the LvlT* plasmid for *car2* integration. We assessed their functionality and compared their fluorescence intensities by bulk fluorescence measurements using their near-optimal Ex and Em wavelengths (Fig. 7; Supplementary Table S2). While the utilized CDS for the blue FP Electra2 ^85^ and the cyan FP mTurquoise2 ^86^ did not result in fluorescence levels above the background fluorescence of cells without an FP, all other ten FPs were detectable (Fig. 7A, Supplementary Fig. S9). However, their fluorescence intensities differ by up to two orders of magnitude, with eGFP and the yellow FP mVenus ^87^ showing by far the highest fluorescence intensities of nearly 10^5^ RFU. The red FPs mCherry ^88^ and mKate2 ^89^, both well established in *U. maydis* ^47,49,50,81,90–92^, exhibit second highest fluorescence intensities around 10^4^ RFU followed by the large Stokes shift (LSS) FPs mT-Sapphire* ^93^ and LSSmOrange ^94^. At the lower end of the intensity spectrum are mTagBFP2 ^95^ and LSSmApple ^96^ as well as the synthetic RFP mScarlet-I ^97^ and LSSmKate2 ^98^, with the latter two exhibiting the lowest fluorescence intensities with values below 10^3^ RFU. Overall, we established several new fluorescent reporters for *U. maydis*, integrated a broad palette of FPs into the UstiGate system, and provide fluorescence intensity data that, together with their spectral properties, provide a valuable basis for selecting suitable FPs for specific applications.

**Fig. 7:**
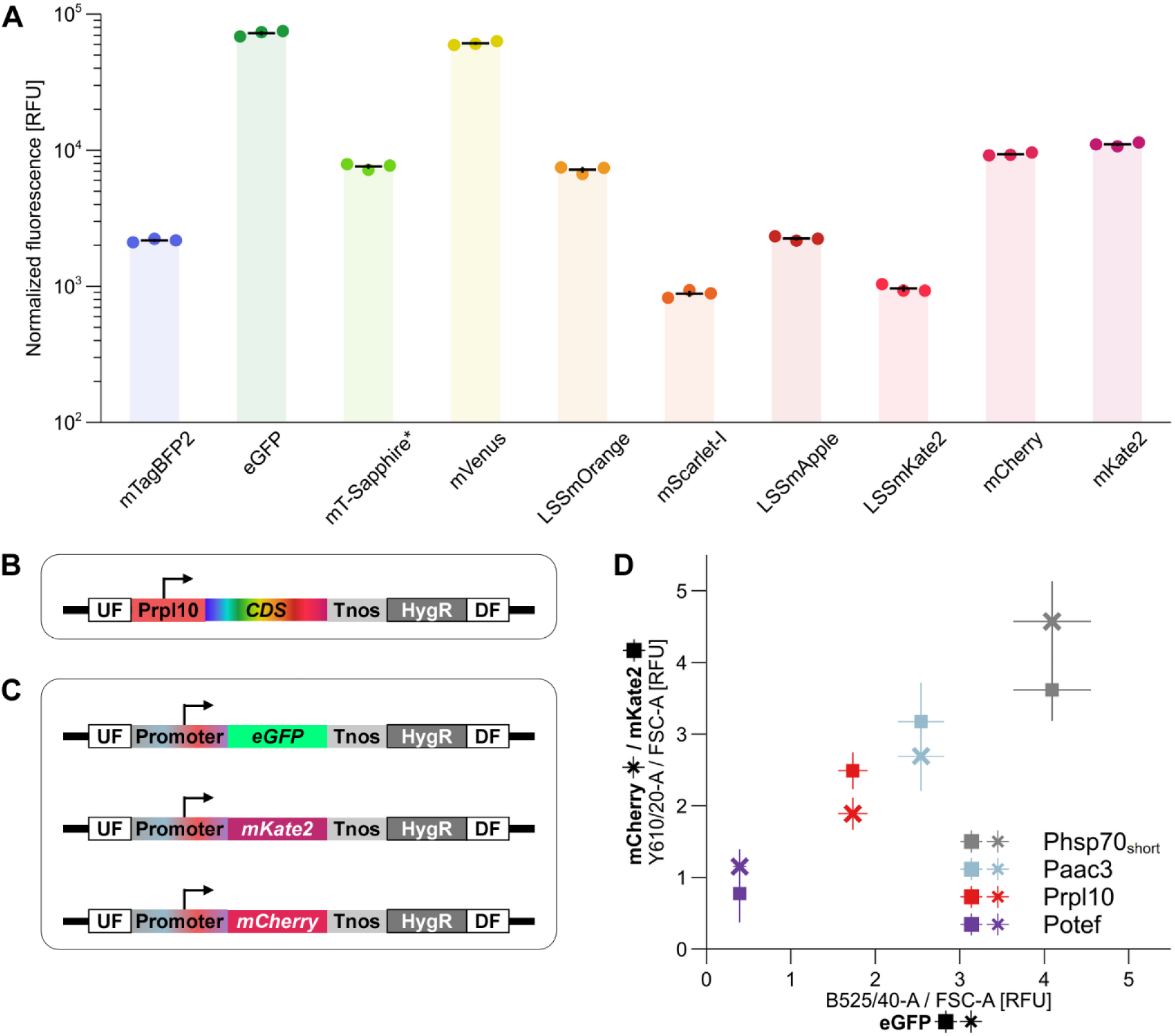
Expansion of the fluorescent reporter spectrum and impact of reporter CDS on relative promoter strength. **A)** Comparative evaluation of the fluorescence intensity of ten fluorescent proteins (FPs) in *U. maydis*. FPs were evaluated by measuring bulk fluorescence of undiluted, exponentially growing cultures in a plate reader. Fluorescence was measured at near-optimal excitation and emission wavelength for each FP (see Supplementary Table S2) while using a constant gain setting of 150. Fluorescence values were normalized to OD_600_ and the background fluorescence of cells without an FP was subtracted. Each dot represents an individual biological replicate from independent experiments, and black lines indicate the arithmetic mean ± standard deviation. mT-Sapphire*, mT-Sapphire with the following mutations: N147T S148A L232H. **B)** Schematic representation of the genetic setup of the utilized strains. CDSs of the FPs were fused downstream of Prpl10 and integrated into the *car2* locus. UF, DF, up- and downstream flank of the *car2* locus; Tnos, nopaline synthase terminator; HygR, hygromycin resistance cassette. **C)** Genetic setup used to analyze the impact of the reporter CDS on relative promoter strength. Four promoters, Phsp70_short_, Paac3, Prpl10 and Potef, were each combined with *eGFP*, *mCherry* and *mKate2* and integrated into the *car2* locus. **D)** Green (eGFP) or red (mCherry, mKate2) fluorescence intensity of cells in the exponential growth phase was determined using the B525/40-A or the Y610/20-A channel of the flow cytometer, respectively, and fluorescence values were normalized to forward-scatter area (FSC-A) to account for differences in cell size. The arithmetic mean ± standard deviation of ∼ 30,000 cells per strain, analyzed in three independent experiments, is plotted with normalized fluorescence of the eGFP reporter strains on the x- and the corresponding values of the mCherry- or mKate2-producing strains on the y-axis.

To analyze potential effects of the downstream reporter CDS on the relative promoter strength, we integrated *mCherry* and *mKate2* into *U. maydis* downstream of the same four promoters previously tested at the different loci with *eGFP* (Fig. 7C). We selected the two RFPs as alternative reporters, as they exhibit comparably strong fluorescence intensities and together with eGFP, the three FPs constitute distinct CDS as they are derived from different ancestral proteins ^78,88,89^. The normalized fluorescence of the eGFP strains and the respective values of the mCherry or mKate2 strains generally show a linear trend, but there are two exceptions: Phsp70_short_ shows lower relative promoter strength with *mKate2* and Potef shows higher relative strength with *mCherry* (Fig. 7D, Supplementary Fig. S10). However, consistent with findings in *S. cerevisiae* ^64,99,100^, the relative expression strength is largely independent of the CDS, indicating that our broad screen based on eGFP provides a reliable read-out for gene expression strength.

### 2.7. A novel autoinduction promoter based on maltose

In addition to constitutive promoters, inducible promoters have a wide range of applications enabling for example the functional characterization of essential genes in the laboratory or the separation of cell growth and production phase in industrial processes ^101,102^. Beside tetracycline-controlled expression systems ^47,48^, two native nutrient-dependent inducible promoters have previously been characterized in *U. maydis*. The nitrate-inducible and ammonium-repressed Pnar1 promoter of the nitrate reductase gene ^40^ is for example critical for controlled filamentous growth in the widely used laboratory strain AB33 ^41,103^. Pcrg is carbon-source regulated, namely induced by arabinose and repressed by glucose ^39^, and can for example be exploited for autoinduction, as *U. maydis* preferentially consumes glucose before switching to arabinose ^104^.

Here, we established a second native autoinduction promoter based on a glucose-maltose switch. Expression of *IMA1*, encoding the major isomaltase in *S. cerevisiae*, has been reported to be strongly induced by maltose ^105^ and a homolog of Ima1 has recently been described for *U. maydis* ^106^. Approx. 3 kb-long promoter regions of *ima1* and *crg* were integrated into the UstiGate system to generate eGFP fusions (Fig. 8A) and the subsequently generated reporter strains exhibited markedly increased eGFP fluorescence when cultivated on arabinose or maltose, respectively, compared to glucose (Fig. 8B). Regarding promoter strength, both promoters rank between the strongest constitutive promoters (Fig. 4), with Pima1 being 1.3-fold stronger than Pcrg. Moreover, for both promoters, the onset of eGFP fluorescence is gradually shifted to later time points with increasing glucose proportions in the culture (Fig. 8C, D), although growth initiated simultaneously (Supplementary Fig. S11). This indicates that Pima1 is, like Pcrg, glucose-repressible. To verify this, we added glucose to arabinose/maltose cultures of eGFP-d2-L reporter strains and thus enabled near real-time observation of glucose-repression. In both strains, Pcrg^d2-L^ and Pima1^d2-L^, fluorescence decreased immediately upon glucose addition and increased again only after approx. 10 hours, likely as a consequence of glucose depletion (Fig. 8E, F). In contrast, the addition of arabinose or maltose did not cause a decline in fluorescence and biomass formation was nearly identical (Supplementary Fig. S12).

**Fig. 8:**
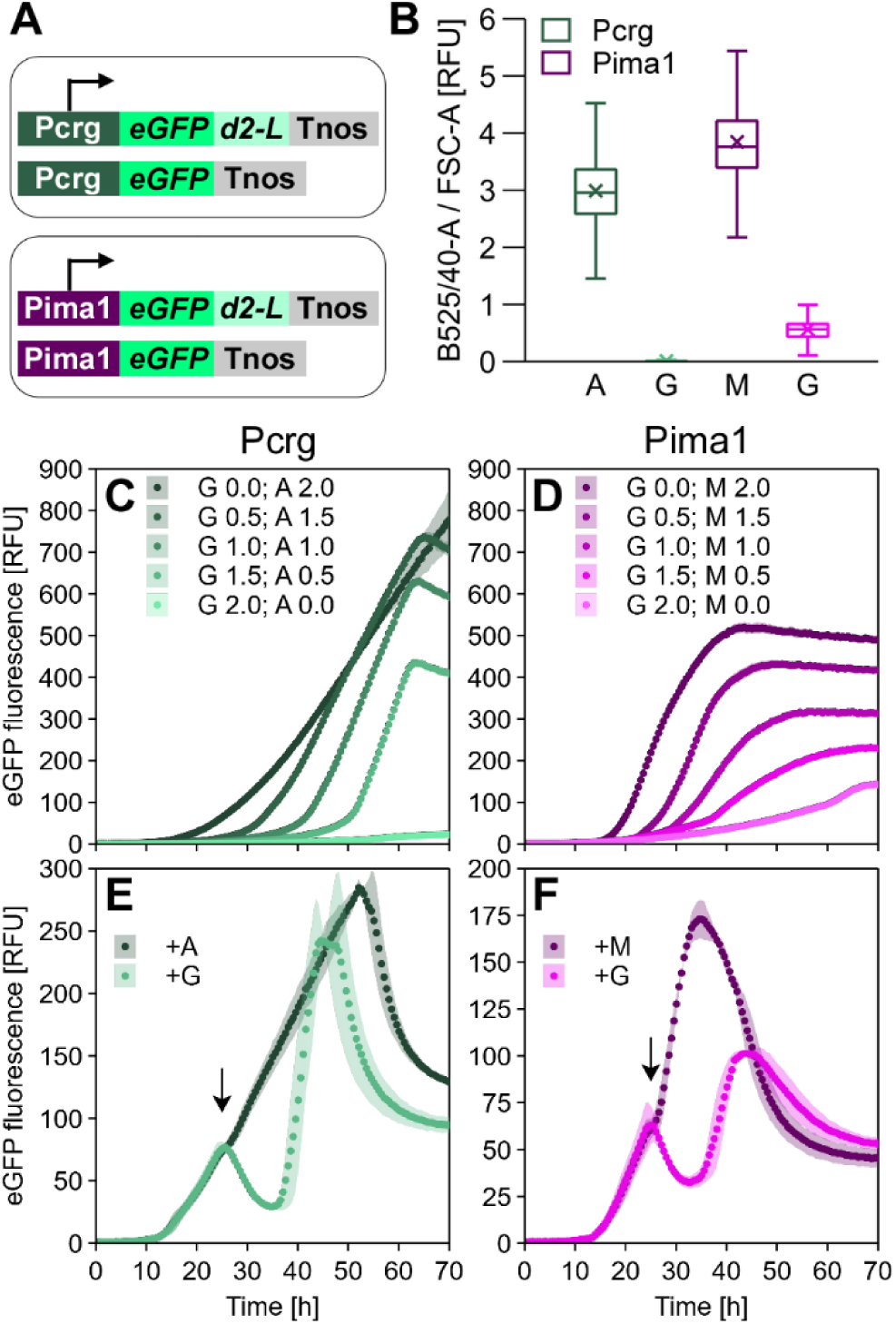
Characterization of the novel auto-induction promoter Pima1. **A)** Schematic representation of the genetic setup of the utilized reporter strains. Pcrg and Pima1 were combined with *eGFP* with or without degron tag (*d2-L*) and integrated into the *car2* locus. Tnos, nopaline synthase terminator. **B)** eGFP reporter strains were cultivated with arabinose (A) / maltose (M) or glucose (G) and measured on single-cell level using flow cytometry. eGFP fluorescence intensity of each cell was determined in the B525/40-A channel and normalized to forward-scatter area (FSC-A) to account for differences in cell size. Boxplots show the statistics including the arithmetic mean (x) of ∼ 30,000 cells per strain analyzed in three independent experiments. Range of the whiskers indicate population variability referring to the extent of dispersion around the median. **C**, **D**) Comparative eGFP fluorescence online monitoring during growth. Biomass data of the cultivations are provided in Supplementary Fig. S11. Cells were grown in pure glucose (G 2.0; A/M 0.0) or pure inducing sugar arabinose or maltose (G 0.0; A/M 2.0), as well as different ratios of both sugars. For both promoters eGFP fluorescence increases later with increasing glucose proportion in the culture. **E, F)** Pcrg and Pima1 are repressed by glucose as demonstrated with eGFP-d2-L reporter strains growing on arabinose and maltose, respectively, and the addition of glucose (G) in comparison to arabinose (A) or maltose (M) at about 25 h of cultivation (arrow). The graphs (**C**-**F)** show arithmetic mean and standard deviation of biological duplicates across independent experiments.

Notably, Pima1 exhibits a higher basal expression level than Pcrg, which increases during cultivation on pure glucose (Fig. 8B, C, D). This may indicate that Pima1 is not solely maltose-induced, but also slightly active under low-glucose conditions or during carbon starvation. De-repression by carbon source depletion has been reported for numerous carbon source dependent promoters in different yeast species, such as P*_SUC2_* of *S. cerevisiae* ^107,108^. As Ima1 exhibits significant sucrose-hydrolytic activity ^106^, we also tested sucrose as alternative inducer. The results reveal a concentration-dependent induction of Pima1, but with expression levels only slightly above the baseline (Supplementary Fig. S13). A similar pattern is observed with maltose as inducer when a shortened Pima1 variant with only 1,000 bp (Pima1_short_) is used (Supplementary Fig. S13).

Taken together, the results establish Pima1 as a novel glucose-repressed and maltose-inducible promoter suitable for strong autoinduction in *U. maydis*. Compared to Pcrg, this offers the advantage that maltose, a disaccharide composed of two glucose molecules, is metabolized identical to glucose after hydrolysis and therefore does not require a major metabolic shift. In contrast, arabinose catabolism proceeds through four alternating reduction and oxidation reactions followed by phosphorylation and results in d-xylulose-5-phosphate, an intermediate of the pentose phosphate pathway ^109^. The system thus represents a useful addition to the comparable Pcrg system, exhibiting higher basal expression levels but offering the advantage of a reduced metabolic shift.

### 2.8. Comparative evaluation of terminators

To reduce the risk of undesired homologous recombination in multigene plasmids due to repeated sequences, we integrated seven different terminators into the UstiGate toolbox (Supplementary Table S3). Beside the primarily used *nos* terminator ^38^, we domesticated the native terminator from the heat shock protein 70 (Thsp70) and Tcyc1 from the iso-1-cytochrome c from *S. cerevisiae*, which are employed in the HygR and NatR resistance cassettes ^28^, respectively, as well as Tprm9 from a yeast pheromone-regulated membrane protein ^47,110^. Moreover, three new native terminators were implemented using the downstream region of the translation elongation factor gene (Ttef) and of the two genes encoding for the 60S ribosomal proteins L10 and L40 (Trpl10 and Trpl40). We assembled all terminators downstream of a Prpl10:e*GFP* reporter fusion in the LvlT* plasmid for *car2* integration to analyze their influence on gene expression (Fig. 9A). The data suggest that the terminator indeed influences eGFP accumulation, with differences of up to 2.7-fold (exponential phase) and 3-fold (stationary phase), and Ttef and Tnos being on the upper and Tcyc1 on the lower end (Fig. 9B; Supplementary Fig. S14). The influence of the tested terminators on expression strength may also be transferable to other promotors and CDSs, as demonstrated in the yeast toolkit study analyzing five terminators in combination with three different promoters and FPs ^64^. However, for applications requiring precise expression levels, we recommend characterizing individual promoter-CDS-terminator combinations to ensure that the desired expression strength is obtained.

**Fig. 9:**
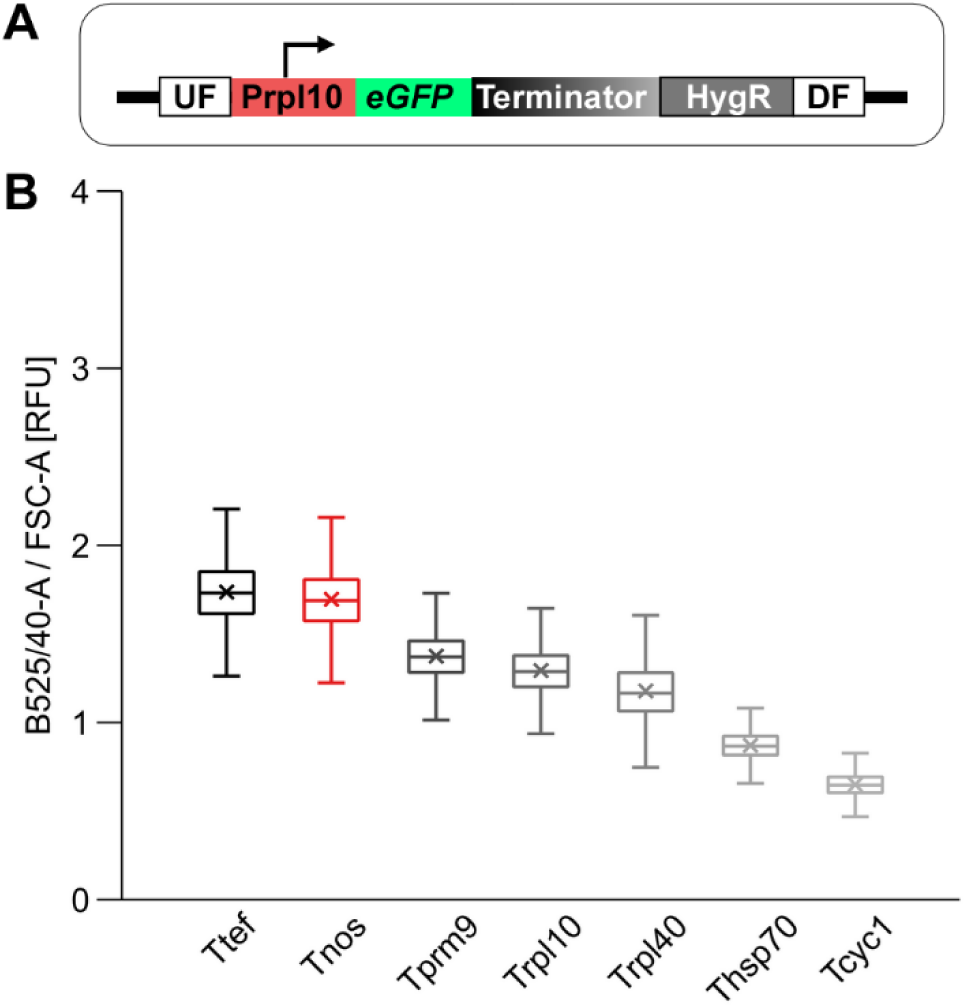
Comparative evaluation of different terminators. **A)** Schematic representation of the genetic setup of the utilized reporter strains with varying terminator sequences (indicated by shades of grey). Terminators were fused downstream of a Prpl10:*eGFP* reporter construct and integrated into the *car2* locus. UF, DF, up- and downstream flank of the *car2* locus; HygR, hygromycin resistance cassette. **B)** eGFP reporter strains were measured during the exponential growth phase (OD_600_ ∼1) on single-cell level using flow cytometry. eGFP fluorescence intensity of each cell was determined in the B525/40-A channel and normalized to forward-scatter area (FSC-A) to account for differences in cell size. Boxplots show the statistics including the arithmetic mean (x) of ∼ 30,000 cells per strain analyzed in three independent experiments. Range of the whiskers indicate population variability referring to the extent of dispersion around the median.

### 2.9. UstiGate-enabled multigene expression

Finally, to demonstrate the efficient assembly of four TUs as well as their straightforward, simultaneous genomic integration in *U. maydis*, we aimed to generate a “rainbow” strain producing four FPs with distinct emission maxima as an easy read-out. To this end, we combined the CDS of eGFP, mKate2, LSSmOrange and mTagBFP2 in Lvl1 with four different promoters and terminators from our toolbox to minimize repetitive sequences. Subsequently, five different final LvlT plasmids were assembled using the Lvl1 parts of the *car2* flanking regions and a resistance cassette. Each construct comprised either one of the TUs individually to generate control strains for each FP or all four TUs together to create the “rainbow” strain (Fig. 10A). In the single-TU constructs, the corresponding dummies (Supplementary Fig. S1) were applied for LvlT positions that were not filled with TUs. The fluorescence spectra of the rainbow strain show the expected emission maxima of all four FPs and exhibit largely comparable fluorescence intensities to the strains in which the FPs were integrated individually (Fig. 10B). Only the emission intensity of the blue FP is markedly reduced in the “rainbow” strain, which was further confirmed by flow cytometry (Supplementary Fig. S15). This effect may be explained by Förster resonance energy transfer (FRET) between FPs in close proximity within the cytoplasm of the “rainbow” strain. According to the FPbase FRET Calculator (https://www.fpbase.org/fret/) mTagBFP2, an efficient FRET donor ^95^, is predicted to transfer energy to eGFP or LSSmOrange with both donor-acceptor pairs exhibiting a Förster radius of approx. 5 nm. Nevertheless, all four FPs were detectable at the single-cell level and by fluorescence microscopy (Fig. 10C), confirming the successful simultaneous integration of four TUs within a single construct and in a single transformation step. Hence, UstiGate significantly speeds up genetic engineering in *U. maydis*, paving the way for efficient pathway transplantation and metabolic engineering with up to four modifications at a time.

**Fig. 10:**
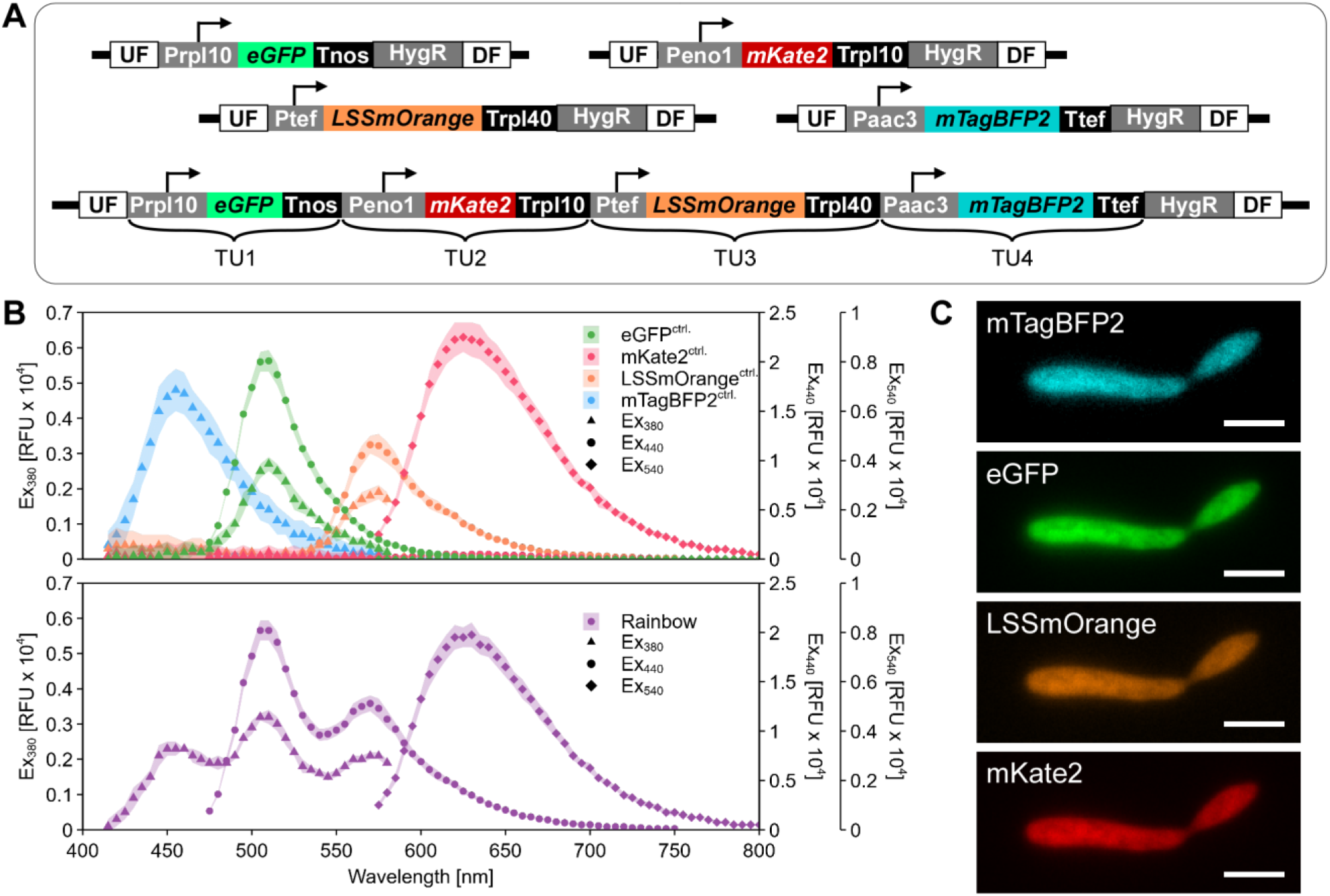
Generation of a rainbow strain demonstrates simultaneous integration of four TUs. **A)** Genomic architecture of the strains harboring one fluorescent protein (control strains) and the “rainbow” strain with four transcriptional units (TU1 - 4) for expression of the four fluorescent proteins (eGFP, mKate2, LSSmOrange, mTagBFP2). All constructs contain the flanking regions of the *car2* locus (UF, DF) and the hygromycin resistance cassette (HygR). All TUs were driven by different promoters of comparable strength (Prpl10, Peno1, Ptef, Paac3) and terminators were also varied (Tnos, Trpl10, Trpl40, Ttef) to avoid repeated sequences in the multigene construct. **B)** Fluorescence spectra of the control (^ctrl.^) strains (top) and the “rainbow” strain (bottom), measured with a gain setting of 150. Excitation wavelengths of 380 nm, 440 nm and 540 nm were applied for the spectra of mTagBFP2, eGFP/LSSmOrange and mKate2, respectively. Arrows depict emission peaks of the four fluorescent proteins. The spectra show arithmetic mean and standard deviation of biological triplicates across independent experiments. **C)** Fluorescence micrographs depicting a single “rainbow” cell. Different, appropriate excitation wavelengths and emission filters were applied for mTagBFP2, eGFP, LSSmOrange and mKate2 detection. Scale bars, 5 µm.

## 3. Conclusions

In the present study we improved genetic manipulation for synthetic biology approaches in the fungal basidiomycete model *Ustilago maydis*. To this end, we implemented a systematic modular cloning system based on the Golden Gate principle that is fully compatible with the established MoClo syntax from the plant and cyanobacterial field. Importantly, while adopting the overall structure we also established novel organism-specific elements like special level T vectors, constituting short-cuts for efficient targeting of distinct genomic loci. Beyond standard applications like gene deletion, the system supports the efficient generation of multigene expression constructs with up to four transcriptional units, which can be implemented in the fungal genome in a single step. As a basis for flexible applications, our system provides the community with a collection of 92 basic parts and vectors, including over 20 different promoters, a selection of terminators, resistance cassettes and fluorescent proteins, as well as special level T plasmids for several genomic loci and additional dummy vectors (Supplementary Fig. S16; Addgene kit deposit in progress). The promoters were characterized at the single-cell level as well as via continuous monitoring in different media. Moreover, effects of insertion locus, terminator and reporter were evaluated and a new auto-induction promoter based on maltose was implemented. Besides an improved cloning procedure and an expanded molecular toolbox, we also present a new locus for genomic insertion of constructs based on a color-screen, further streamlining genetic engineering. Together, this toolkit pushes *U. maydis* research further, enabling even the transplantation of complex heterologous pathways with significantly reduced workload. In addition, we expect that the toolbox is easily transferable to related basidiomycete fungi of interest ranging from closely related smut fungi like *Sporisorium reilianum*, *Ustilago hordei* or *Ustilago cynodontis* ^111–113^ to basidiomycetous human pathogens like *Cryptococcus* spp. or biotech workhorses like *Pseudozyma* spp. ^114,115^.

## 4. Methods

### 4.1. Gene accession numbers

The genes mentioned in this study have the following accession numbers: *car2* (*UMAG_06287*); *car1* (*UMAG_04210*); *cco1* (*UMAG_00965*); *ima1* (*UMAG_15026*); *upp3* (*UMAG_11908*); *pep4* (*UMAG_04926*); *ip* (*UMAG_00844*). The accession numbers of downstream or upstream genes of characterized promoters and terminators are listed in Supplementary Table S1 Supplementary Table S3.

### 4.2. Molecular biology methods

All plasmids utilized or generated in this study are listed in Supplementary Table S4. The UstiGate system uses acceptor vectors, dummies and linkers from the Golden Gate plant MoClo toolkit and from the CyanoGate toolbox. Both can be acquired from Addgene (kit no. 1000000044; ^55^ and kit no. 1000000146; ^68^). All generated plasmids were obtained using Golden Gate ^52^ and the new UstiGate standard described in the results section (see 2.1). The Addgene deposit of the UstiGate kit is currently in progress. Utilized enzymes were purchased from NEB (Ipswich, MA, USA).

For the domestication of genetic elements, specific primer overhangs were used depending on the genetic element and the intended level 0 (Lvl0) acceptor vector (Table 1). In addition to the BbsI sites corresponding to the MoClo syntax, SspI and SfiI recognition sites were preserved from previous cloning strategies and incorporated during domestication of up- and downstream flanks. While SspI sites are required for the generation of linear transformation constructs with homologous blunt ends ^26^, SfiI sites enable rapid exchange of genetic elements between the flanking regions across vectors ^28^.

**Table 1:**
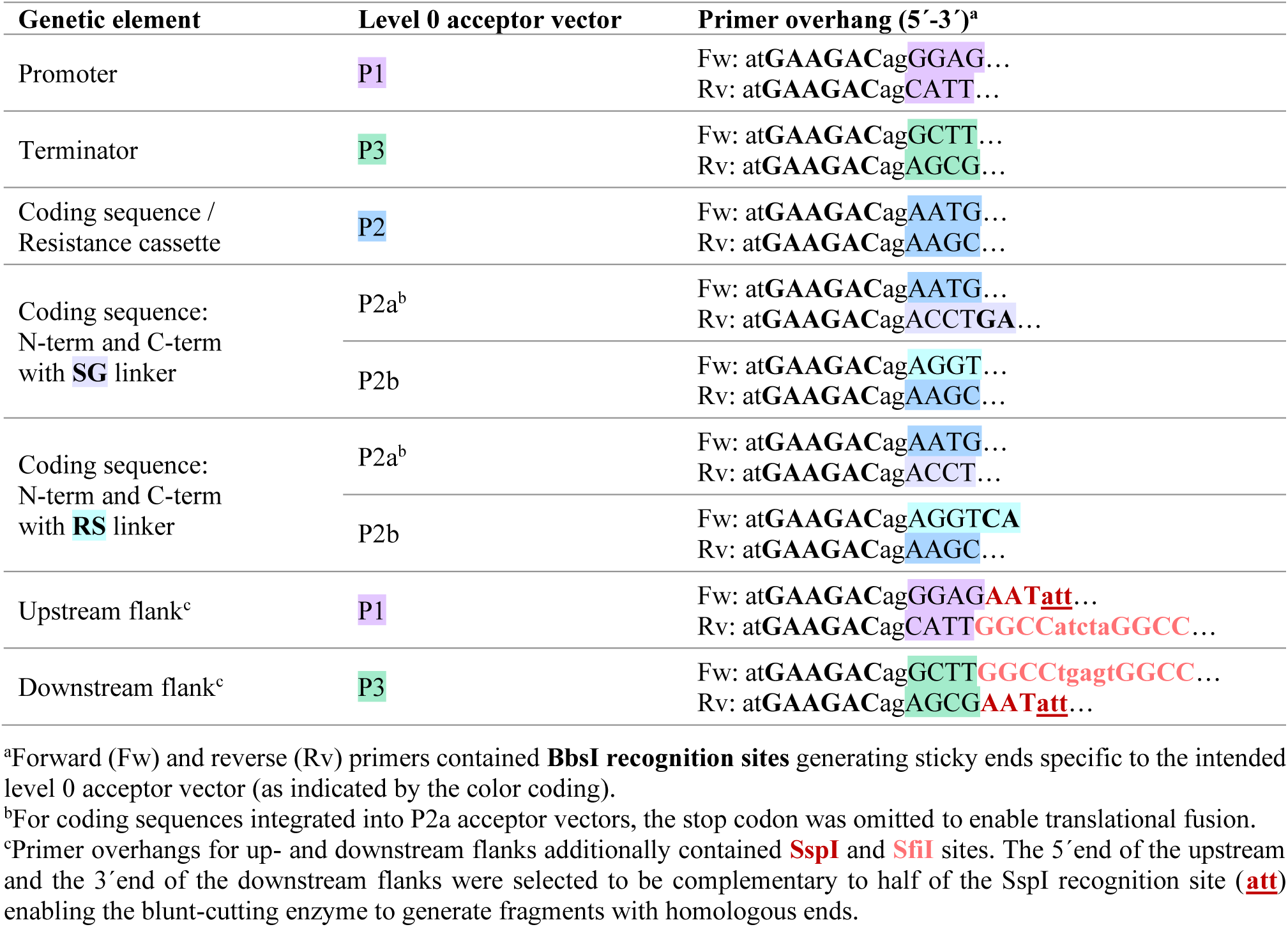
Primer overhangs used for the integration of different genetic elements into level 0 acceptor vectors.

Removal of internal BbsI, BsaI, SspI and SfiI recognition sites during integration of genetic elements into Lvl0 acceptor vectors was achieved by primer-mediated mutagenesis. Two oppositely-oriented primers introduced the desired point mutation within an exact 4 bp overlapping region and contained the standard “atGAAGACag” overhang to add BbsI recognition sites for subsequent Golden Gate assembly. Using this approach, genetic elements split into up to three fragments were routinely assembled. When multiple point mutations within a region of approx. 100 bp were required, one mutation was introduced within the described 4 bp overlap, while additional mutations were incorporated through modifications in an extended primer sequence of up to 120 bp in length. Within coding sequences, silent point mutations were introduced to not alter the encoded amino acid sequence. For native *U. maydis* sequences, genomic DNA of *U. maydis* strain UM521 was used as template for PCR reactions ^116^. The genomic sequence for this strain is stored at the EnsemblFungi database (https://fungi.ensembl.org/Ustilago_maydis/Info/Index; note that *U. maydis* is also known as *Mycosarcoma maydis*). Purified PCR products were assembled into the suitable Lvl0 acceptor vector in a Golden Gate reaction using 50 ng acceptor vector and a molar vector:insert ratio of 1:5. Two standard Golden Gate protocols were used depending on the required type IIs restriction enzyme, as BbsI-HF exhibits higher efficiency in combination with the NEBridge® Ligase Master Mix than with the T4 DNA Ligase Buffer (Supplementary Table S5). The reactions were performed in a PCR cycler selecting either a short (2.75 h) or a long (6.5 h) Golden Gate program (Supplementary Table S6) based on the complexity (number of fragments) of the assembly and the overall experimental schedule. For the generation of special level T (LvlT*) plasmids, 0.5 µL of fresh BbsI were added to the reaction after the Golden Gate program, and the mixture was incubated at 37 °C for at least 30 min before transformation into *E. coli*. Based on the expected efficiency, chemically competent *E. coli* Top10 cells (Invitrogen/Life Technologies) were transformed with 2-10 µL of the Golden Gate reaction. Selection plates contained 100 µg mL^-1^ ampicillin or spectinomycin and 50 µg mL^-1^ X-Gal. Resulting colonies were screened using PCR and correct candidates were verified via sequencing.

### 4.3. Strain generation

All *U. maydis* strains used or generated in the present study are listed in Supplementary Table S7. Strains were generated by homologous recombination yielding genetically stable derivatives of the laboratory strain AB33 ^41^. To this end, protoplasts were generated and transformed with SspI-linearized plasmids using established protocols ^25^. All strains were verified by Southern blot analysis and stored in glycerol stocks at −70°C ^25^.

### 4.4. Cultivation

*U. maydis* strains were streaked from glycerol stocks onto complete medium (CM, ^117^) agar plates containing 10 g L^-1^ glucose and 20 g L^-1^ agar and incubated at 28 °C for 1 - 2 days. Liquid cultures were generally cultivated in glass tubes and incubated at 28 °C on a rotating wheel. Precultures consisting of 3 mL CM supplemented with 10 g L^-1^ glucose were inoculated from freshly prepared plates and grown overnight. Main cultures containing 3 mL Verduyn mineral medium (adapted from Verduyn et al. (1992) ^118^: 0.5 g KH_2_PO_4_, 5 g (NH_4_)_2_SO_4_, 0.2 g MgSO_4_ x 7 H_2_O, 0.01 g FeCl_3_ x 6 H_2_O per liter dH_2_O buffered with 0.2 M MES pH 7, supplemented with 1 mL L^-1^ trace element solution and filter-sterilized (0.22 µm filter). Trace element solution: 15 g EDTA (TitriplexIII), 4.5 g ZnSO_4_ x 7 H_2_O, 0.84 g MnCl_2_ x 2 H_2_O, 0.3 g CoCl_2_ x 6 H_2_O, 0.3 g CuSO_4_ x 5 H_2_O, 0.4 g Na_2_MoO_4_ x 2 H_2_O, 4.5 g CaCl_2_ x 2 H_2_O, 1 g H_3_BO_3_, 0.1 g KI and 3 g FeSO_4_ x 7 H_2_O per liter dH_2_O) with 10 g L^-1^ glucose were then inoculated to a starting optical density at 600 nm (OD_600_) of 0.04 using a respective volume of the precultures. Following this procedure, the main cultures reached an OD_600_ of approx. 1 after 18 h of cultivation and were subsequently used for BioLector online growth and fluorescence monitoring or plate reader- and cytometry-based fluorescence measurements.

### 4.5. BioLector online growth and fluorescence monitoring

For experiments conducted in the BioLector I microbioreactor (m2p-labs/Beckman Coulter), round 48-well plates (MTP-R48-B; m2p-labs/Beckman Coulter) were used with a total working volume of approx. 1.5 mL per well. Initially, 1.3 mL of medium were dispensed into each well. Depending on the experiment, either CM, Verduyn medium containing 10 or 20 g L^-1^ of different carbon sources (glucose, sucrose, maltose, or arabinose) or a modified Verduyn medium with reduced nitrogen content and excess glucose (0.5 g (NH_4_)_2_SO_4_ and 50 g L^-1^ glucose) to cause nitrogen-limitation was used. For each strain included in the experiment, an equal volume (typically 150 µL) of the prepared main cultures (see above) were added to the respective wells resulting in an initial OD_600_ of approx. 0.1. The plate was sealed using a sterile, air-permeable adhesive film (VWR, European Article No 391-1262) and incubated in the BioLector at 28°C and 1,000 rpm (orbital shaking diameter: 3 mm) for approx. 3 days. For the addition of sugars or nitrogen during the BioLector experiment, the cultivation was briefly interrupted and 60 µL of sterile dH_2_O, (NH_4_)_2_SO_4_ solution (25 g L^-1^), or glucose, maltose or arabinose solution (250 g L^-1^) were added to the respective wells. Biomass formation was monitored by measuring scattered light using a 620 nm filter with a bandwidth of 10 nm (620/10 nm) and a gain of 20. eGFP fluorescence was measured using a filter module with excitation (Ex) at 480/10 nm and emission (Em) at 520/25 nm. Depending on the fluorescence intensity, a gain of 60 or 70 was selected for data analysis. Both parameters, biomass and fluorescence, were continuously monitored with a measurement interval of 30 min. For the analysis of the resulting data, the lowest starting value of each dataset was used as a blank and subtracted from all other values prior to calculating the arithmetic mean and deviation between biological replicates across independent BioLector experiments.

### 4.6. Plate reader-based bulk fluorescence measurements

Fluorescence intensities of reporter strains with cytosolic fluorescence proteins were measured both as bulk fluorescence using a Tecan Infinite 200 Pro plate reader and at the single-cell level using flow cytometry. For this purpose, prepared main cultures (see above) were diluted in a black-walled, clear-bottom 96-well plate (Greiner 655906) to an OD_600_ of approx. 0.5 - 0.8 in a total volume of 200 µL per well. Fluorescence intensities of the reporter strains were measured during the exponential growth phase at an OD_600_ of approx. 1 and again in the stationary growth phase after an additional 24 h of cultivation. During the exponential growth phase, cultures were diluted 1:2 using phosphate buffered saline (PBS), whereas a 1:10 dilution was required during the stationary growth phase to avoid high abort rates during flow cytometry.

In the Tecan plate reader, absorbance was measured at 600/9 nm (OD_600_), and fluorescence was measured using the near-optimal Ex and Em wavelengths for the different FPs (Supplementary Table S2). For the fluorescence spectra of the “rainbow” strain, Ex wavelengths more distant from the corresponding Em ranges were selected (eGFP, LSSmOrange: Ex 440/10 nm, Em 475-750/20 nm; mKate2: Ex 540/10 nm, Em 570-800/20 nm; mTagBFP2: Ex 380/10 nm, Em 410-580/20 nm). Em was measured every 5 nm across the indicated wavelength ranges. A gain between 100 and 150 was used for all Tecan fluorescence measurements. For the analysis of the resulting data, fluorescence values were first normalized to the corresponding OD_600_, and then the background fluorescence of cells without an FP was subtracted prior to calculating the arithmetic mean and deviation between biological triplicates across independent experiments.

### 4.7. Flow cytometry-based fluorescence measurements at the single-cell level

After the bulk fluorescence measurements, the same 96-well plate was subsequently used for flow cytometry analysis of the reporter strains using a CytoFLEX S Flow Cytometer (Beckman Coulter, Model No.: B75442, SN: BE51180). The cytometer was operated in auto acquisition plate mode with a 5 s backflush between samples and 5 s mixing of each sample. Appropriate laser and detector combinations were selected for the different FPs. Letters thereby describe the Ex laser (V=Violet: 405 nm; B=Blue: 488 nm; Y=Yellow: 561 nm), while numbers describe the Em wavelength and bandwidth of the filters with the applied gains given in parentheses: eGFP: B525/40 (100); mKate2: Y610/20 (500); LSSmOrange V585/15 (1500); mTagBFP2: V450/45 (500). The gain for forward scatter (FSC) was set to 25 and the threshold for detection was set to 2,000 in the FSC-Height channel. For each sample, 10,000 events were recorded with a sample flow rate of 10 µL min^-1^. For data analysis of the cytometry data, gating was performed in the FSC-Area channel, which is correlating with the cell size, defining all events >10^4^ arbitrary units (AU) as *U. maydis* cells, leaving roughly 9,900 cells per sample. Calculations were then performed for the *U. maydis* cells using the area values (-A) of the different channels. Fluorescent values of individual cells were normalized to FSC-A by division of the fluorescence value by the FSC-A value (e.g. B525/40-A / FSC-A) to account for differences in cell size prior to calculating boxplot statistics including all measured cells of the same strain across the biological triplicates from independent experiments.

### 4.8. Fluorescence microscopy

Fluorescence microscopic analysis was conducted with yeast-like cells grown to an OD_600_ of about 0.5 using an openFrame Microscopy System (Cairn GmbH, Ettlingen, Germany) equipped with the SCMOS sensor KINTEIX-22MM-M-C (Teledyne Photometrics, Tucson, AZ, USA) and an LDI-7 solid-state laser (89north, Williston, VT, USA) as illuminator. The system enabled the selection of appropriate Ex wavelengths and Em filters for the four different FPs present in the cells of the “rainbow” strain: eGFP (Ex 470 nm, Em 510/50 nm), mKate2 (Ex 555 nm, Em 630/75 nm), LSSmOrange (Ex 445 nm, Em 560/50 nm) and mTagBFP2 (Ex 405 nm, Em 480/40 nm). Image acquisition was performed using Micro-manager 2.0 ^119^. Image processing included the adjustments of brightness and contrast and was performed with ImageJ (software version 1.54g).

## 5. Author information

### 5.1. Corresponding author

Dr. Kerstin Schipper; Institute for Microbiology, Heinrich Heine University Düsseldorf, Universitätsstraße 1, 40225 Düsseldorf, Germany;

### 5.2. Author contribution

J.C.H. designed and performed experiments with support from V.A.P. J.C.H wrote the original draft, generated figures and tables, and revised the manuscript. K.S. directed the study, supported data evaluation and revised the manuscript. D.H. assisted in flow cytometry experiments, conducted cytometry raw data evaluation, and reviewed and edited the manuscript. I.M.A. reviewed and edited the final draft. K.S. and I.M.A. acquired the funding, and K.S. was responsible for project administration. All authors read and approved the final manuscript.

## 6. Associated content

### 6.1. Supporting information

Additional results and visualization of the UstiGate kit components (Fig. S1–S16), as well as additional details on utilized promoters, fluorescent proteins, terminators, plasmids, experimental procedure, and strains (Tables S1-7) (PDF)

### 6.2. Data availability statement

All data and materials are available upon request. Prior to publication, we will provide all experimental raw data in an Annotated Research Context (ARC) for FAIR data management. The Addgene kit deposit is currently in progress.

## 7. Acknowledgment

We are deeply grateful to Bettina Axler for the excellent technical support in plasmid and strain generation. We thank Johannes Postma for assistance with fluorescence microscopy, Tom Berwanger for plasmid generation, and Dr. Jungho Lee and Dr. Kai Hußnätter for valuable preliminary work and strain generation. We also appreciate Aykut Moumin’s help with plasmid and strain generation. We thank Prof. Dr. Sander Smits and Dr. Athanasios Papadopoulos from the Center for Structural Studies (CSS) at HHU as well as CRC1535 MibiNet project Z01 for providing cpT-Sapphire, LSSmApple and LSSmKate2. We thank Dr. Anna Behle for the domesticated coding sequences of LSSmOrange and mTagBFP2, as well as for her initial input regarding the promoter screen. We thank Dr. St. Elmo Wilken for providing the domesticated coding sequence of Electra2. We are very grateful to Dr. Sabrina Zander for her essential support in generating the annotated research context (ARC) and we thank Dr. Lesley Plücker for providing TPM values of the RNA-seq dataset of Lanver et al. (2018). We also thank Prof. Dr. Michael Feldbrügge for his general support and his valuable comments on the manuscript and Dr. Lesley Plücker and Dr. Florian Altegoer for critical review of the manuscript.

## 8. Funding

This research was generously supported by DFG-SFB1535 - project ID 458090666 to K.S., I.M.A. (project B03) and DFG Major Research Instrumentation INST 208/808-1.

Supplementary Material of the manuscript entitled

## Supplementary Material

**Supplementary Fig. S1:**
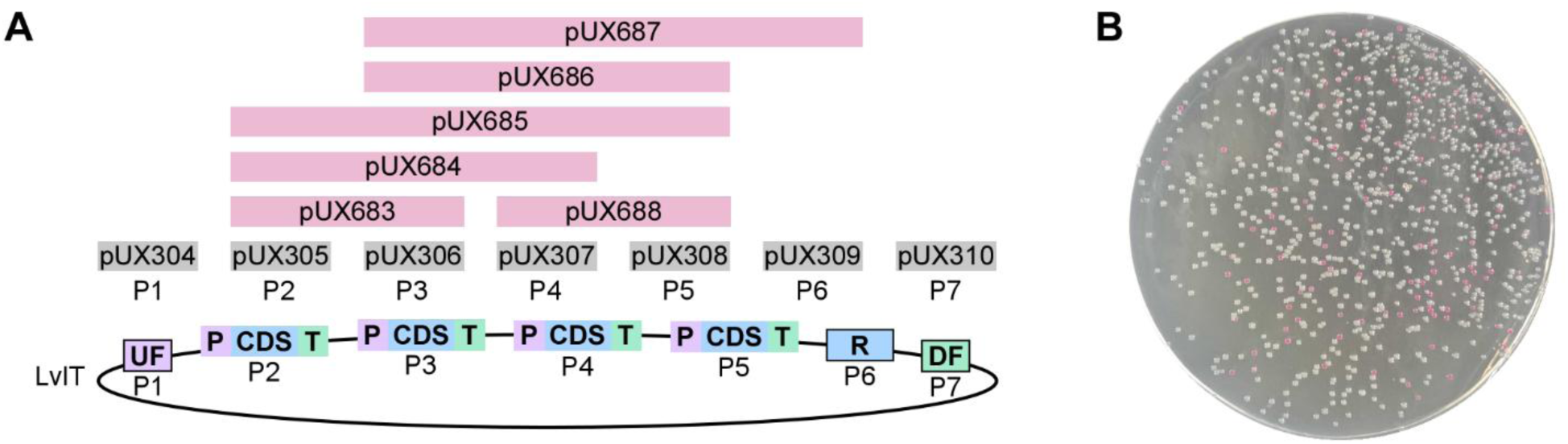
**A)** Available dummy fragments for replacing unused level 1 positions (P1-P7) during level T (LvlT) assembly. Dummies from the original MoClo toolkit ^grey; 55^ replace individual positions and newly designed dummies (pink) replace two to four consecutive positions. pUX, internal plasmid ID, see Supplementary Table S4; UF, DF, upstream- and downstream flank; R, resistance cassette; P, promoter; CDS, coding sequence; T, terminator. **B)** *E. coli* colonies harboring plasmids with (pink) and without (white) an mScarlet-I dropout cassette ^70^.

**Supplementary Fig. S2:**
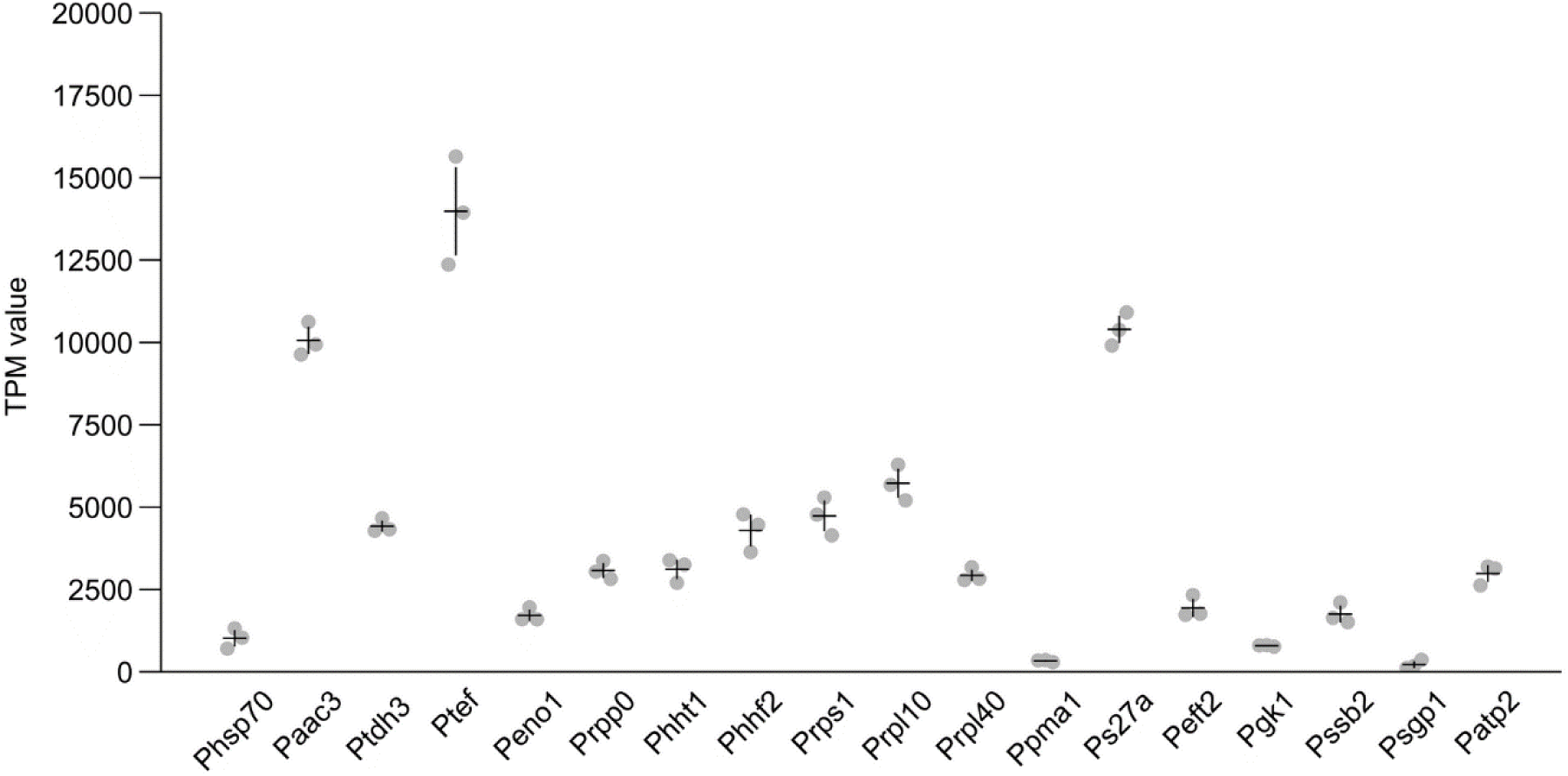
Expression levels of the native downstream genes driven by the indicated promoters, shown as transcripts per million (TPM). RNA-seq data from axenic *U. maydis* cultures in the exponential growth phase were taken from Lanver et al. (2018) ^79^. Each dot represents an individual biological replicate; black lines indicate the arithmetic mean ± standard deviation.

**Supplementary Fig. S3:**
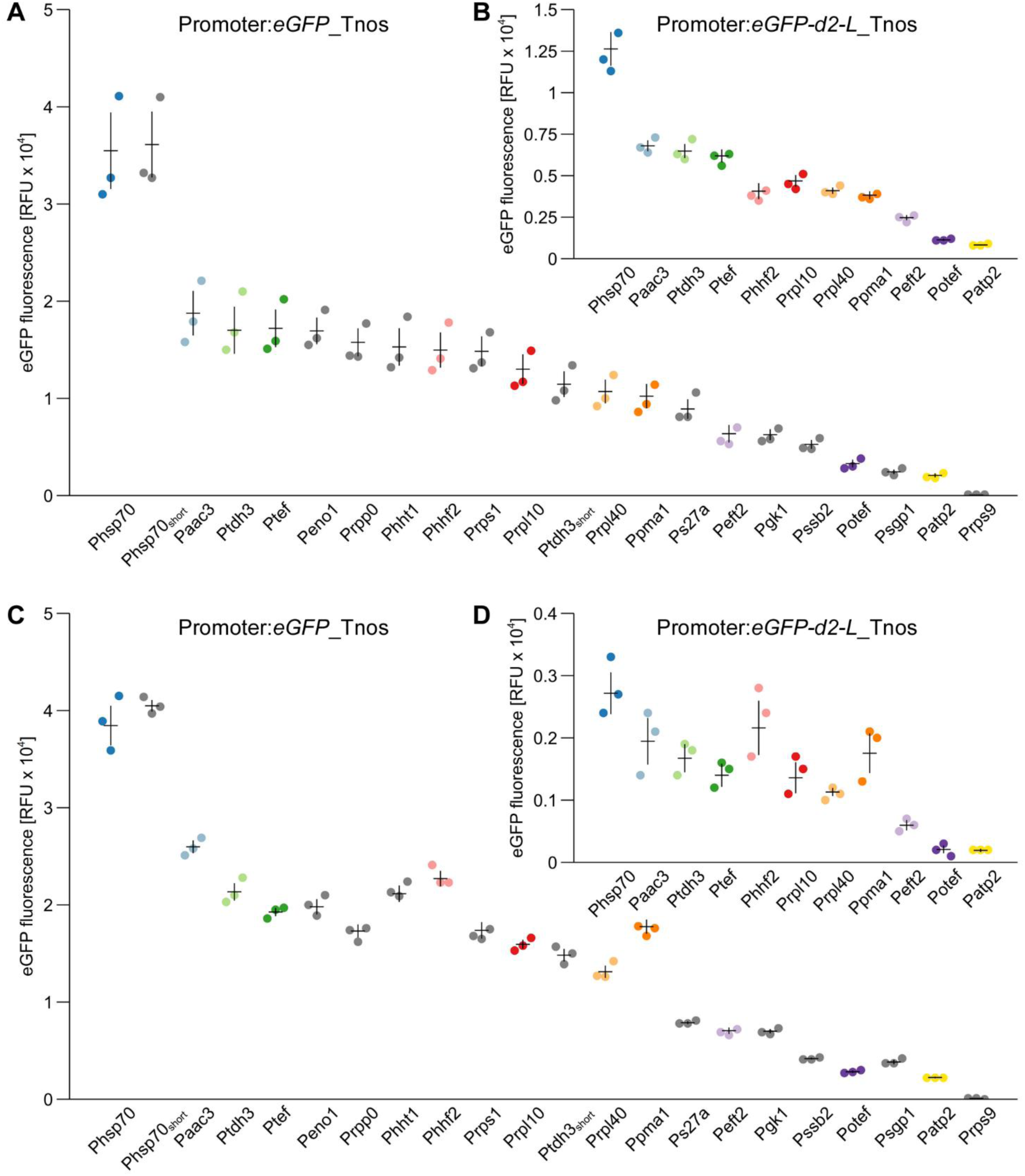
Bulk fluorescence measurements of promoter reporter strains performed in a plate reader. Strains were analyzed during the exponential (**A**, **B**) and the stationary (**C**, **D**) growth phase. eGFP fluorescence was measured at excitation and emission wavelengths of 480 nm and 515 nm, respectively, using a gain setting of 120. A subset of promoters (highlighted by colored dots in **A** and **C**) was additionally analyzed using the destabilized eGFP variant eGFP-d2-L (**B**, **D**). Each dot represents an individual biological replicate from independent experiments, and black lines indicate the arithmetic mean ± standard deviation. Note that the measurements were performed using the same culture material as used for the flow cytometry analyses shown in Fig. 4 and Supplementary Fig. S4.

**Supplementary Fig. S4:**
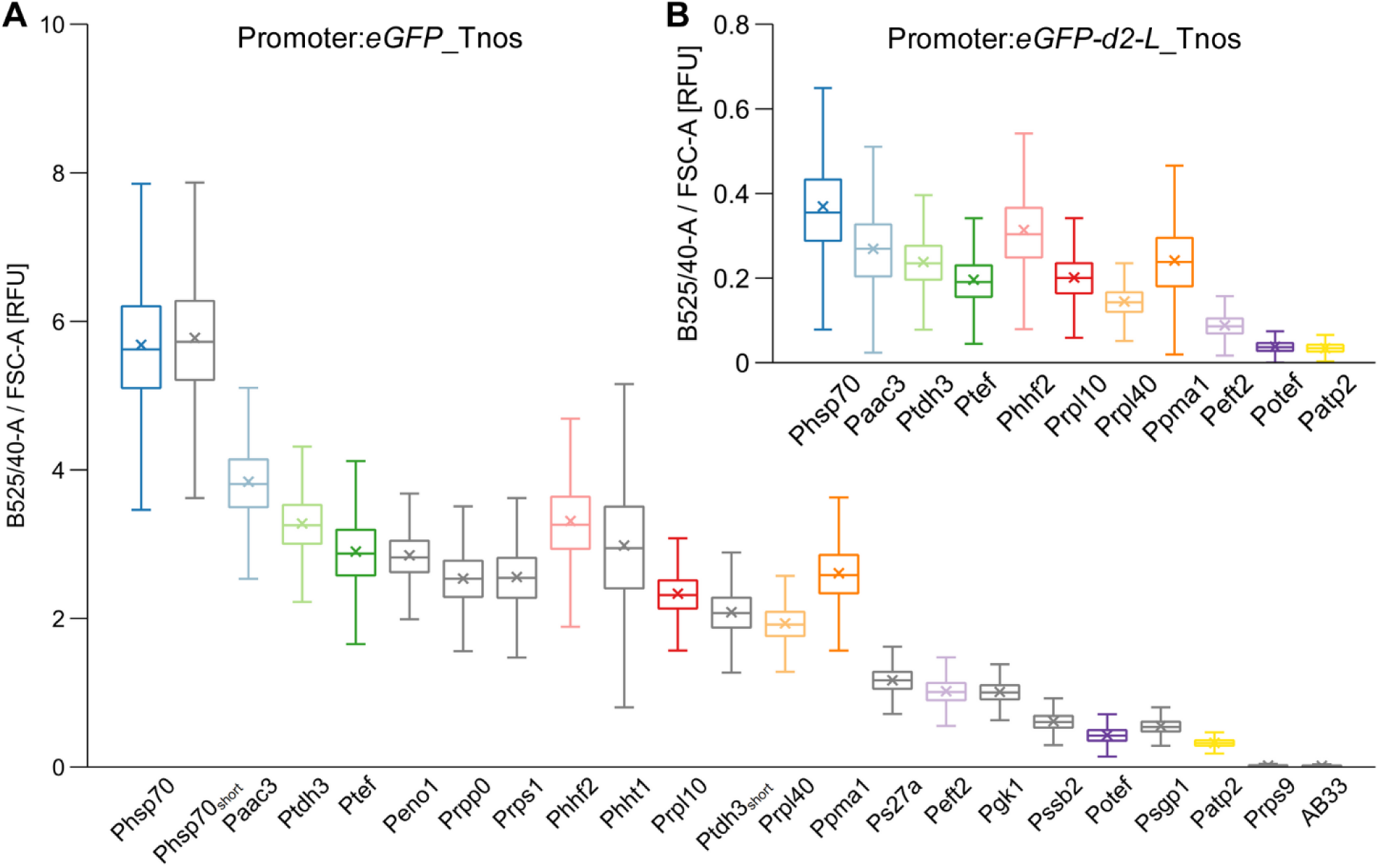
Comparison of different promoters at the single-cell level during the stationary growth phase. **A, B)** eGFP fluorescence intensity of each cell was determined using the B525/40-A channel of the flow cytometer and was normalized to forward-scatter area (FSC-A) to account for differences in cell size. Boxplots show the statistics including the arithmetic mean (x) of ∼ 30,000 cells per strain analyzed in three independent experiments. Range of the whiskers indicate population variability referring to the extent of dispersion around the median. **B)** Approx. half of the promoters (shown with colored boxplots in **A**) were additionally screened with the destabilized eGFP variant eGFP-d2-L. Note that the measurements were performed using the same cultures from which the exponential-phase data are shown in Fig. 4.

**Supplementary Fig. S5:**
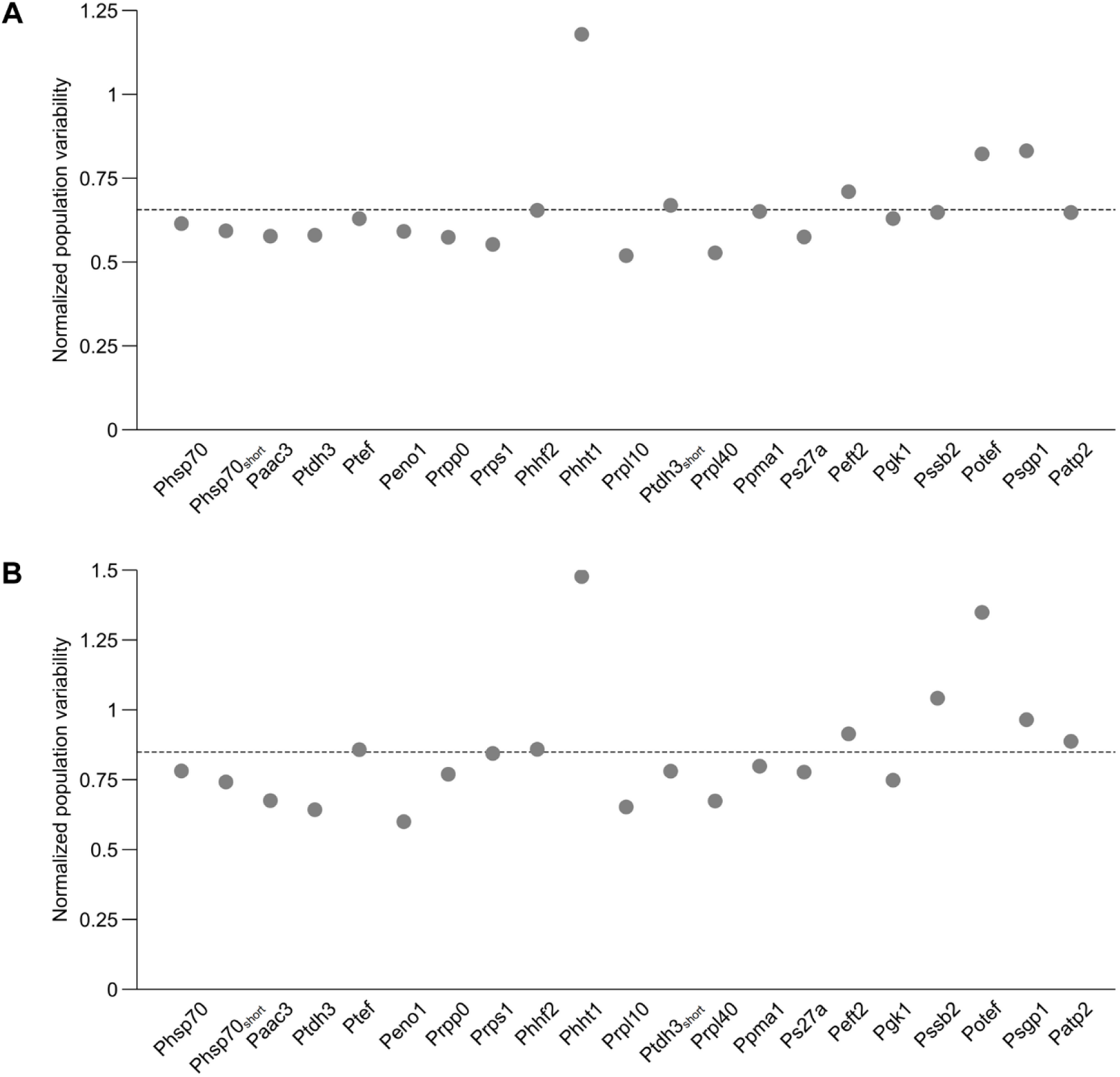
Normalized population variability of promoter reporter strains during the exponential (**A**) and stationary (**B**) growth phase. To account for the difference in overall fluorescence intensity, population dispersion was normalized by dividing the range of the boxplot whiskers (maximum minus minimum; Fig. 4B and Supplementary Fig. S4A) by the corresponding median value. The dotted line represents the arithmetic mean of the normalized population variability across all reporter strains shown.

**Supplementary Fig. S6:**
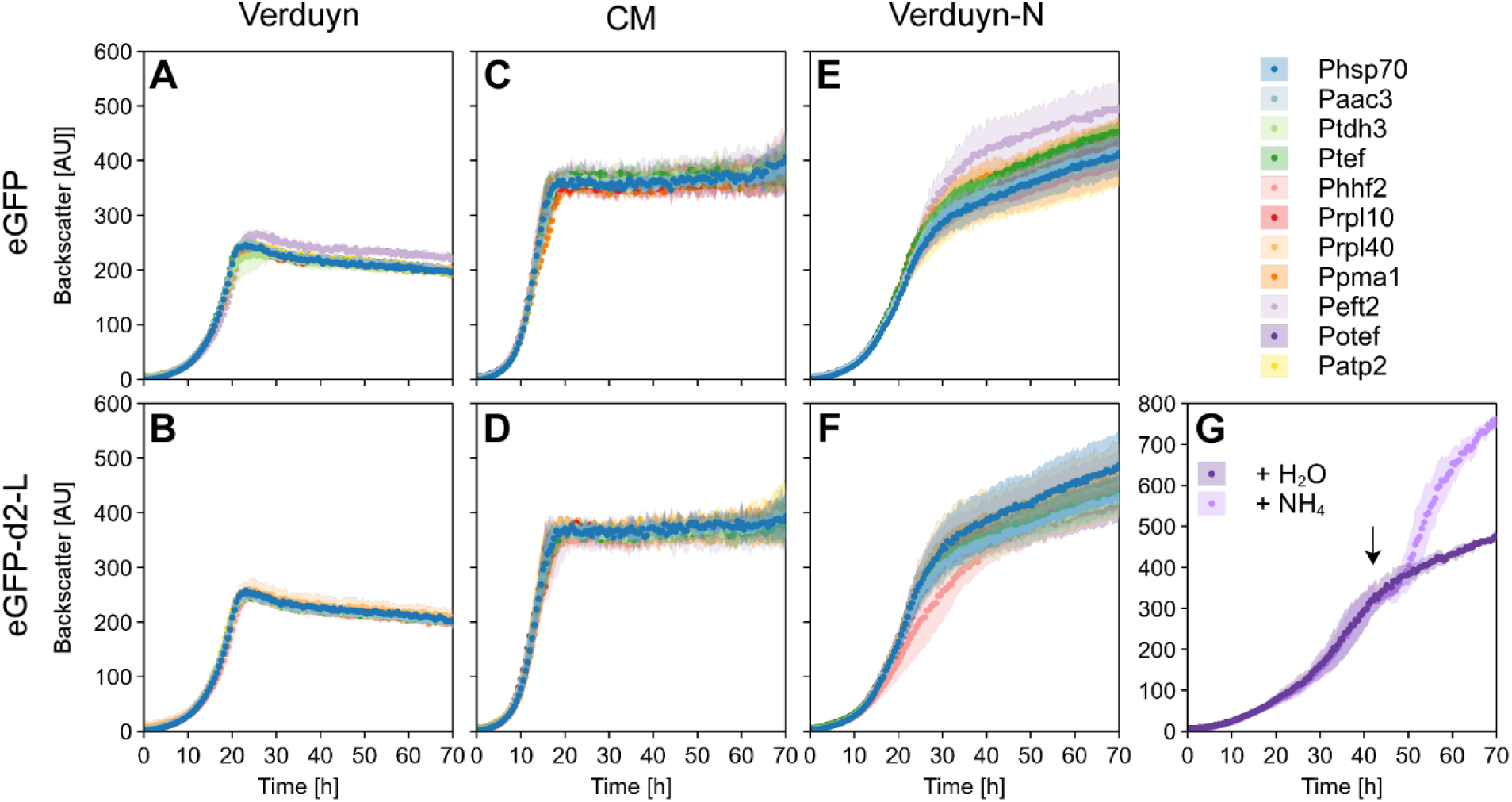
Online monitoring of growth behavior in different media. Strains producing either eGFP (**A**, **C**, **E**) or a destabilized version (eGFP-d2-L; **B**, **D**, **F**) under the control of different promoters were analyzed. Cells were grown in Verduyn mineral medium (**A**, **B**), complete medium (CM; **C**, **D**) or a modified Verduyn variant with excess glucose and reduced nitrogen content (Verduyn-N; **E**, **F**). Online monitoring was performed in a microbioreactor (BioLector) measuring biomass (backscatter; gain 20) and eGFP fluorescence (gain 70). Fluorescence data of the cultivations are provided in Fig. 5. **G**) Cultivation of the Potef:eGFP-d2-L strain in Verduyn-N medium with an ammonium spike introduced at about 42 h of cultivation (arrow). The graphs show arithmetic mean and standard deviation of biological triplicates (**A**, **B**, **E**, **F**) or duplicates (**C**, **D**, **G**) across independent experiments.

**Supplementary Fig. S7:**
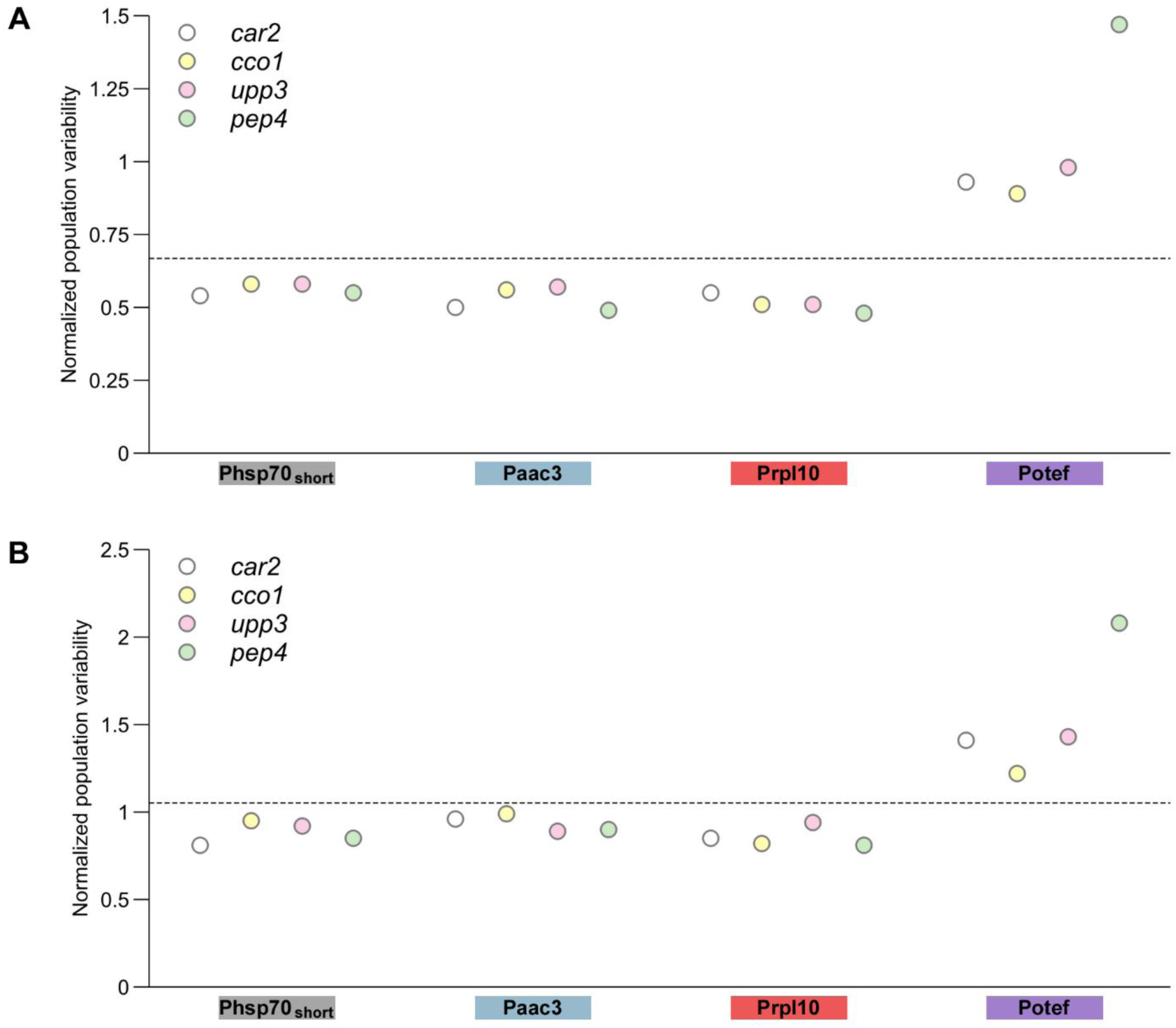
Normalized population variability of promoter reporter strains during the exponential (**A**) and stationary (**B**) growth phase. To account for the difference in overall fluorescence intensity, population dispersion was normalized by dividing the range of the boxplot whiskers (maximum minus minimum; Fig. 6 and Supplementary Fig. S8) by the corresponding median value. The dotted line represents the arithmetic mean of the normalized population variability across all reporter strains shown.

**Supplementary Fig. S8:**
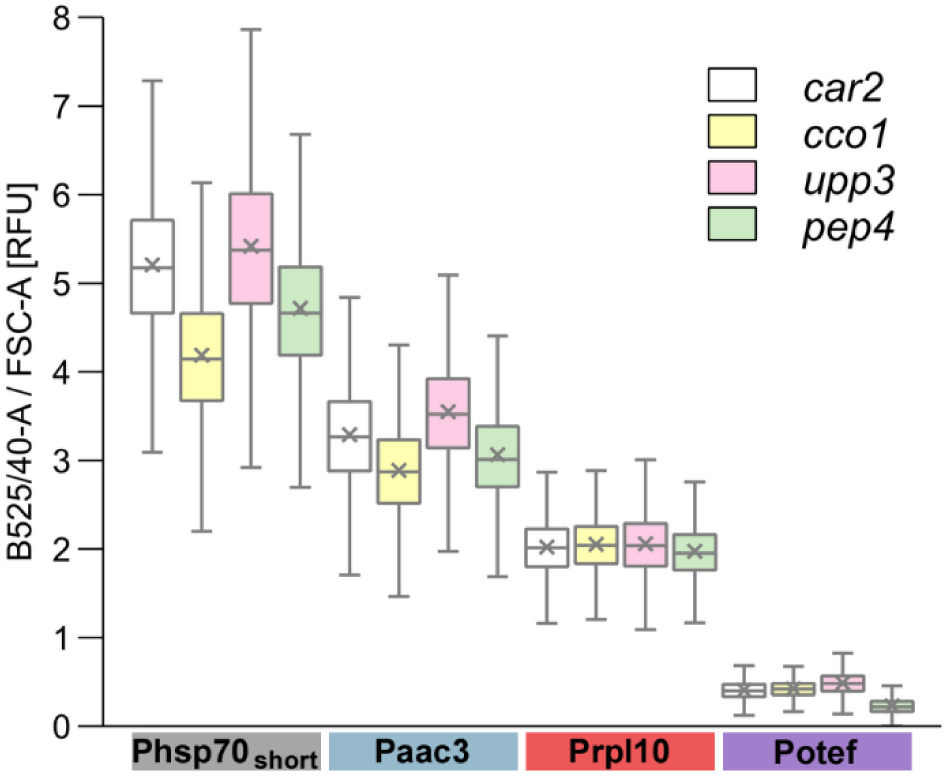
Comparative evaluation of different insertion loci in the stationary growth phase. Four promoters, Phsp70, Paac3, Prpl10 and Potef, were fused with *eGFP* and integrated into the *car2*, *cco1*, *upp3* and *pep4* locus, respectively. eGFP fluorescence intensity of each cell was determined using the B525/40-A channel of the flow cytometer and was normalized to forward-scatter area (FSC-A) to account for differences in cell size. Boxplots show the statistics including the arithmetic mean (x) of ∼ 30,000 cells per strain analyzed in three independent experiments. Range of the whiskers indicate population variability referring to the extent of dispersion around the median. Note that the measurements were performed using the same cultures from which the exponential-phase data are shown in Fig. 6.

**Supplementary Fig. S9:**
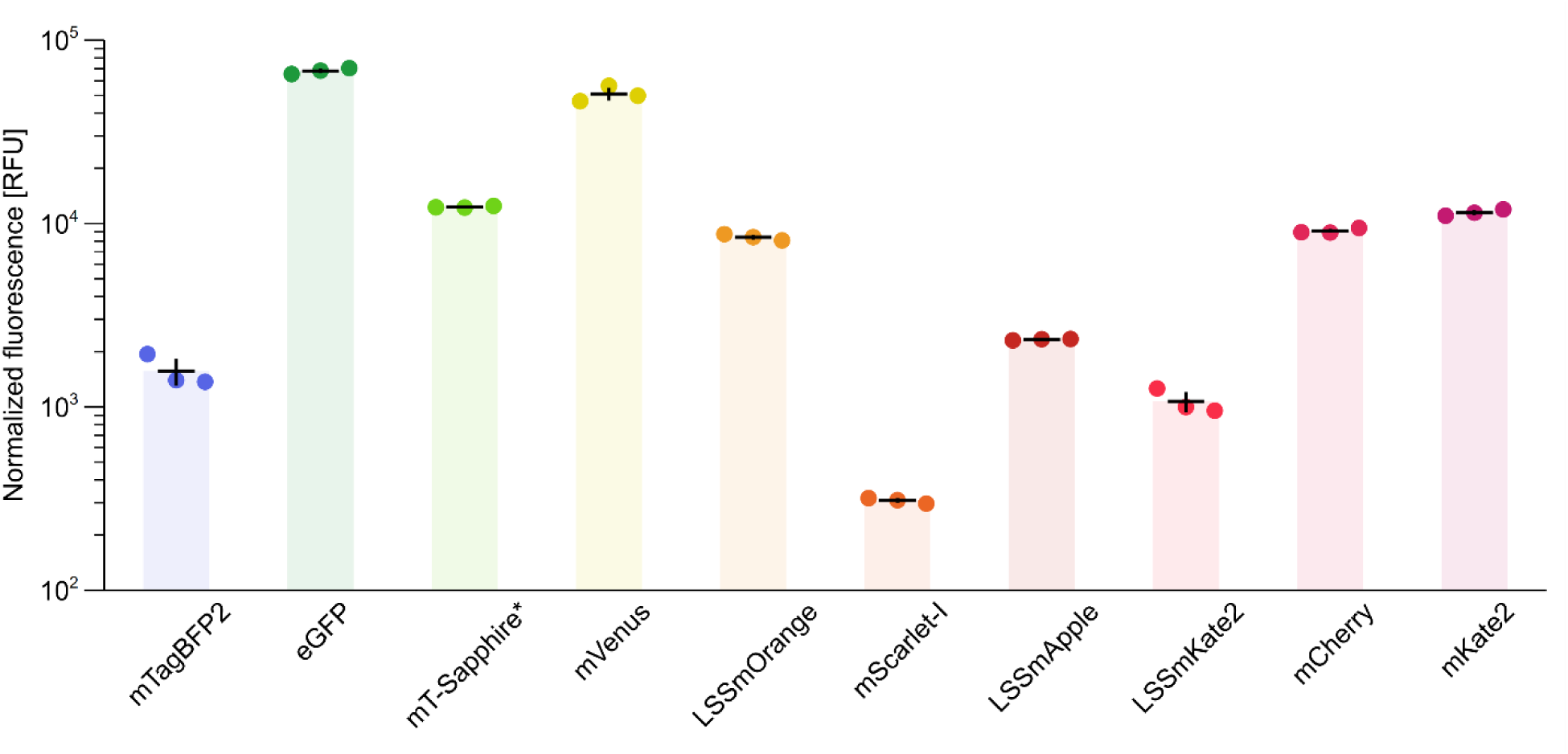
Comparative evaluation of the fluorescence intensity of ten fluorescent proteins (FPs) in *U. maydis*. FPs were evaluated by measuring bulk fluorescence of 1:5 diluted, stationary cultures in a plate reader. Fluorescence was measured at near-optimal excitation and emission wavelength for each FP (see Supplementary Table S2) while using a constant gain setting of 150. Fluorescence values were normalized to OD_600_ and the background fluorescence of cells without an FP was subtracted. Each dot represents an individual biological replicate from independent experiments, and black lines indicate the arithmetic mean ± standard deviation. mT-Sapphire*, mT-Sapphire with the following mutations: N147T S148A L232H. Note that the measurements were performed using the same cultures from which the exponential-phase data are shown in Fig. 7A.

**Supplementary Fig. S10:**
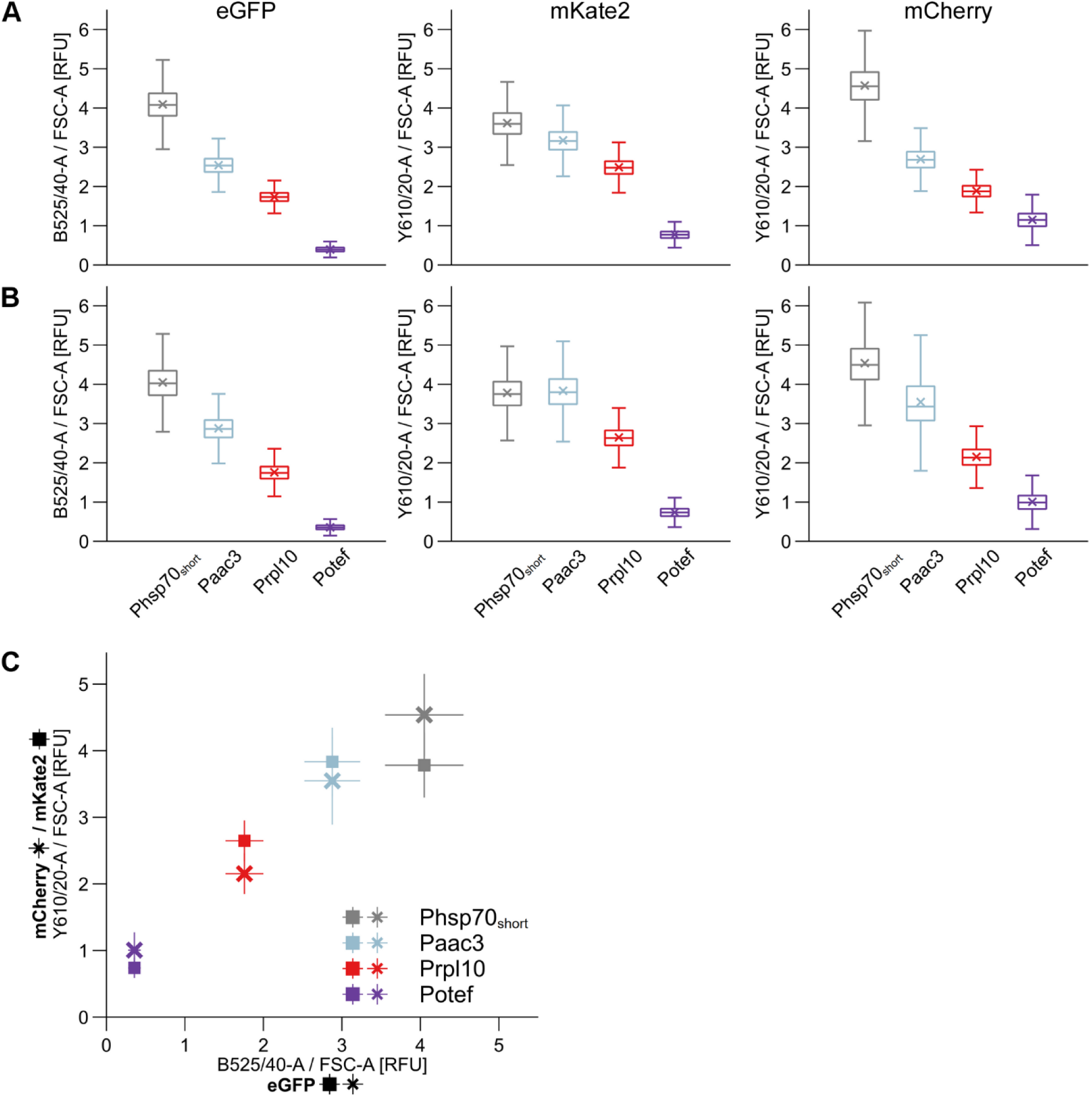
Comparison of three different fluorescent proteins as reporters for promoter activity in the exponential (**A**) and stationary (**B**, **C**) growth phase. Four promoters, Phsp70_short_, Paac3, Prpl10 and Potef, were each combined with *eGFP*, *mCherry* and *mKate2* and integrated into the *car2* locus. Fluorescence intensity of each cell was determined using the B525/40-A (eGFP) or the Y610/20-A (mCherry and mKate2) channel of the flow cytometer, and the fluorescence values were normalized to forward-scatter area (FSC-A) to account for differences in cell size. **A**, **B**) Boxplots show the statistics including the arithmetic mean (x) of ∼ 30,000 cells per strain analyzed in three independent experiments. Range of the whiskers indicate population variability referring to the extent of dispersion around the median. **C**) The arithmetic mean from **B** ± standard deviation is plotted with normalized fluorescence of the eGFP reporter strains on the x- and the corresponding values of the mCherry- or mKate2-producing strains on the y-axis. Note that the data were obtained using the same cultures from which the exponential-phase data are shown in Fig. 7D.

**Supplementary Fig. S11:**
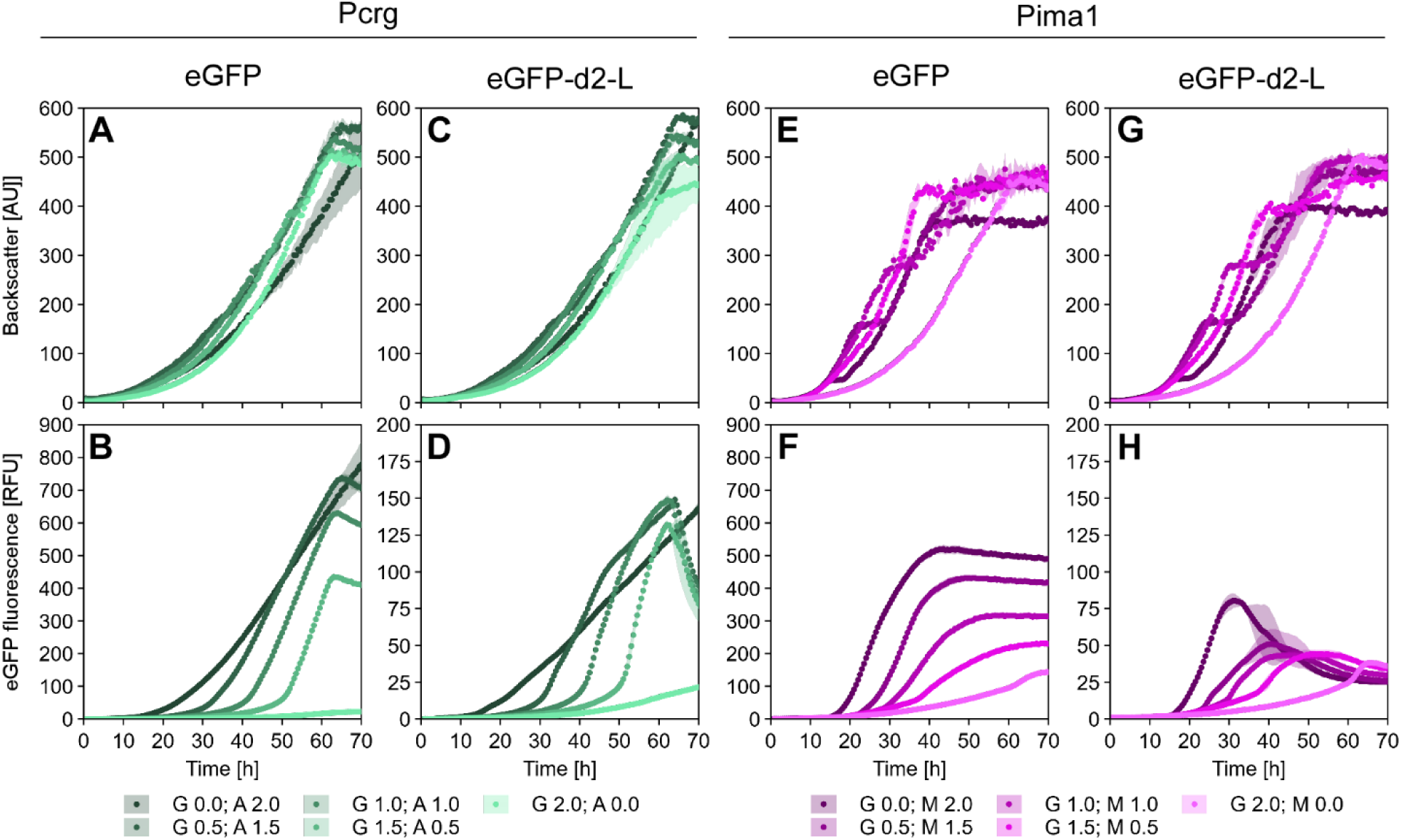
Online monitoring of growth (backscatter; **A**, **C**, **E**, **G**) and eGFP fluorescence (**B**, **D**, **F**, **H**) of Pcrg and Pima1 reporter strains with either eGFP or a destabilized version (eGFP-d2-L) as reporter. Cells were grown in Verduyn mineral medium supplemented with glucose (G 2.0; A/M 0.0) or the inducing sugars arabinose or maltose (G 0.0; A/M 2.0), as well as different ratios of both sugars. The graphs show arithmetic mean and standard deviation of biological duplicates across independent experiments. Note that **B** and **F** show the same data as shown in Fig. 8 C and D.

**Supplementary Fig. S12:**
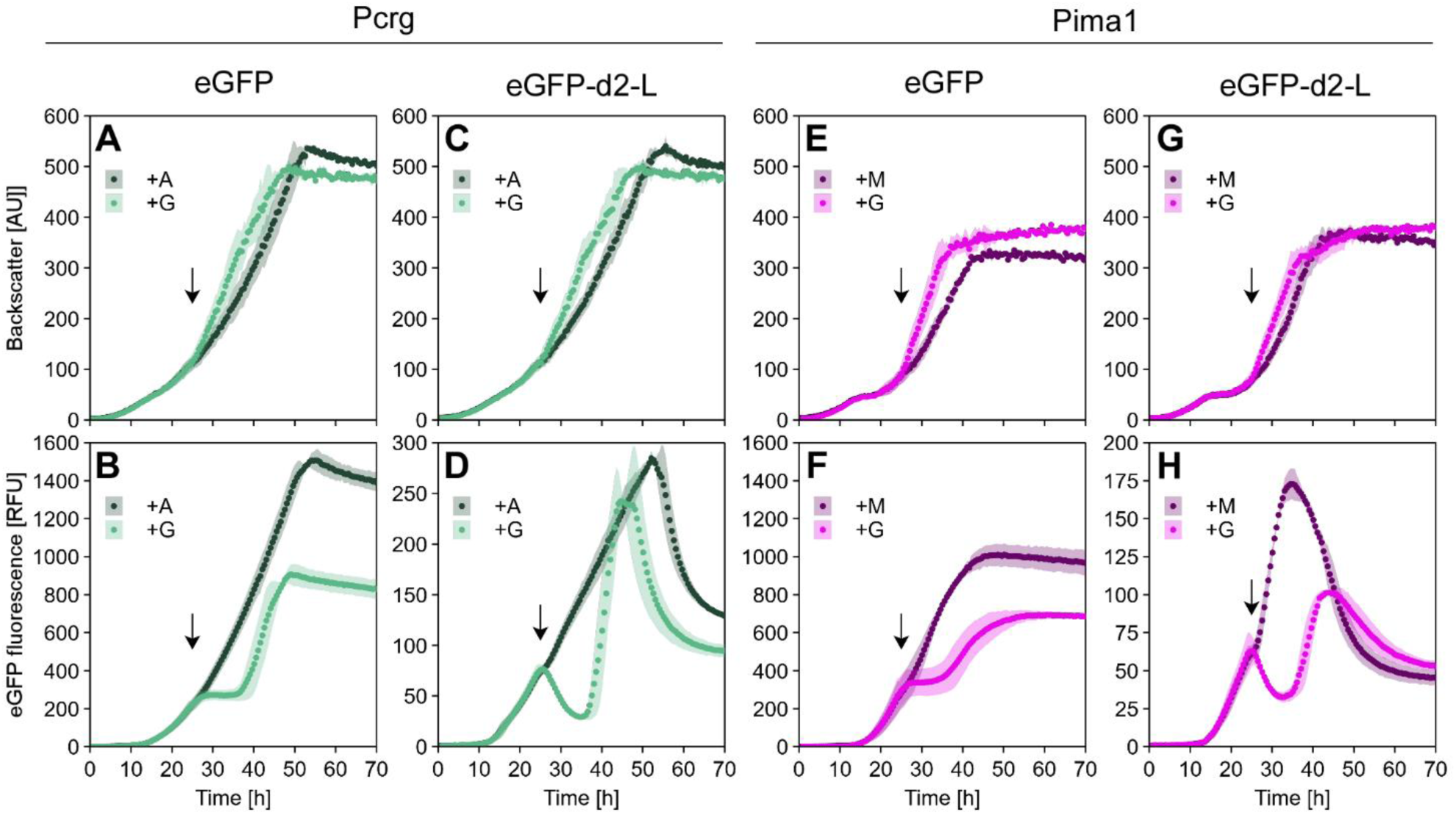
Online monitoring of growth (backscatter; **A**, **C**, **E**, **G**) and eGFP fluorescence (**B**, **D**, **F**, **H**) of Pcrg and Pima1 reporter strains with either eGFP or a destabilized version (eGFP-d2-L) as reporter. Cells were grown in Verduyn mineral medium supplemented with arabinose (Pcrg) or maltose (Pima1). At about 25 h of cultivation (arrow), either glucose (G) or more arabinose (A) or maltose (M) were added to the culture wells. The graphs show arithmetic mean and standard deviation of biological duplicates across independent experiments. Note that **D** and **H** show the same data as shown in Fig. 8 E and F.

**Supplementary Fig. S13:**
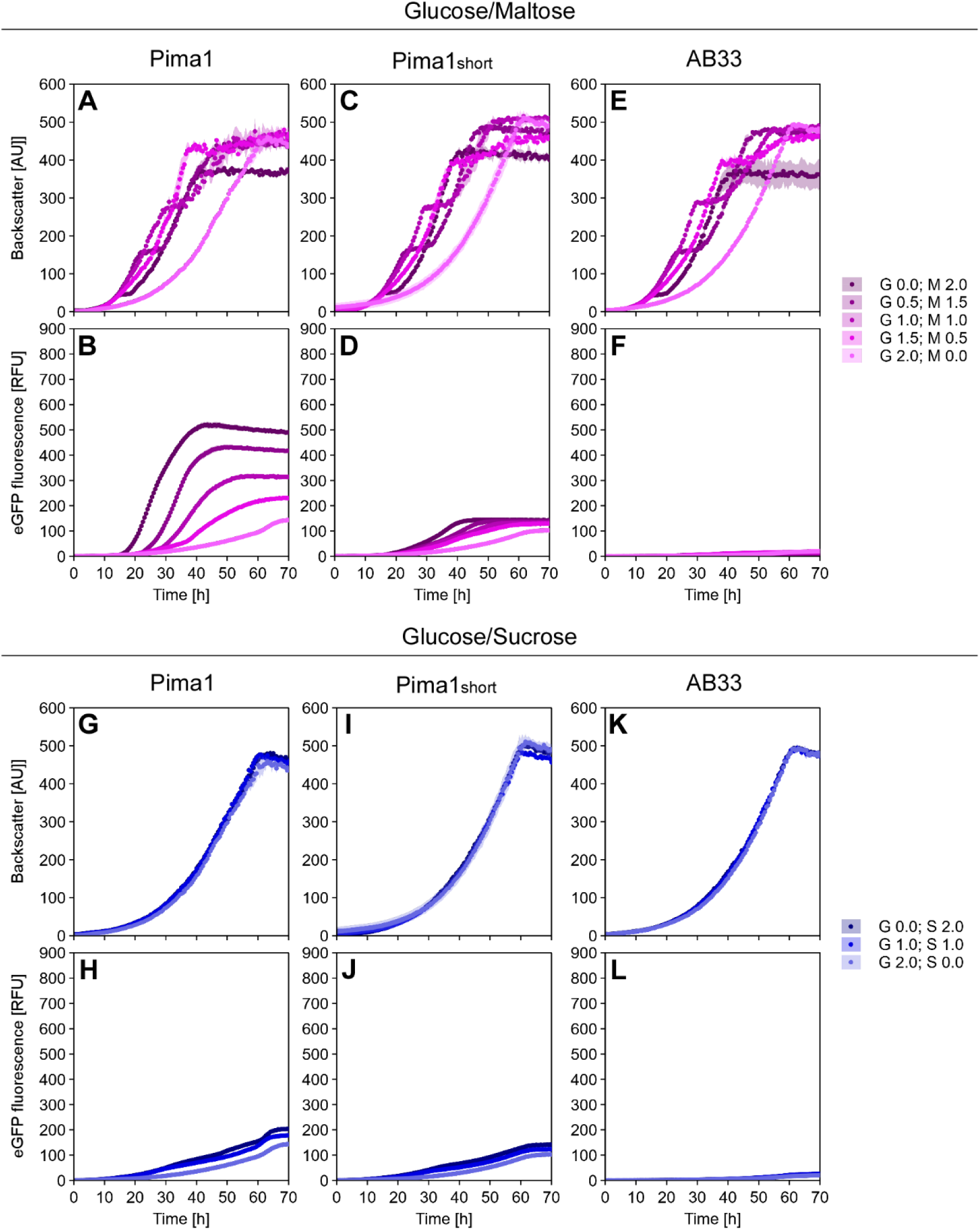
Online monitoring of growth (backscatter; **A**, **C**, **E**, **G**, **I**, **K**) and eGFP fluorescence (**B**, **D**, **F**, **H**, **J**, **L**) of the Pima1:eGFP and the Pima1_short_:eGFP reporter strains as well as their progenitor strain (AB33). Cells were grown in Verduyn mineral medium supplemented with glucose (G 2.0; A/M 0.0) or the potentially inducing sugars maltose or sucrose (G 0.0; M/S 2.0), as well as different ratios of both sugars. The graphs show arithmetic mean and standard deviation of biological duplicates across independent experiments. Note that **B** shows the same data as shown in Fig. 8D and Supplementary Fig. S11F.

**Supplementary Fig. S14:**
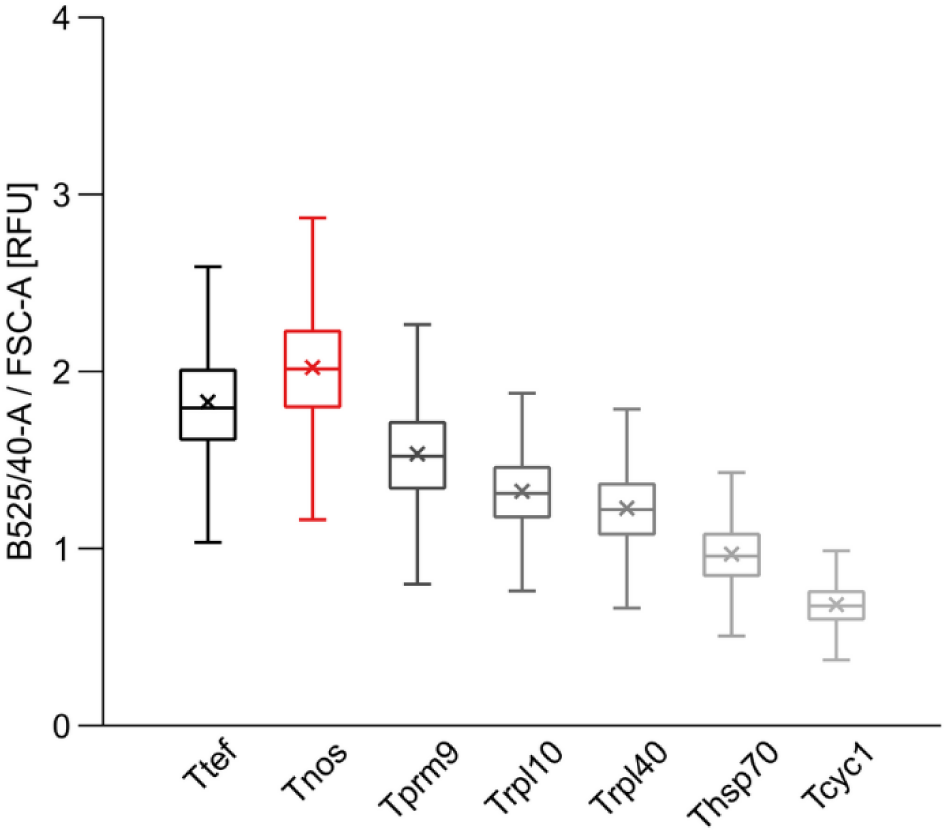
Comparative evaluation of different terminators in the stationary growth phase. The *rpl10* promoter was combined with *eGFP* and one of the seven terminators, respectively, and integrated into the *car2* locus. eGFP fluorescence intensity of each cell was determined using the B525/40-A channel of the flow cytometer and was normalized to forward-scatter area (FSC-A) to account for differences in cell size. Boxplots show the statistics including the arithmetic mean (x) of ∼ 30,000 cells per strain analyzed in three independent experiments. Range of the whiskers indicate population variability referring to the extent of dispersion around the median. Note that the measurements were performed using the same cultures from which the exponential-phase data are shown in Fig. 9.

**Supplementary Fig. S15:**
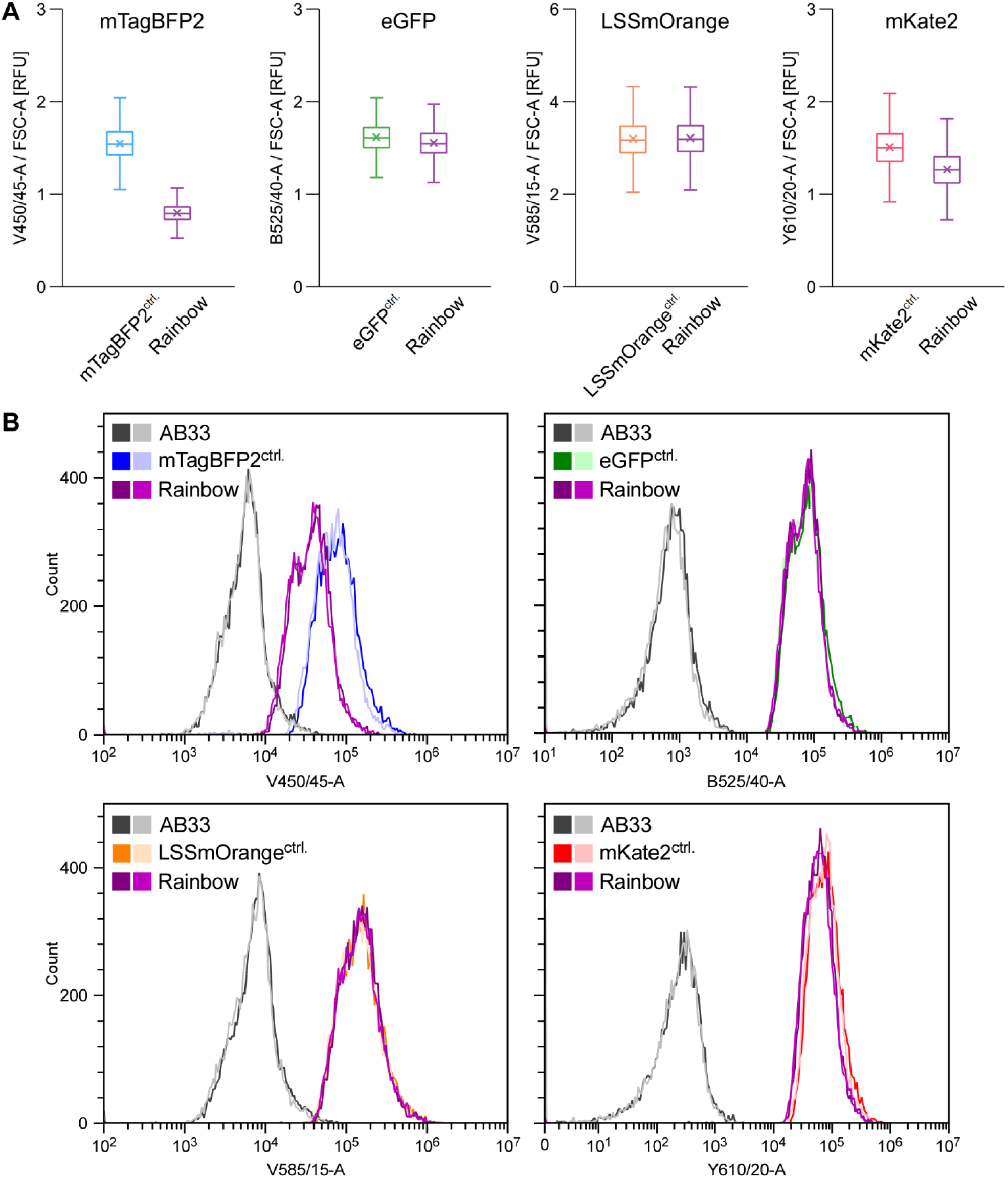
Comparison of the rainbow strain harboring four different fluorescent proteins with control (^ctrl.^) strains producing each fluorescent protein individually at the single-cell level. **A)** mTagBFP2, eGFP, LSSmOrange and mKate2 fluorescence intensity of each cell was determined using the V450/45-A, the B525/40-A, the V585/15-A or the Y610/20-A channel of the flow cytometer, respectively, and fluorescence values were normalized to forward-scatter area (FSC-A) to account for differences in cell size. Boxplots show the statistics including the arithmetic mean (x) of ∼ 30,000 cells per strain analyzed in three independent experiments. Range of the whiskers indicate population variability referring to the extent of dispersion around the median. **B)** Cytometer histograms showing the progenitor strain (AB33), the control strains (mTagBFP2^ctrl.^, eGFP^ctrl.^, LSSmOrange^ctrl.^ and mKate2^ctrl.^) and the rainbow strain in the four different channels. Two biological replicates are shown as indicated by the darker and lighter shade of the same color, respectively.

**Supplementary Fig. S16:**
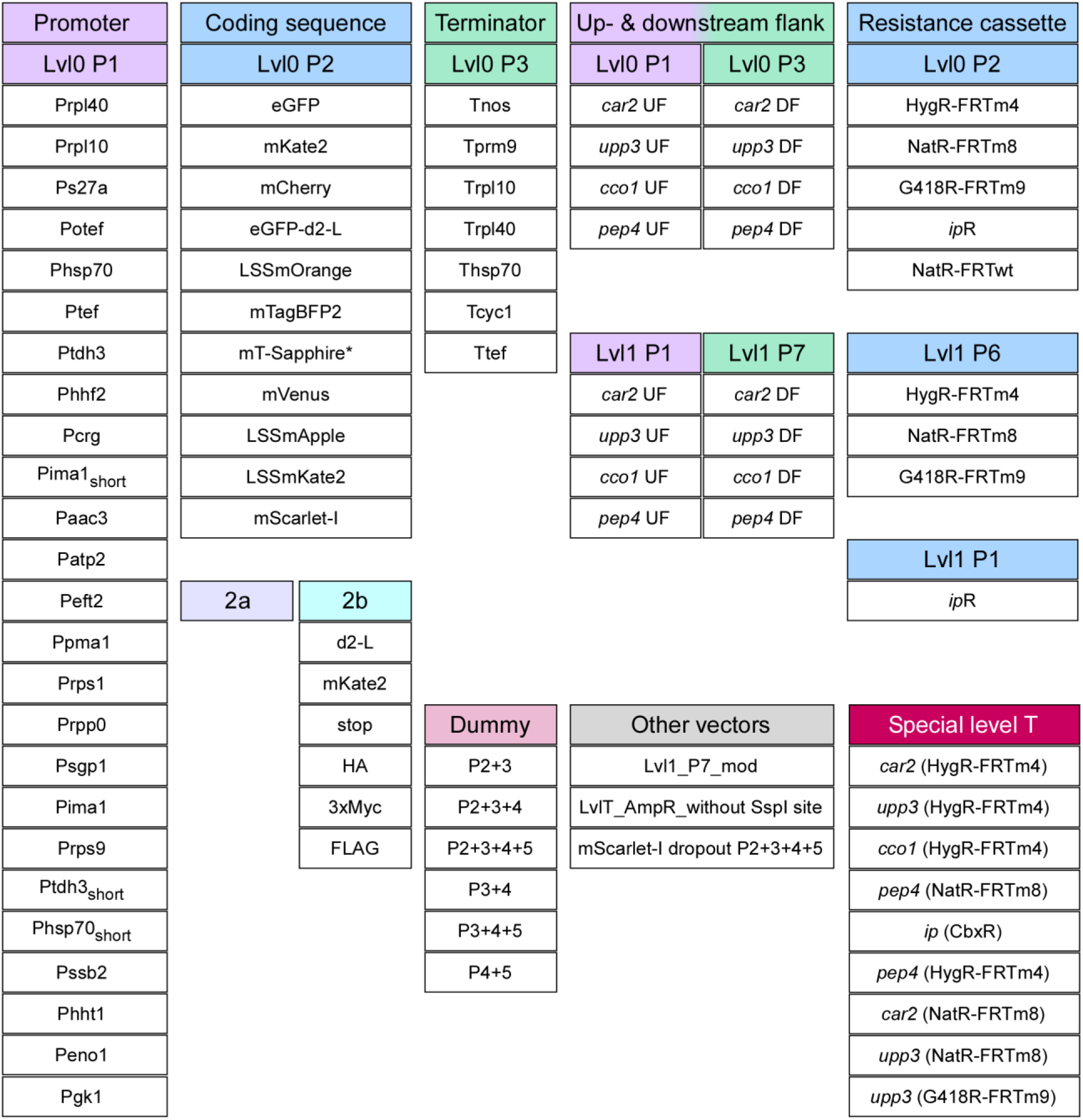
Overview of the UstiGate plasmid collection with 92 basic parts and vectors (Addgene kit deposit in progress). The toolkit includes more than 20 different promoters, a selection of terminators, several fluorescent proteins and protein tags. Up- (UF) and downstream flanks (DF) of genomic insertion loci as well as resistance cassettes are available as level 0 (Lvl0) and level 1 (Lvl1) parts at their respective position (P). The kit also comprises special level T plasmids for several genomic loci and additional dummy vectors.

**Supplementary Table S1:**
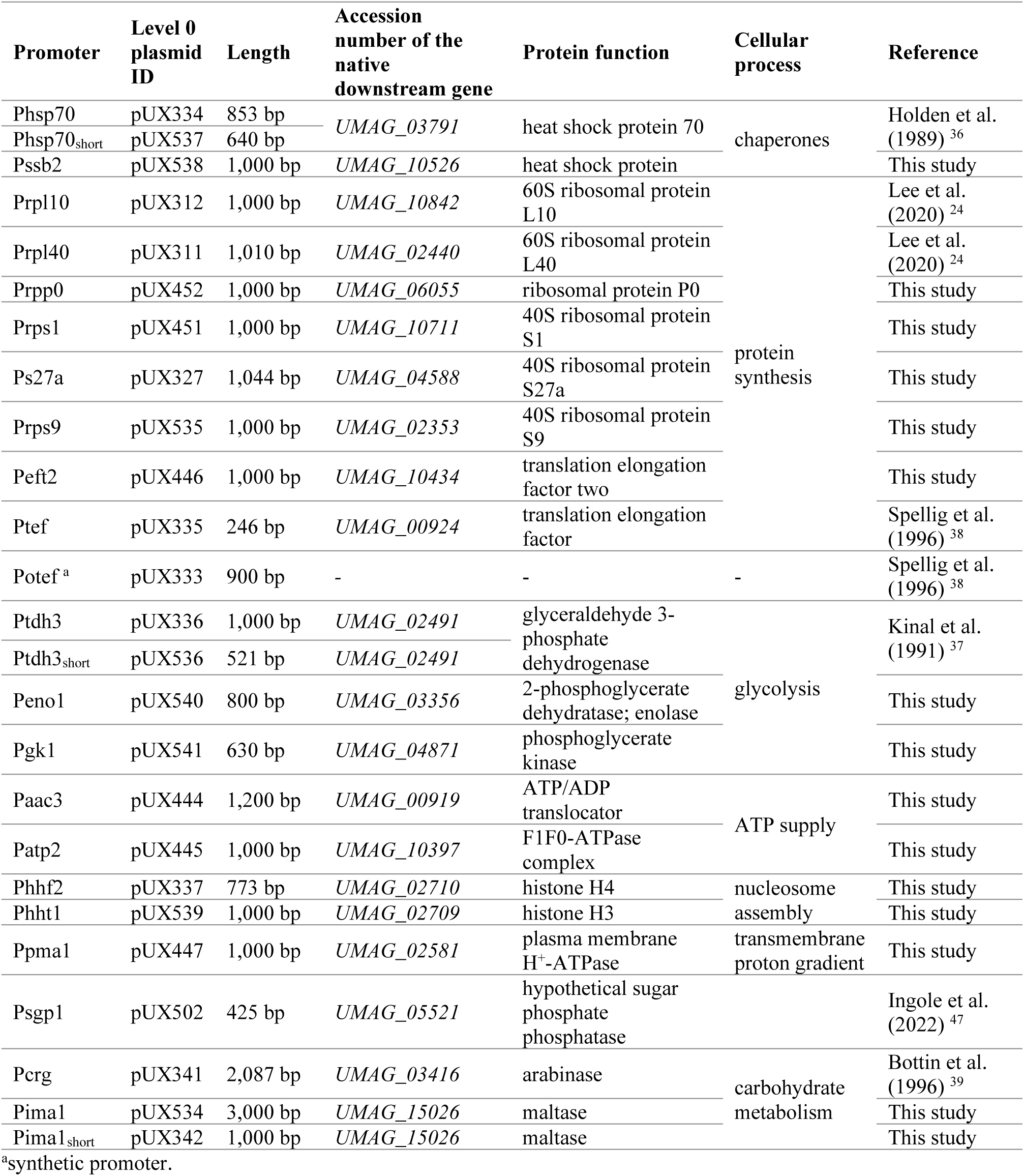
Promoters characterized in this study.

**Supplementary Table S2:** Fluorescent proteins utilized in this study.

| Fluorescent protein | FPbase ID | Max. Ex [nm]<sup>a</sup> | Max. Em [nm]<sup>a</sup> | Ex [nm]<sup>b</sup> | Em [nm]<sup>b</sup> | Reference |
| --- | --- | --- | --- | --- | --- | --- |
| Electra2 | 5GJX3 | 403 | 454 | 400/9 | 450/20 | Papadaki et al. (2022)<sup>85</sup> |
| mTagBFP2 | ZO7NN | 399 | 454 | 400/9 | 450/20 | Subach et al. (2011)<sup>95</sup> |
| mTurquoise2 | 7AV5G | 434 | 474 | 435/9 | 475/20 | Goedhart et al. (2012)<sup>86</sup> |
| eGFP | R9NL8 | 488 | 507 | 480/9 | 515/20 | Cormack et al. (1996)<sup>78</sup> |
| mT-Sapphire*<sup>c</sup> | KSK4V | 399 | 511 | 400/9 | 510/20 | Ai et al. (2008)<sup>93</sup> |
| mVenus | WCSN6 | 515 | 527 | 505/9 | 540/20 | Kremers et al. (2006)<sup>87</sup> |
| LSSmOrange | OVJRJ | 437 | 572 | 435/9 | 570/20 | Shcherbakova et al. (2012)<sup>94</sup> |
| mScarlet-I | 6VVTK | 569 | 593 | 565/9 | 600/20 | Bindels et al. (2017)<sup>97</sup> |
| LSSmApple | JMXXU | 490 | 600 | 490/9 | 600/20 | Ejike et al. (2024)<sup>96</sup> |
| LSSmKate2 | RU9EF | 460 | 605 | 460/9 | 605/20 | Piatkevich et al. (2010)<sup>98</sup> |
| mCherry | ZERB6 | 587 | 610 | 580/9 | 615/20 | Shaner et al. (2004)<sup>88</sup> |
| mKate2 | DBBO8 | 588 | 633 | 590/9 | 635/20 | Shcherbo et al. (2009)<sup>89</sup> |

<sup>a</sup>excitation (Ex) and emission (Em) maxima of the fluorescent protein.
<sup>b</sup>excitation (Ex) and emission (Em) wavelength/bandwidth utilized for plate reader-based fluorescence measurements.
<sup>c</sup>mT-Sapphire\*: mT-Sapphire with the following mutations: N147T S148A L232H.

**Supplementary Table S3:** Terminators characterized in this study.

| Terminator | Level 0 plasmid ID | Length | Accession number of the native upstream gene | Protein function | Cellular process | Reference |
| --- | --- | --- | --- | --- | --- | --- |
| Thsp70 | pUX542 | 500 bp | UMAG_03791 | heat shock protein 70 | chaperone | Brachmann et al. (2004)<sup>28</sup> |
| Ttef | pUX544 | 300 bp | UMAG_00924 | translation elongation factor | protein synthesis | This study |
| Trpl10 | pUX382 | 500 bp | UMAG_10842 | 60S ribosomal protein L10 |  | This study |
| Trpl40 | pUX383 | 400 bp | UMAG_02440 | 60S ribosomal protein L40 |  | This study |
| Tnos | pUX313 | 296 bp | - | nopaline synthase |  | Spellig et al. (1996)<sup>38</sup> |
| Tprm9 | pUX357 | 250 bp | YAR031W | pheromone regulated membrane protein |  | Curran et al. (2013)<sup>110</sup><br>Ingole et al. (2022)<sup>47</sup> |
| Tcyc1 | pUX543 | 280 bp | YJR048W | cytochrome c | cellular respiration | Brachmann et al. (2004)<sup>28</sup> |

**Supplementary Table S4:** Plasmids used or generated in this study.

| Plasmid | Resistance | Internal plasmid ID | Addgene kit | Reference |
| --- | --- | --- | --- | --- |
| pICH41295_Lvl0_P1 | SpecR | pUX285 | #1000000044 | Weber et al. (2011)<sup>55</sup> |
| pICH41308_Lvl0_P2 | SpecR | pUX286 | #1000000044 | Weber et al. (2011)<sup>55</sup> |
| pICH41276_Lvl0_P3 | SpecR | pUX287 | #1000000044 | Weber et al. (2011)<sup>55</sup> |
| pICH41258_Lvl0_P2a | SpecR | pUX347 | #1000000044 | Weber et al. (2011)<sup>55</sup> |
| pICH41264_Lvl0_P2b | SpecR | pUX348 | #1000000044 | Weber et al. (2011)<sup>55</sup> |
| pICH47732_Lvl1_P1 | AmpR | pUX288 | #1000000044 | Weber et al. (2011)<sup>55</sup> |
| pICH47742_Lvl1_P2 | AmpR | pUX289 | #1000000044 | Weber et al. (2011)<sup>55</sup> |
| pICH47751_Lvl1_P3 | AmpR | pUX290 | #1000000044 | Weber et al. (2011)<sup>55</sup> |
| pICH47761_Lvl1_P4 | AmpR | pUX291 | #1000000044 | Weber et al. (2011)<sup>55</sup> |
| pICH47772_Lvl1_P5 | AmpR | pUX292 | #1000000044 | Weber et al. (2011)<sup>55</sup> |
| pICH47781_Lvl1_P6 | AmpR | pUX293 | #1000000044 | Weber et al. (2011)<sup>55</sup> |
| pICH47791_Lvl1_P7 | AmpR | pUX294 | #1000000044 | Weber et al. (2011)<sup>55</sup> |
| Lvl1_P7_mod | AmpR | pUX349 | <sup>a</sup> | This study |
| pICH41722_linker_P1-7 | SpecR | pUX297 | #1000000044 | Weber et al. (2011)<sup>55</sup> |
| pICH41744_linker_P2-7 | SpecR | pUX298 | #1000000044 | Weber et al. (2011)<sup>55</sup> |
| pICH41766_linker_P3-7 | SpecR | pUX299 | #1000000044 | Weber et al. (2011)<sup>55</sup> |
| pICH41780_linker_P4-7 | SpecR | pUX300 | #1000000044 | Weber et al. (2011)<sup>55</sup> |
| pICH41800_linker_P5-7 | SpecR | pUX301 | #1000000044 | Weber et al. (2011)<sup>55</sup> |
| pICH41822_linker_P6-7 | SpecR | pUX302 | #1000000044 | Weber et al. (2011)<sup>55</sup> |
| pICH50866_linker_P7-7 | SpecR | pUX303 | #1000000044 | Weber et al. (2011)<sup>55</sup> |
| pICH54011_Dummy_P1 | AmpR | pUX304 | #1000000044 | Weber et al. (2011)<sup>55</sup> |
| pICH54022_Dummy_P2 | AmpR | pUX305 | #1000000044 | Weber et al. (2011)<sup>55</sup> |
| pICH54033_Dummy_P3 | AmpR | pUX306 | #1000000044 | Weber et al. (2011)<sup>55</sup> |
| pICH54044_Dummy_P4 | AmpR | pUX307 | #1000000044 | Weber et al. (2011)<sup>55</sup> |
| pICH54055_Dummy_P5 | AmpR | pUX308 | #1000000044 | Weber et al. (2011)<sup>55</sup> |
| pICH54066_Dummy_P6 | AmpR | pUX309 | #1000000044 | Weber et al. (2011)<sup>55</sup> |
| pICH54077_Dummy_P7 | AmpR | pUX310 | #1000000044 | Weber et al. (2011)<sup>55</sup> |
| Dummy_P2+3 | AmpR | pUX683 | <sup>a</sup> | This study |
| Dummy_P2+3+4 | AmpR | pUX684 | <sup>a</sup> | This study |
| Dummy_P2+3+4+5 | AmpR | pUX685 | <sup>a</sup> | This study |
| Dummy_P3+4 | AmpR | pUX686 | <sup>a</sup> | This study |
| Dummy_P3+4+5 | AmpR | pUX687 | <sup>a</sup> | This study |
| Dummy_P4+5 | AmpR | pUX688 | <sup>a</sup> | This study |
| pCAT.334_LvlT_SpecR | SpecR | pUX295 | #1000000146 | Vasudevan et al. (2019)<sup>68</sup> |
| pCAT.015_LvlT_AmpR | AmpR | pUX296 | #1000000146 | Vasudevan et al. (2019)<sup>68</sup> |
| LvlT_AmpR_without SspI site | AmpR | pUX453 | <sup>a</sup> | This study |
| pC0.116_unmark linker | SpecR | pUX317 | #1000000146 | Vasudevan et al. (2019)<sup>68</sup> |
| pC0.117_up flank linker | SpecR | pUX318 | #1000000146 | Vasudevan et al. (2019)<sup>68</sup> |
| pC0.118_down flank linker | SpecR | pUX319 | #1000000146 | Vasudevan et al. (2019)<sup>68</sup> |
| pC0.119_sacB up linker | SpecR | pUX320 | #1000000146 | Vasudevan et al. (2019)<sup>68</sup> |
| pC0.120_AbR down linker | SpecR | pUX321 | #1000000146 | Vasudevan et al. (2019)<sup>68</sup> |
| pC0.121_AbR up linker | SpecR | pUX322 | #1000000146 | Vasudevan et al. (2019)<sup>68</sup> |
| Lvl0_P1_Prpl40 | SpecR | pUX311 | <sup>a</sup> | This study |
| Lvl0_P1_Prpl10 | SpecR | pUX312 | <sup>a</sup> | This study |
| Lvl0_P1_Ps27a | SpecR | pUX327 | <sup>a</sup> | This study |
| Lvl0_P1_Potef | SpecR | pUX333 | <sup>a</sup> | This study |
| Lvl0_P1_Phsp70 | SpecR | pUX334 | <sup>a</sup> | This study |
| Lvl0_P1_Ptef | SpecR | pUX335 | <sup>a</sup> | This study |
| Lvl0_P1_Ptdh3 | SpecR | pUX336 | <sup>a</sup> | This study |
| Lvl0_P1_Phhf2 | SpecR | pUX337 | <sup>a</sup> | This study |
| Lvl0_P1_Pcrg | SpecR | pUX341 | <sup>a</sup> | This study |
| Lvl0_P1_Pima1<sub>short</sub> | SpecR | pUX342 | <sup>a</sup> | This study |
| Lvl0_P1_Paac3 | SpecR | pUX444 | <sup>a</sup> | This study |
| Lvl0_P1_Patp2 | SpecR | pUX445 | <sup>a</sup> | This study |
| Lvl0_P1_Peft2 | SpecR | pUX446 | <sup>a</sup> | This study |
| Lvl0_P1_Ppma1 | SpecR | pUX447 | <sup>a</sup> | This study |
| Lvl0_P1_Prps1 | SpecR | pUX451 | <sup>a</sup> | This study |
| Lvl0_P1_Prpp0 | SpecR | pUX452 | <sup>a</sup> | This study |
| Lvl0_P1_Psgp1 | SpecR | pUX502 | <sup>a</sup> | This study |
| Lvl0_P1_Pima1 | SpecR | pUX534 | <sup>a</sup> | This study |
| Lvl0_P1_Prps9 | SpecR | pUX535 | <sup>a</sup> | This study |
| Lvl0_P1_Ptdh3<sub>short</sub> | SpecR | pUX536 | <sup>a</sup> | This study |
| Lvl0_P1_Phsp70<sub>short</sub> | SpecR | pUX537 | <sup>a</sup> | This study |
| Lvl0_P1_Pssb2 | SpecR | pUX538 | <sup>a</sup> | This study |
| Lvl0_P1_Phht1 | SpecR | pUX539 | <sup>a</sup> | This study |
| Lvl0_P1_Peno1 | SpecR | pUX540 | <sup>a</sup> | This study |
| Lvl0_P1_Pgk1 | SpecR | pUX541 | <sup>a</sup> | This study |
| Lvl0_P2_eGFP | SpecR | pUX314 | <sup>a</sup> | This study |
| Lvl0_P2_eGFP-d2-L | SpecR | pUX390 | <sup>a</sup> | This study |
| Lvl0_P2_mKate2 | SpecR | pUX315 | <sup>a</sup> | This study |
| Lvl0_P2_mCherry | SpecR | pUX358 | <sup>a</sup> | This study |
| Lvl0_P2_LSSmOrange | SpecR | pUX668 | <sup>a</sup> | This study |
| Lvl0_P2_Electra2 | SpecR | pUX669 |  | This study |
| Lvl0_P2_mTagBFP2 | SpecR | pUX670 | <sup>a</sup> | This study |
| Lvl0_P2_mT-Sapphire* | SpecR | pUX698 | <sup>a</sup> | This study |
| Lvl0_P2_mVenus | SpecR | pUX788 | <sup>a</sup> | This study |
| Lvl0_P2_LSSmApple | SpecR | pUX789 | <sup>a</sup> | This study |
| Lvl0_P2_LSSmKate2 | SpecR | pUX790 | <sup>a</sup> | This study |
| Lvl0_P2_mScarlet-I | SpecR | pUX791 | <sup>a</sup> | This study |
| Lvl0_P2_mTurquoise2 | SpecR | pUX792 |  | This study |
| Lvl0_P2b_d2-L_stop | SpecR | pUX389 | <sup>a</sup> | This study |
| Lvl0_P2b_RS linker_mKate2 | SpecR | pUX484 | <sup>a</sup> | This study |
| Lvl0_P2b_stop | SpecR | pUX554 | <sup>a</sup> | This study |
| Lvl0_P2b_HA_stop | SpecR | pUX555 | <sup>a</sup> | This study |
| Lvl0_P2b_3xMyc_stop | SpecR | pUX556 | <sup>a</sup> | This study |
| Lvl0_P2b_FLAG_stop | SpecR | pUX601 | <sup>a</sup> | This study |
| Lvl0_P3_Tnos | SpecR | pUX313 | <sup>a</sup> | This study |
| Lvl0_P3_Tprm9 | SpecR | pUX357 | <sup>a</sup> | This study |
| Lvl0_P3_Trpl10 | SpecR | pUX382 | <sup>a</sup> | This study |
| Lvl0_P3_Trpl40 | SpecR | pUX383 | <sup>a</sup> | This study |
| Lvl0_P3_Thsp70 | SpecR | pUX542 | <sup>a</sup> | This study |
| Lvl0_P3_Tcyc1 | SpecR | pUX543 | <sup>a</sup> | This study |
| Lvl0_P3_Ttef | SpecR | pUX544 | <sup>a</sup> | This study |
| Lvl0_P1_car2_UF | SpecR | pUX328 | <sup>a</sup> | This study |
| Lvl0_P3_car2_DF | SpecR | pUX329 | <sup>a</sup> | This study |
| Lvl0_P1_upp3_UF | SpecR | pUX330 | <sup>a</sup> | This study |
| Lvl0_P3_upp3_DF | SpecR | pUX331 | <sup>a</sup> | This study |
| Lvl0_P1_cco1_UF | SpecR | pUX384 | <sup>a</sup> | This study |
| Lvl0_P3_cco1_DF | SpecR | pUX385 | <sup>a</sup> | This study |
| Lvl0_P1_pep4_UF | SpecR | pUX408 | <sup>a</sup> | This study |
| Lvl0_P3_pep4_DF | SpecR | pUX409 | <sup>a</sup> | This study |
| Lvl0_P2_HygR-FRTm4 | SpecR | pUX332 | <sup>a</sup> | This study |
| Lvl0_P2_NatR-FRTm8 | SpecR | pUX407 | <sup>a</sup> | This study |
| Lvl0_P2_G418R-FRTm9 | SpecR | pUX428 | <sup>a</sup> | This study |
| Lvl0_P1_ipR | SpecR | pUX454 | <sup>a</sup> | This study |
| Lvl0_P2_NatR-FRTwt | SpecR | pUX475 | <sup>a</sup> | This study |
| Lvl1_P1_car2_UF | AmpR | pUX351 | <sup>a</sup> | This study |
| Lvl1_P7_car2_DF | AmpR | pUX352 | <sup>a</sup> | This study |
| Lvl1_P1_upp3_UF | AmpR | pUX353 | <sup>a</sup> | This study |
| Lvl1_P7_upp3_UF | AmpR | pUX354 | <sup>a</sup> | This study |
| Lvl1_P1_pep4_UF | AmpR | pUX416 | <sup>a</sup> | This study |
| Lvl1_P7_pep4_DF | AmpR | pUX417 | <sup>a</sup> | This study |
| Lvl1_P1_cco1_UF | AmpR | pUX419 | <sup>a</sup> | This study |
| Lvl1_P7_cco1_DF | AmpR | pUX420 | <sup>a</sup> | This study |
| Lvl1_P6_HygR-FRTm4 | AmpR | pUX355 | <sup>a</sup> | This study |
| Lvl1_P6_NatR-FRTm8 | AmpR | pUX418 | <sup>a</sup> | This study |
| Lvl1_P1_ipR | AmpR | pUX455 | <sup>a</sup> | This study |
| Lvl1_P6_G418R-FRTm9 | AmpR | pUL204 | <sup>a</sup> | This study |
| mScarlet-I dropout_P2+3+4+5 | AmpR | pUX909 | <sup>a</sup> | This study |
| Special LvlT_<br>car2_HygR-FRTm4 | AmpR | pUX361 | <sup>a</sup> | This study |
| Special LvlT_<br>car2_NatR-FRTm8 | AmpR | pUX715 | <sup>a</sup> | This study |
| Special LvlT_<br>upp3_HygR-FRTm4 | AmpR | pUX362 | <sup>a</sup> | This study |
| Special LvlT_<br>cco1_HygR-FRTm4 | AmpR | pUX421 | <sup>a</sup> | This study |
| Special LvlT_<br>pep4_HygR-FRTm4 | AmpR | pUX493 | <sup>a</sup> | This study |
| Special LvlT_<br>pep4_NatR-FRTm8 | AmpR | pUX425 | <sup>a</sup> | This study |
| Special LvlT_ipS_CbxR | AmpR | pUX456 | <sup>a</sup> | This study |
| Special LvlT_<br>upp3_NatR-FRTm8 | AmpR | pIL263 | <sup>a</sup> | This study |
| Special LvlT_<br>upp3_G418R-FRTm8 | AmpR | pUL676 | <sup>a</sup> | This study |
| Potef:eGFP_Tnos<br>(car2, HygR-FRTm4) | AmpR | pUX363 |  | This study |
| Pcrg:eGFP_Tnos<br>(car2, HygR-FRTm4) | AmpR | pUX365 |  | This study |
| Prpl10:eGFP_Tnos<br>(car2, HygR-FRTm4) | AmpR | pUX368 |  | This study |
| Prpl40:eGFP_Tnos<br>(car2, HygR-FRTm4) | AmpR | pUX369 |  | This study |
| Ps27a:eGFP_Tnos<br>(car2, HygR-FRTm4) | AmpR | pUX370 |  | This study |
| Phsp70:eGFP_Tnos<br>(car2, HygR-FRTm4) | AmpR | pUX371 |  | This study |
| Ptef:eGFP_Tnos<br>(car2, HygR-FRTm4) | AmpR | pUX372 |  | This study |
| Ptdh3:eGFP_Tnos<br>(car2, HygR-FRTm4) | AmpR | pUX373 |  | This study |
| Phhf2:eGFP_Tnos<br>(car2, HygR-FRTm4) | AmpR | pUX374 |  | This study |
| Pima1<sub>short</sub>:eGFP_Tnos<br>(car2, HygR-FRTm4) | AmpR | pUX375 |  | This study |
| Paac3:eGFP_Tnos<br>(car2, HygR-FRTm4) | AmpR | pUX459 |  | This study |
| Patp2:eGFP_Tnos<br>(car2, HygR-FRTm4) | AmpR | pUX460 |  | This study |
| Peft2:eGFP_Tnos<br>(car2, HygR-FRTm4) | AmpR | pUX461 |  | This study |
| Ppma1:eGFP_Tnos<br>(car2, HygR-FRTm4) | AmpR | pUX462 |  | This study |
| Prps1:eGFP_Tnos<br>(car2, HygR-FRTm4) | AmpR | pUX466 |  | This study |
| Prpp0:eGFP_Tnos<br>(car2, HygR-FRTm4) | AmpR | pUX467 |  | This study |
| Pcrg:eGFP-d2-L_Tnos<br>(car2, HygR-FRTm4) | AmpR | pUX495 |  | This study |
| Prpl10:eGFP_Trpl10 (car2, HygR-FRTm4) | AmpR | pUX497 |  | This study |
| Prpl10:eGFP_Trpl40 (car2, HygR-FRTm4) | AmpR | pUX498 |  | This study |
| Prpl10:eGFP_Tprm9 (car2, HygR-FRTm4) | AmpR | pUX499 |  | This study |
| Phhf2:eGFP-d2-L_Tnos (car2, HygR-FRTm4) | AmpR | pUX514 |  | This study |
| Phsp70:eGFP-d2-L_Tnos (car2, HygR-FRTm4) | AmpR | pUX515 |  | This study |
| Prpl10:eGFP-d2-L_Tnos (car2, HygR-FRTm4) | AmpR | pUX516 |  | This study |
| Potef:eGFP-d2-L_Tnos (car2, HygR-FRTm4) | AmpR | pUX517 |  | This study |
| Paac3:eGFP-d2-L_Tnos (car2, HygR-FRTm4) | AmpR | pUX518 |  | This study |
| Ptdh3:eGFP-d2-L_Tnos (car2, HygR-FRTm4) | AmpR | pUX519 |  | This study |
| Ppma1:eGFP-d2-L_Tnos (car2, HygR-FRTm4) | AmpR | pUX520 |  | This study |
| Ptef:eGFP-d2-L_Tnos (car2, HygR-FRTm4) | AmpR | pUX521 |  | This study |
| Peft2:eGFP-d2-L_Tnos (car2, HygR-FRTm4) | AmpR | pUX522 |  | This study |
| Patp2:eGFP-d2-L_Tnos (car2, HygR-FRTm4) | AmpR | pUX523 |  | This study |
| Prpl40:eGFP-d2-L_Tnos (car2, HygR-FRTm4) | AmpR | pUX524 |  | This study |
| Psgp1:eGFP_Tnos (car2, HygR-FRTm4) | AmpR | pUX526 |  | This study |
| Pima1:eGFP_Tnos (car2, HygR-FRTm4) | AmpR | pUX570 |  | This study |
| Prps9:eGFP_Tnos (car2, HygR-FRTm4) | AmpR | pUX572 |  | This study |
| Ptdh3<sub>short</sub>:eGFP_Tnos (car2, HygR-FRTm4) | AmpR | pUX573 |  | This study |
| Phsp70<sub>short</sub>:eGFP_Tnos (car2, HygR-FRTm4) | AmpR | pUX574 |  | This study |
| Pssb2:eGFP_Tnos (car2, HygR-FRTm4) | AmpR | pUX575 |  | This study |
| Phht1:eGFP_Tnos (car2, HygR-FRTm4) | AmpR | pUX576 |  | This study |
| Peno1:eGFP_Tnos (car2, HygR-FRTm4) | AmpR | pUX577 |  | This study |
| Pgk1:eGFP_Tnos (car2, HygR-FRTm4) | AmpR | pUX578 |  | This study |
| Prpl10:eGFP_Thsp70 (car2, HygR-FRTm4) | AmpR | pUX579 |  | This study |
| Prpl10:eGFP_Tcyc1 (car2, HygR-FRTm4) | AmpR | pUX580 |  | This study |
| Prpl10:eGFP_Ttef (car2, HygR-FRTm4) | AmpR | pUX581 |  | This study |
| Paac3:mKate2_Tnos (car2, HygR-FRTm4) | AmpR | pUX610 |  | This study |
| Prpl10:mKate2_Tnos (car2, HygR-FRTm4) | AmpR | pUX611 |  | This study |
| Potef:mKate2_Tnos (car2, HygR-FRTm4) | AmpR | pUX612 |  | This study |
| Phsp70<sub>short</sub>:eGFP_Tnos (cco1, HygR-FRTm4) | AmpR | pUX613 |  | This study |
| Paac3:eGFP_Tnos (cco1, HygR-FRTm4) | AmpR | pUX614 |  | This study |
| Prpl10:eGFP_Tnos (cco1, HygR-FRTm4) | AmpR | pUX615 |  | This study |
| Potef:eGFP_Tnos (cco1, HygR-FRTm4) | AmpR | pUX616 |  | This study |
| Phsp70<sub>short</sub>:eGFP_Tnos (upp3, HygR-FRTm4) | AmpR | pUX617 |  | This study |
| Paac3:eGFP_Tnos (upp3, HygR-FRTm4) | AmpR | pUX618 |  | This study |
| Prpl10:eGFP_Tnos (upp3, HygR-FRTm4) | AmpR | pUX619 |  | This study |
| Potef:eGFP_Tnos (upp3, HygR-FRTm4) | AmpR | pUX620 |  | This study |
| Phsp70<sub>short</sub>:eGFP_Tnos (pep4, HygR-FRTm4) | AmpR | pUX621 |  | This study |
| Paac3:eGFP_Tnos (pep4, HygR-FRTm4) | AmpR | pUX622 |  | This study |
| Prpl10:eGFP_Tnos (pep4, HygR-FRTm4) | AmpR | pUX623 |  | This study |
| Potef:eGFP_Tnos (pep4, HygR-FRTm4) | AmpR | pUX624 |  | This study |
| Lvl1_P2_Prpl10:eGFP_Tnos | AmpR | pUX671 |  | This study |
| Lvl1_P3_Peno1:mKate2_Trpl10 | AmpR | pUX672 |  | This study |
| Lvl1_P4_Ptef:LSSmOrange_Trpl40 | AmpR | pUX673 |  | This study |
| Lvl1_P5_Paac3:mTagBFP2_Ttef | AmpR | pUX675 |  | This study |
| Prpl10:eGFP_Tnos (car2, HygR-FRTm4) | SpecR | pUX676 |  | This study |
| Peno1:mKate2_Trpl10 (car2, HygR-FRTm4) | SpecR | pUX677 |  | This study |
| Ptef:LSSmOrange_Trpl40 (car2, HygR-FRTm4) | SpecR | pUX678 |  | This study |
| Paac3:mTagBFP2_Ttef (car2, HygR-FRTm4) | SpecR | pUX680 |  | This study |
| Pima1:eGFP-d2-L_Tnos (car2, HygR-FRTm4) | AmpR | pUX682 |  | This study |
| Prpl10:eGFP_Tnos_Peno1:mKate2_Trpl10_Ptef:LSSmOrange_Trpl40_Paac3:mTagBFP2_Ttef (car2, HygR-FRTm4) | SpecR | pUX714 |  | This study |
| Phsp70<sub>short</sub>:eGFP_Tnos (car2, NatR-FRTm8) | AmpR | pUX716 |  | This study |
| Phsp70<sub>short</sub>:mKate2_Tnos (car2, NatR-FRTm8) | AmpR | pUX717 |  | This study |
| Phsp70<sub>short</sub>:mCherry_Tnos (car2, NatR-FRTm8) | AmpR | pUX782 |  | This study |
| Paac3:mCherry_Tnos (car2, HygR-FRTm4) | AmpR | pUX783 |  | This study |
| Prpl10:mCherry_Tnos (car2, HygR-FRTm4) | AmpR | pUX784 |  | This study |
| Potef:mCherry_Tnos (car2, HygR-FRTm4) | AmpR | pUX785 |  | This study |
| Prpl10:Electra2_Tnos (car2, HygR-FRTm4) | AmpR | pUX786 |  | This study |
| Prpl10:mT-Sapphire*_Tnos (car2, HygR-FRTm4) | AmpR | pUX787 |  | This study |
| Prpl10:LSSmOrange_Tnos (car2, HygR-FRTm4) | AmpR | pUX797 |  | This study |
| Prpl10:mTagBFP2_Tnos (car2, HygR-FRTm4) | AmpR | pUX798 |  | This study |
| Prpl10:mVenus_Tnos (car2, HygR-FRTm4) | AmpR | pUX799 |  | This study |
| Prpl10:LSSmApple_Tnos (car2, HygR-FRTm4) | AmpR | pUX800 |  | This study |
| Prpl10:LSSmKate2_Tnos (car2, HygR-FRTm4) | AmpR | pUX801 |  | This study |
| Prpl10:mScarlet-I_Tnos (car2, HygR-FRTm4) | AmpR | pUX802 |  | This study |
| Prpl10:mTurquoise2_Tnos (car2, HygR-FRTm4) | AmpR | pUX803 |  | This study |
| upp3Δ::NatR | AmpR | pUX323 |  | This study |
| Lvl1_P1_pep4Δ::NatR-FRTm8 | AmpR | pUX512 |  | This study |

<sup>a</sup>Addgene kit deposit is currently in progress.

**Supplementary Table S5:** Standard protocols for Golden Gate reactions.

| Protocol for Golden Gate reactions using BbsI<sup>a</sup> |  | Protocol for Golden Gate reactions using BsaI<sup>b</sup> |  |
| --- | --- | --- | --- |
| Lvl0 / LvlT acceptor vector (50 ng/μL) | 1 μL | Lvl1 / LvlT* acceptor vector (50 ng/μL) | 1 μL |
| Insert<sup>c</sup> / Lvl1 plasmid (50 ng/μL) | x μL / 1 μL | Lvl0 plasmid (50 ng/μL) | 1 μL |
| ... | ... | ... | ... |
| NEBridge® Ligase Master Mix | 5 μL | T4 DNA Ligase Buffer | 2 μL |
|  |  | T4 DNA Ligase | 1 μL |
| BbsI-HF | 1 μL | BsaI-HF | 1 μL |
| dH<sub>2</sub>O | ad 15 μL | dH<sub>2</sub>O | ad 20 μL |

<sup>a</sup>for the integration of genetic elements into a level 0 (Lvl0) acceptor vector or for the assembly of level 1 (Lvl1) parts into a level T (LvlT) acceptor vector.
<sup>b</sup>for the assembly of level 0 (Lvl0) parts into a level 1 (Lvl1) acceptor vector or a special level T plasmid (LvlT\*).
<sup>c</sup>purified PCR products were applied with a molar vector:insert ratio of 1:5.

**Supplementary Table S6:**
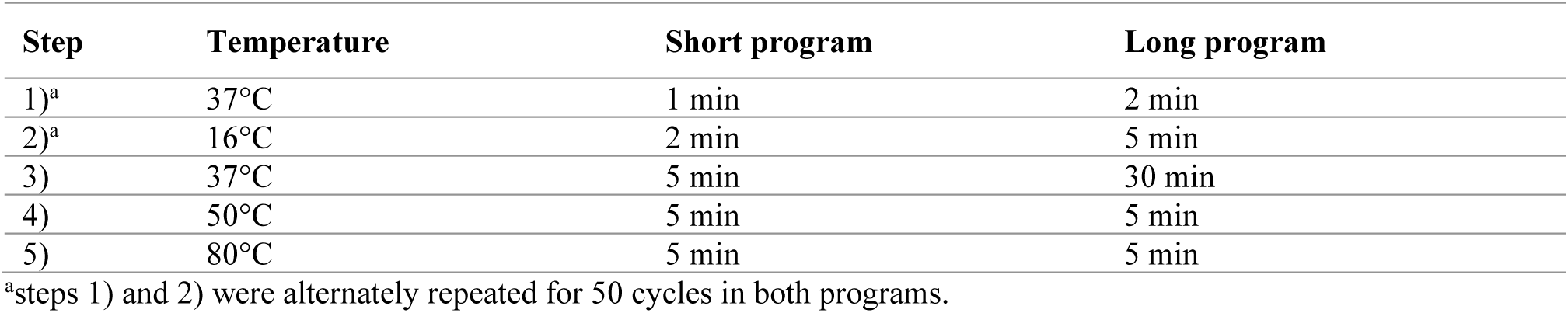
Standard short and long Golden Gate program.

**Supplementary Table S7:**
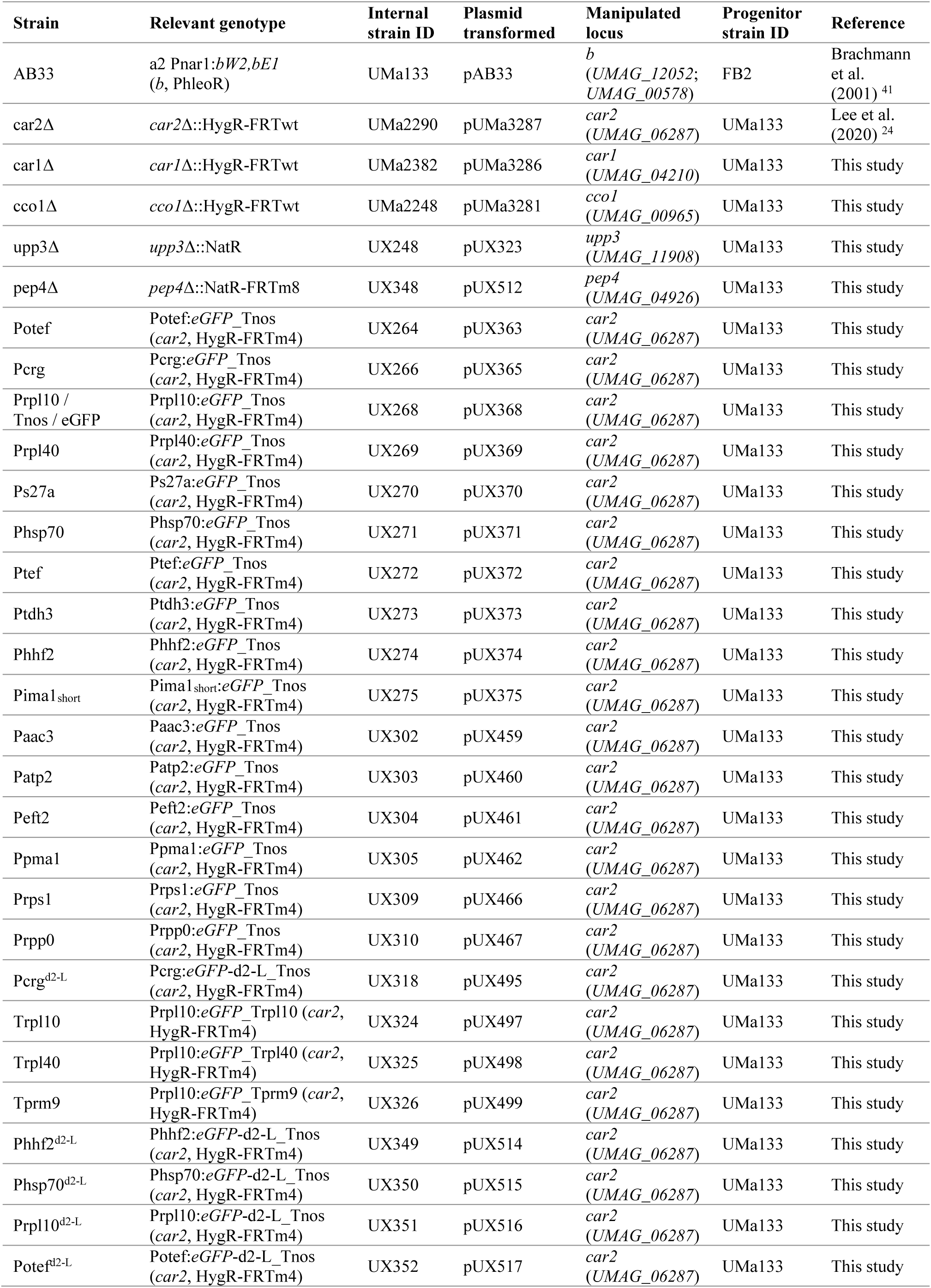
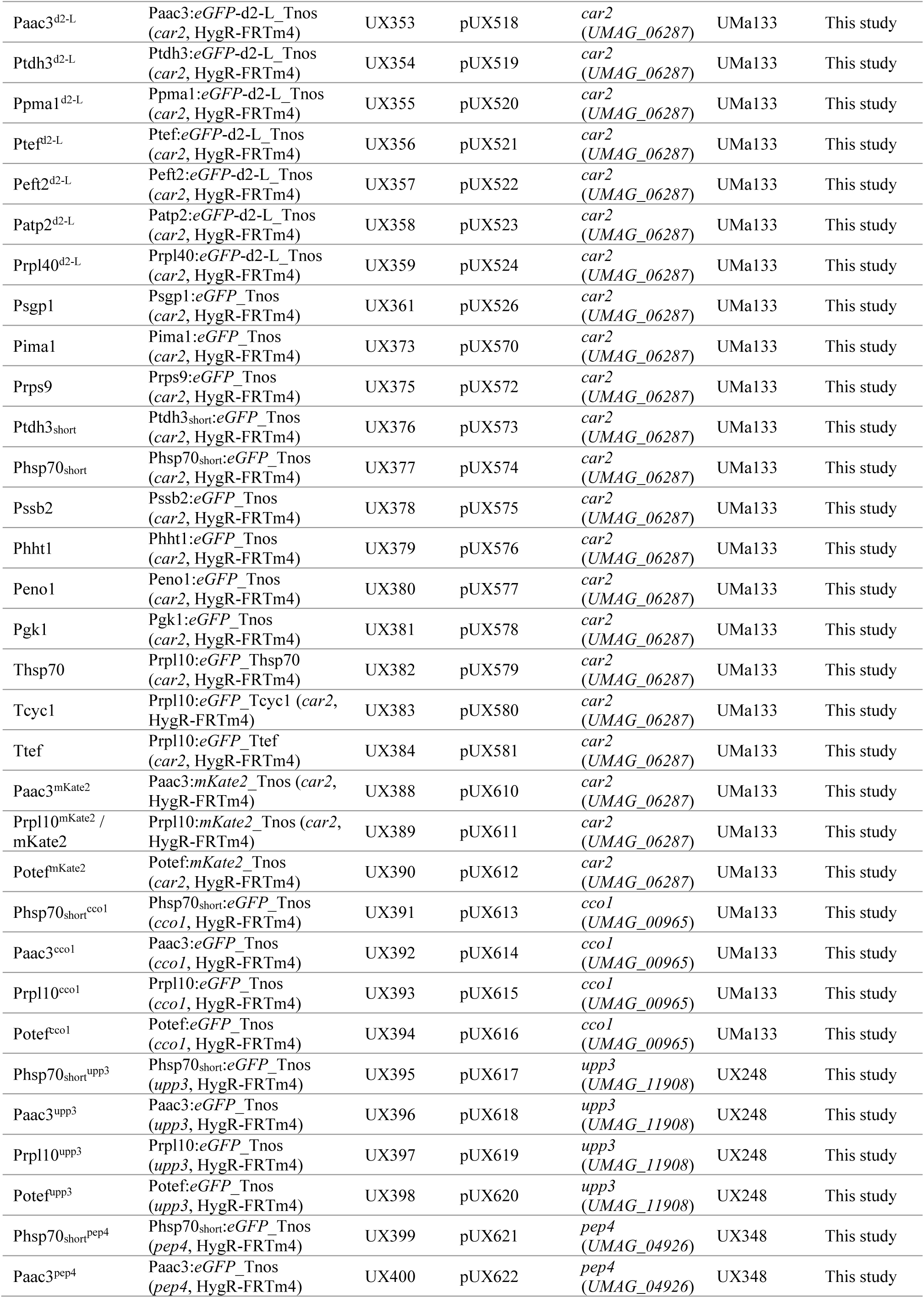
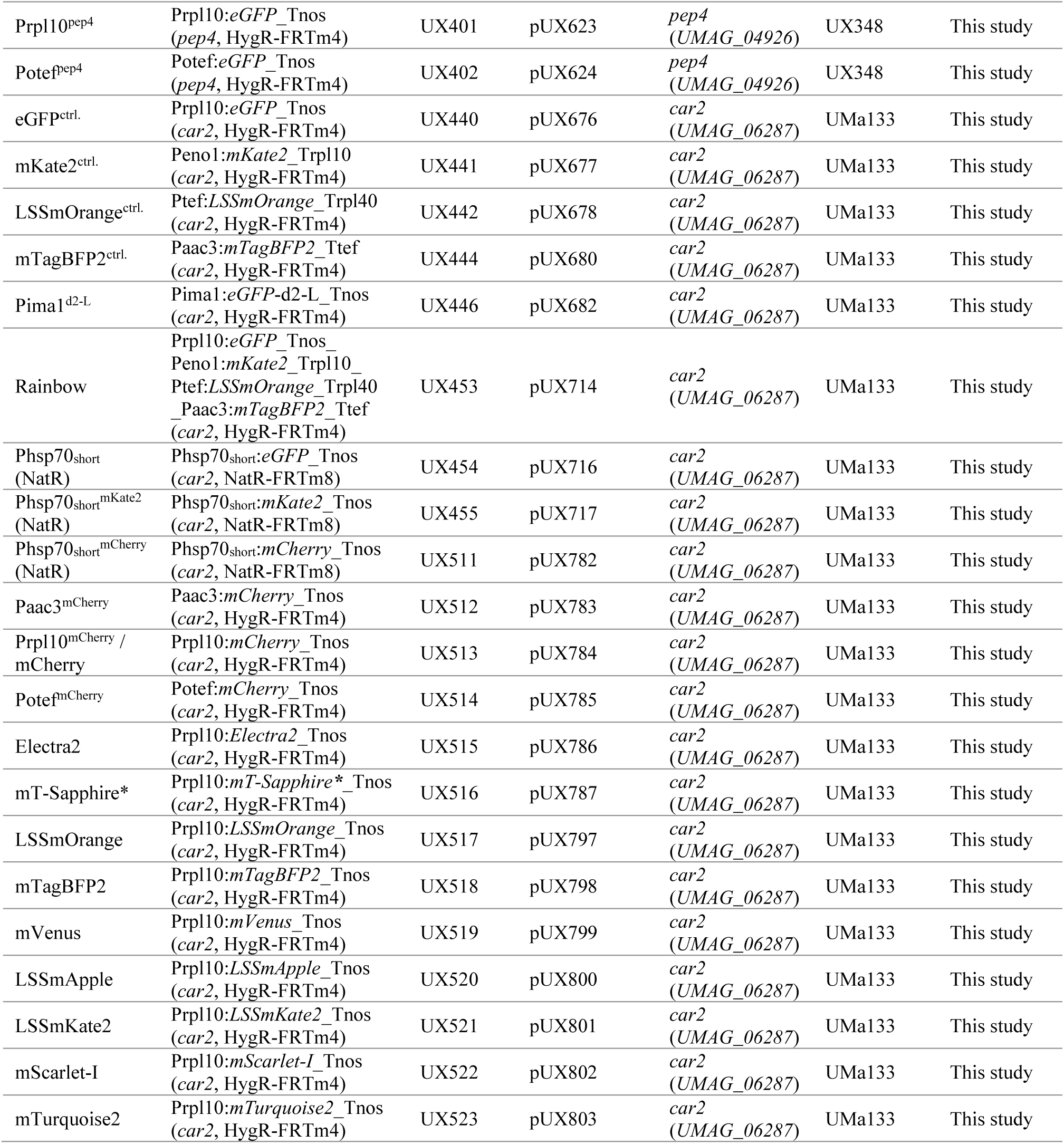
*U. maydis* strains used or generated in this study.

